# A high-level programming language for generative biology with Proto

**DOI:** 10.64898/2026.06.22.733870

**Authors:** Aditi T. Merchant, Daniel Guo, Ben Viggiano, Lucas Brennan-Almaraz, Evelyn Hur, Tina Mai, Peter Yin, Samuel H. King, Euan A. Ashley, Brian L. Hie

## Abstract

Programmable composition of complex systems is a longstanding goal of biological research. Generative modeling has improved the reliability of computational design, but existing methods are highly specialized and are difficult to extend or compose. Here, we introduce Proto, a high-level programming language for generative biology. By composing a small set of abstract primitives into structured programs, Proto encodes generative design campaigns across diverse modalities and scales—spanning DNA, RNA, proteins, ligands, and their interactions. Proto readily incorporates predictive models into generative workflows, which we leveraged to design alternatively spliced introns with experimental validation in human cell lines. Proto is natively multi-objective, enabling the design of promoter-repressor pairs with leading experimental success rates for synthetic protein-DNA design. Alongside AI agents, Proto enables the specification of complex pathways and regulatory logic through natural language instructions. We openly release Proto, including software infrastructure and user interfaces, to enable widespread access to generative biological programming.

## 1. Introduction

Biological engineering aims to design complex systems, from individual molecules to multicellular logic, with the same programmability as software or computer hardware (Basu et al., 2005; Cameron et al., 2014; Cane et al., 1998; Endy, 2005; Gardner et al., 2000; Gross et al., 1989; Huang et al., 2016; Jinek et al., 2012; Nielsen et al., 2016). However, despite decades of progress, this design process remains laborious and often heuristic. Researchers select and combine parts from nature, rely on intuition or large-scale screening, and iterate through costly build-test-learn cycles to navigate vast combinatorial design spaces (Arnold, 1998; Canton et al., 2008; Kosuri et al., 2013; Moon et al., 2012; Wang et al., 2009).

Advances in generative models have begun to improve the controllability and reliability of biological design (Hie and Yang, 2022; Listgarten and Jiang, 2026), but also face limitations that constrain the complexity of designed systems. Generative protein design has yielded *de novo* protein backbones, binders, and enzymes with highly divergent sequences and structures, but is largely limited to individual molecules and functions that require existing structural information (Dauparas et al., 2022; Pacesa et al., 2025; Watson et al., 2023). Generative genomics has enabled the design of multi-component and multi-modal systems, including complete bacteriophage genomes, but is constrained to the genetic architectures and synteny observed in nature (Brixi et al., 2026; Gosai et al., 2024; King et al., 2025; Merchant et al., 2025; Nguyen et al., 2024; Wang et al., 2026). Large language models achieve strong generalist capabilities in knowledge retrieval, multi-step reasoning, and tool use, including for biological questions, but are mostly bounded by existing human knowledge, beyond which they remain prone to hallucination (Brown et al., 2020; Ji et al., 2023; Schick et al., 2023; Wei et al., 2022).

Each of these approaches addresses a portion of the demands of generative design for complex systems—such as biophysical models, evolutionary priors, or flexible reasoning—but none alone provides a unified system for composing these capabilities across scales and modalities. Moreover, many of these systems remain difficult to use without domain expertise, and naively combining these diverse methodologies without a unifying abstraction would only compound the overall complexity. Just as high-level programming languages like Verilog and C enabled the design of large-scale integrated circuits and computer programs, we also require an intuitive and accessible language that enables the design of complex biological systems.

Here, we introduce Proto, a high-level programming language for generative biology (**Figure 1A**). Proto encodes design campaigns into structured programs that compose generative models of biological sequences, predictive models of sequence desirability, and optimization procedures that steer generation toward more desirable sequences. Traditional biological programming, by contrast, defines its constituent modules with literal biological sequences, such as natural proteins, guide RNAs, or regulatory DNA elements, which are composed into chimeric constructs (Arnold, 1998; Endy, 2005; Morsut et al., 2016; Shetty et al., 2008; Voigt et al., 2002; Wang and Doudna, 2023). However, recombining parts from nature using manual intuition, rule-based heuristics, or high-throughput screens can struggle to achieve complex synthetic functions or large-scale combinatorial complexity. A key conceptual contribution of this work is to therefore define modularity and compositionality at a functional or a semantic level, and to leverage generative modeling to bridge the gap between this higher level of abstraction and low-level biological sequences (**Figure 1B**) (Hie et al., 2022).

**Figure 1.**
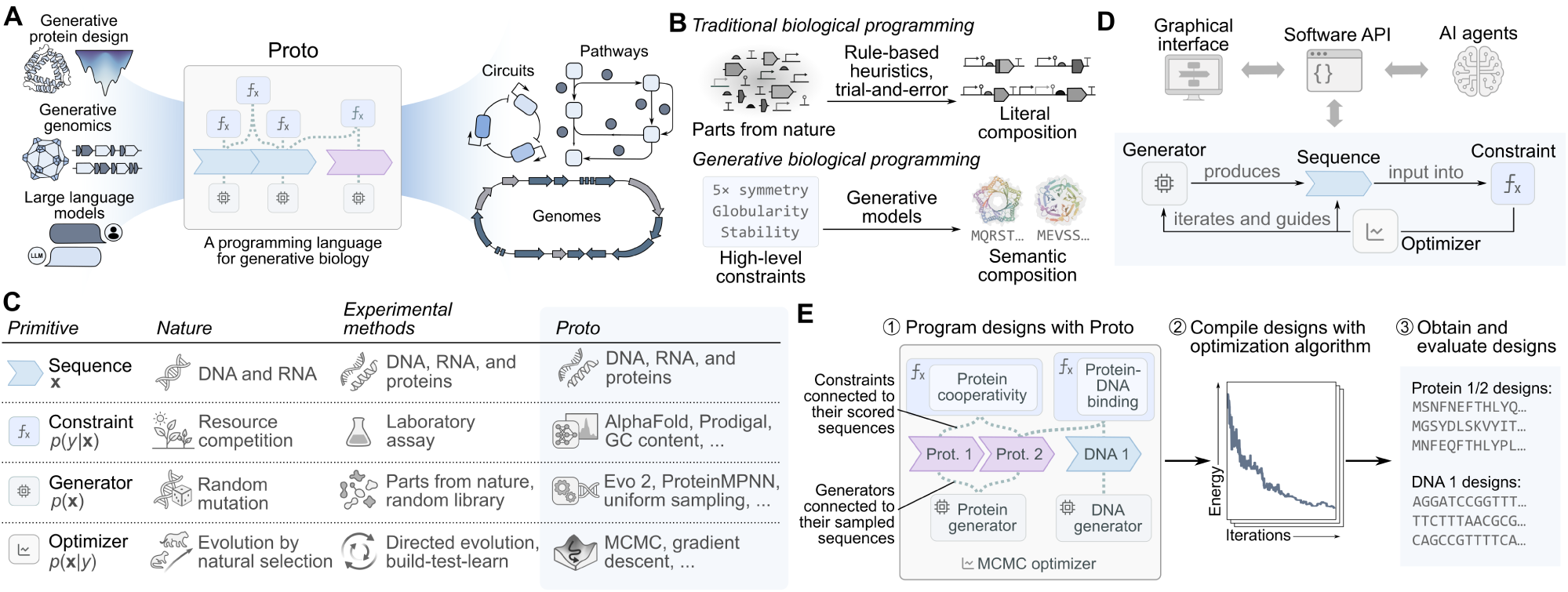
| Overview of Proto. (**A**) Proto composes generative and predictive models of DNA, RNA, proteins, ligands, and their interactions, and integrates with increasingly advanced LLMs and AI coding agents. Through this composition, Proto enables multi-objective, multi-modal, and multi-scale biological design. (**B**) Proto defines modularity and compositionality at a functional or semantic level and uses generative modeling to bridge this abstraction to low-level biological sequences, while preserving global functional coherence. In contrast, traditional biological programming composes literal sequence parts via intuition, heuristics, or trial-and-error, which can result in more brittle designs. (**C**) The four primitives of the Proto language—sequences, constraints, generators, and optimizers—together with their analogs in natural and experimental biological design and their formulation as factors of an energy-based model with target distribution *π*(*x*) ∝ *p*(*x*) exp(− *f* (*x*)/*T*), where *y* = *f* (*x*). (**D**) In Proto, generators propose candidate sequences, constraints score them, and optimizers steer generation toward lower-energy (more desirable) designs. Proto also contains higher-level interfaces that include a Python API (local library and hosted cloud version), a graphical user interface, and an agentic interface with general-purpose AI coding agents. (**E**) Proto’s overall workflow requires encoding a biological design task as a set of constraints and generators tied to sequences. An optimization algorithm then composes the generative models and compiles the full set of constraints into a unified energy function, producing a final set of designed sequences.

We first designed Proto for generality; rather than building around any particular model or modality, we built Proto around universal abstractions of design campaigns—sequences, constraints, generators, and optimizers—so that the language accommodates the rapid pace of development in biological AI (**Figure 1C**). We also designed Proto to be practically accessible; beyond providing a theoretical framework, Proto implements software infrastructure, a user interface, and compatibility with AI agents that enable usability across a broad range of researchers (**Figure 1D**). Finally, we designed Proto to be highly expressive; by composing simple primitives into modular programs, Proto has the capacity to encode systems at scales from individual molecules to multi-step pathways.

We first leveraged Proto to reimplement diverse design tasks from the existing literature, including symmetric protein homo-oligomers, *de novo* protein monomers, CRISPR-Cas systems, multi-kilobase chromatin accessibility patterns, and antibody complementarity-determining regions (CDRs). We next designed and experimentally validated novel systems that span all modalities of the central dogma (alternatively spliced RNA introns and pairs of DNA promoters and protein repressors) in eukaryotic and prokaryotic contexts. We achieved desired functions while testing only tens of designs, indicating strong success rates relative to existing approaches. To extend the complexity of achievable designs, we integrated Proto with general-purpose AI coding agents to specify preliminary designs for a cancer-targeting lentiviral therapy, a multi-step human signaling pathway, and hundreds of protein complexes spanning the human proteome. In total, our results demonstrate how a small, intuitive set of general primitives can express biological designs of substantial complexity, expanding the practical reach of generative biology.

## 2. Results

### 2.1. Proto: Preliminaries and primitives

AI models for biology continue to rapidly proliferate, including generative models of protein sequence, protein backbones, RNA, and DNA (Kim et al., 2026), as well as predictive models of protein structure, gene expression, and many other phenotypes (Dubey and Shen, 2026; Park et al., 2025). However, despite this heterogeneity, the application of these tools to biological design converges on a remarkably simple set of shared abstractions (Hie et al., 2022; Listgarten and Jiang, 2026).

First, biological designs are encoded as a set of sequences (for example, of DNA, RNA, or proteins). Second, a candidate sequence is evaluated based on its desirability, for example, via a function *f* : ***χ***^*L*^ → ℝ that maps a sequence *x* of length *L* over vocabulary X to a scalar score *f* (*x*); here, we will assume that lower scores are better. Third, plausible candidate sequences need to be generated, that is, sampled from some distribution *p*(*x*) over sequence space. Finally, an optimization algorithm must steer generation toward desirable sequences, informally, sampling from *p*(*x*| *f* (*x*) is low) rather than from *p*(*x*) alone (Brixi et al., 2026; Brookes et al., 2019; Listgarten and Jiang, 2026).

A useful theoretical observation is that such a design task can be described by an energy-based model (EBM) (LeCun et al., 2006) with target distribution *π*(*x*) ∝ *p*(*x*) exp(− *f* (*x*)/*T*), in which the generator’s distribution *p*(*x*) acts as a prior, the constraint *f* acts as an energy, and the temperature *T* controls the trade-off between fidelity to *p* and minimization of *f* (**Figure S1A**; **Methods**). We can combine multiple constraints by summing their outputs into an aggregate constraint *f* = *f*_1_ + . . . + *f*_*k*_, which is equivalent to a product of experts (Hinton, 2002) over their Boltzmann factors exp(− *f*_*i*_ (*x*)/*T*) (Du and Kaelbling, 2024). Conveniently, this EBM formulation admits a compact graphical representation of a biological design campaign (**Figure S1B,C**). To obtain high-scoring sequences, we treat *π* as an energy landscape over which Metropolis-Hastings MCMC supplies stochastic moves, and where annealing the temperature *T* downward over the course of optimization favors the modes of *π*, providing a general black-box optimizer (Hastings, 1970; Kirkpatrick et al., 1983; Metropolis et al., 1953). Other optimizers, including gradient-based methods, are often more efficient in practice (**Methods**).

This framework and its components have analogs in natural and experimental biological design (**Figure 1C**) and also motivate the four primitives of the Proto language. In Proto, a sequence is a typed variable representing a DNA, RNA, protein, or ligand string. A constraint is any function that takes one or more sequences as input and returns a scalar output; this function can wrap a simple statistic like GC content or a neural network like AlphaFold. A generator is any procedure that proposes candidate sequences by sampling from a learned or pre-specified distribution, including autoregressive language models, diffusion models, or even a random sampler. An optimizer implements an iterative loop that minimizes an aggregate constraint score by accepting, rejecting, or ranking generated sequences. A minimal interface specifies each primitive in which generators produce sequences, constraints score them, and optimizers guide sequence generation toward better scores (**Figure 1D,E**).

To compose these primitives into complex design programs, Proto provides three higher-level interfaces that make Proto accessible across a range of computational expertise (**Figure 1D**). The first is a Python Application Programming Interface (API) in which sequences, constraints, generators, and optimizers are instantiated as objects and wired together programmatically. We provide this API both as an open-source library for local execution and as a hosted API for preconfigured cloud execution (**Figures S2, S3**, and **S4**). The second is a graphical interface that allows users to specify and connect sequence, generator, and constraint elements in a web application without writing code (https://proto.evodesign.org/; **Figure S5**). The third is an agentic workflow that integrates Proto with general-purpose AI coding agents through system prompts and a Model Context Protocol (MCP) interface to plan and write Proto programs from natural language descriptions, with additional integration into our web application. Beyond the core design tasks themselves, developing and hardening each of these interfaces required substantial engineering effort (**Discussion**).

### 2.2. Proto recapitulates prior design campaigns

To provide example programs and to demonstrate Proto’s flexibility, we first encoded five design tasks from the existing literature within the Proto design language and reproduced their generated results *in silico*. Despite spanning a wide array of molecular modalities, generative models, predictive models, and optimization strategies, all of these tasks can be expressed in the Proto language (**Figure 2**).

**Figure 2.**
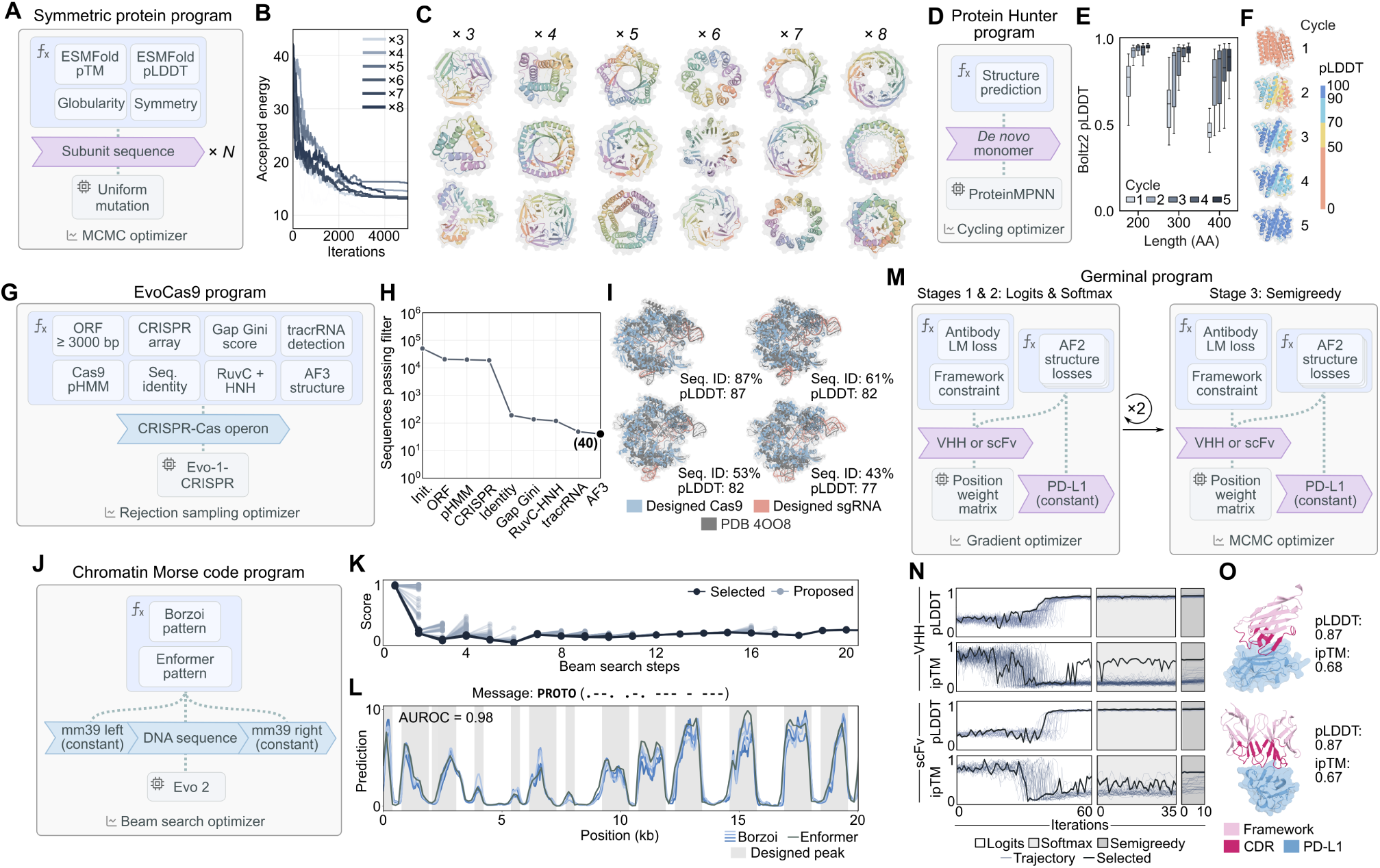
| Programming diverse design campaigns with Proto. (**A–C**) *De novo* symmetric protein homo-oligomers where uniform mutation serves as the generator; confidence, symmetry, and globularity of ESMFold-predicted structures serve as constraints; Metropolis-Hastings simulated annealing serves as the optimizer (**A**, program schematic; **B**, optimization trajectory), yielding predicted symmetric assemblies from trimers to octamers (**C**). (**D–F**) *De novo* protein monomers via Protein Hunter, alternating Boltz-2 structure prediction (constraint) with ProteinMPNN sequence redesign (generator) in a cycling optimizer (**D**), with designs converging to high structural confidence across cycles (**E, F**). (**G–I**) Multi-modal CRISPR-Cas systems where fine-tuned Evo 1 serves as the generator, bioinformatic filters and structure prediction serve as constraints, and a rejection-sampling procedure filters candidate sequences (**G, H**), producing 40 passing designs with putative Cas9 proteins and guide RNAs from 48,000 sampled loci (**I**). (**J–L**) Multi-kilobase chromatin accessibility design where Evo 2 serves as an autoregressive generator, Enformer and Borzoi serve as accessibility constraints, and a beam-search algorithm serves as the optimizer (**J, K**), yielding a 20-kb sequence predicted to encode the Morse-code message “PROTO” when integrated into the mouse genome (**L**; AUROC = 0.98). (**M–O**) *De novo* antibody CDR design via Germinal using multi-objective optimization over AlphaFold 2 (AF2) structure-based losses and AbLang antibody language model-based losses (**M, N**), where gradient descent serves as the optimizer, producing PD-L1 binders in VHH and scFv backbones with high AF2 confidence (**O**; pLDDT > 0.8, ipTM > 0.6).

We first considered the design of *de novo* symmetric protein homo-oligomers by Hie et al. (2022) (**Figure 2A–C**). These homo-oligomers were generated by iteratively proposing random mutations, predicting the resulting structure with ESMFold (Lin et al., 2023), and accepting mutations that improved structural confidence and symmetry. In Proto, uniform mutation is the generator, ESMFold confidence and scores for globularity and symmetry serve as constraints whose aggregate score defines an energy, and simulated annealing via Metropolis-Hastings MCMC implements the accept-reject optimizer loop that guides energy minimization. Our reimplementation yields predicted symmetric assemblies ranging from trimers to octamers (**Figures 2C** and **S6A,B**).

A complementary approach to protein design illustrates how different generators and optimizers can be substituted within the same language. The Protein Hunter method of Cho et al. (2025) designs *de novo* protein monomers by alternating between a diffusion-based structure predictor (Boltz-2, the Proto constraint) (Passaro et al., 2025) and an inverse folding model (ProteinMPNN, the Proto generator) (Dauparas et al., 2022) that redesigns the sequence to better match the predicted structure, where each iteration refines both sequence and structure toward a self-consistent solution (this alternating cycle defines the Proto optimizer) (**Figure 2D**). Across successive cycles, the designed monomers converge to high structural confidence (**Figures 2E,F** and **S6C**).

We next encoded a more complex, multi-modal design task of generating CRISPR-Cas systems (**Figure 2G–I**). Nguyen et al. (2024) used the Evo 1 genomic language model fine-tuned on CRISPR-Cas loci to generate operonic sequences encoding both a protein component and two noncoding RNAs that work together as a functional system. A series of computational filters, including CRISPR array identification and structure prediction of the candidate Cas9, are applied to retain only the most promising designs. In Proto, fine-tuned Evo 1 is the generator, each filter is a constraint, and a rejection sampling optimizer retains sequences passing all filters (**Figure 2G**). Sampling 48,000 sequences resulted in 40 passing designs containing putative Cas9 proteins and guide RNAs (**Figures 2I** and **S6D**), consistent with the previously reported *in-silico* filter pass rates for CRISPR-Cas generation (Nguyen et al., 2024).

The fourth task reimplements the design by Brixi et al. (2026) of 20-kilobase (kb) DNA sequences with predicted chromatin accessibility profiles that match a target pattern such as a Morse code message. Evo 2 autoregressively decodes candidate DNA sequences (generator), while Enformer and Borzoi (Avsec et al., 2021; Linder et al., 2025) predict the resulting chromatin accessibility tracks (constraints) (**Figure 2J**). A beam search optimizer retains and extends only those partial sequences whose predicted profiles best match the target, yielding final sequences that encode the desired pattern (**Figures 2K** and **S6E**). With this program, we designed a 20-kb region of DNA that is predicted to encode the Morse code message “PROTO” when integrated into the mouse genome (AUROC = 0.98) (**Figure 2L**).

Finally, we reimplemented Germinal, by Mille-Fragoso et al. (2025), which achieves *de novo* design of antibody CDRs using gradient-based, multi-objective optimization. AlphaFold 2 (AF2) confidence scores (Jumper et al., 2021), geometric properties of the predicted antibody-antigen structure, and the likelihood under an antibody-specific language (AbLang) model (Olsen et al., 2022) together define a composite constraint. Backpropagation through these constraints produces a sampling distribution over candidate sequences (generator), and the cycle of forward scoring and backward optimization repeats until convergence (optimizer) (**Figures 2M,N** and **S6F,G**). With this approach, we designed PD-L1 binders with *de novo* CDRs in both VHH and scFv backbones to have high AF2 confidence metrics (pLDDT > 0.8, ipTM > 0.6) (**Figure 2O**).

By recapitulating prior design campaigns with Proto, we demonstrate both the generalizability and expressiveness of our unified framework, which could in turn promote greater reproducibility of biological design pipelines.

### 2.3. Intron design for cell-line-specific gene regulation

While most synthetic gene regulation operates at the level of transcription (Brophy and Voigt, 2014; Gosai et al., 2024), splicing offers a relatively underexplored and complementary axis for gene regulation (Chen et al., 2025; Ling et al., 2022). However, rational design in this setting is challenging, as splicing outcomes are determined by a complex regulatory grammar alongside the combinatorial activity of RNA-binding proteins (Barash et al., 2010; Wang et al., 2008).

An important feature of Proto is that it enables generative design campaigns to rapidly benefit from new task-specific biological AI models. As an example, we integrated into Proto the recently developed sequence-to-function genomic model, AlphaGenome, which predicts cell-line-specific splice-site usage (SSU) as one of its outputs (Avsec et al., 2026). We then leveraged AlphaGenome to design intronic sequences with cell-line-specific effects on pre-mRNA splicing.

We used Proto to design intronic sequences with alternative splicing across human cell lines, with either increased or decreased mis-splicing in HepG2 or SH-SY5Y relative to K562 (**Table S1** and **Data S1**). AlphaGenome directly predicts SSU for both K562 and HepG2; given that AlphaGenome does not directly predict SSU for SH-SY5Y, but does predict SSU for neural cell ontologies, we tested how well these predictions would transfer to SH-SY5Y. The resulting Proto program generated a single intronic sequence in which AlphaGenome SSU and SpliceTransformer donor-acceptor splicing predictions served as constraints, a simple random proposal served as the generator, and designs were optimized with MCMC (**Figure 3A**; **Methods**). To make these designs robust to the sequence context in which they would ultimately function, we evaluated every constraint across diverse genetic contexts (**Figure 3A**).

**Figure 3.**
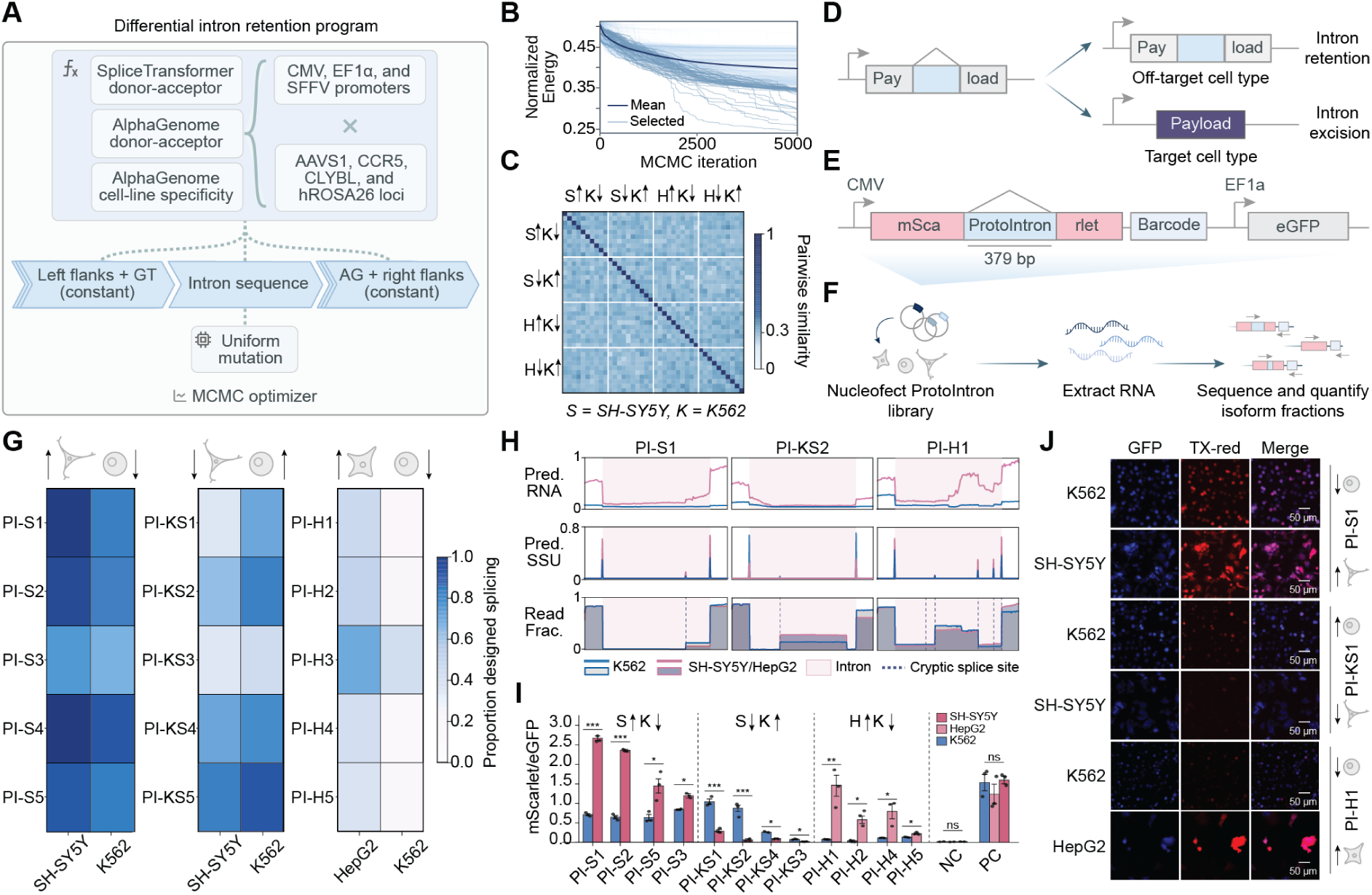
| Designing alternatively spliced introns in human cell lines with Proto. (**A**) SpliceTransformer and AlphaGenome donor-acceptor scores and AlphaGenome cell-line specificity serve as constraints, which are evaluated in several plasmid and human genomic contexts to minimize context-dependent variability. At each generation step, a uniform mutation generator proposes a change on top of a constitutively spliced intron or randomly initialized sequence; proposals are accepted or rejected using a MCMC optimizer. (**B**) Plot showing normalized energy (sum of energy scores divided by the number of constraints) over optimization iterations. Trajectories converge across all design campaigns: SH-SY5Y-correct/K562-mis-spliced (S↑K↓), SH-SY5Y-mis-spliced/K562-correct (S↓K↑), HepG2-correct/K562-mis-spliced (H↑K↓) and HepG2-mis-spliced/K562-correct (H↓K↑). (**C**) Pairwise similarity scores confirm sequence diversity across experimentally tested designed introns within and between campaigns. (**D**) Intron retention in off-target cells sequesters a downstream payload while excision in target cells enables its translation. (**E**) Schematic of plasmid setup for screening generated ProtoIntrons. ProtoIntrons are inserted within the mScarlet coding sequence, followed by a constant barcode for isoform quantification, in a dual-reporter construct where eGFP is independently driven by EF1*α*. Successful splicing restores full-length mScarlet translation. (**F**) Isoform fractions and splicing variability are quantified by RNA sequencing across transfected cell lines. (**G**) Per-construct heatmaps of proportion of spliced products matching designed splicing confirm cell-line differential splicing across multiple cell-line combinations and designed directions. PI, ProtoIntron. (**H**) AlphaGenome-predicted splice site usage aligns well with measured cryptic and canonical splicing events for representative differentially spliced ProtoIntrons, while predicted RNA expression diverges more substantially. (**I**) mScarlet/eGFP ratios measured by fluorescence microscopy generally confirm differential payload translation at the protein level. S, SH-SY5Y; K, K562; H, HepG2; PC, HBB2c intron; NC, HBB2c intron reversed. (**J**) Representative fluorescence microscopy images demonstrating cell-line-specific variability in mScarlet expression across design campaigns.

Across all four design directions, optimization found low-energy intron designs (**Figure 3B**) while preserving substantial sequence diversity among top candidates (**Figures 3C** and **S7**). From this pool of generated sequences, we experimentally screened 65 candidates total from across design directions, which we refer to as ProtoIntrons.

To evaluate these designs in a functional context, we embedded each ProtoIntron within the mScarlet coding sequence of a dual-reporter construct such that successful excision restores full-length mScarlet (**Figure 3D,E**). After nucleofecting each ProtoIntron library into its corresponding cell lines, we quantified per-construct isoform fractions via next-generation sequencing (NGS) (**Figure 3F**; **Methods**).

Of the ProtoIntrons successfully assayed by NGS, 32% exhibited significant differential splicing in the designed direction (**Figures 3G, S8**, and **S9**); for comparison, a previous study that performed computational design of alternative splicing achieved a success rate of <7% after testing ∼ 10^3^ sequences (Chen et al., 2025), albeit in different experimental systems. One of our designs, PI-KS1, which was optimized for lower mis-splicing in K562, exhibited just 36% splicing in SH-SY5Y compared to 71% within K562. Other ProtoIntrons, such as PI-S1 and PI-H1, also showed the same target-cell-line bias, with PI-S1 showing greater splicing in SH-SY5Y over K562 (by a difference of 11%) and PI-H1 showing greater splicing in HepG2 over K562 (by a difference of 30%). AlphaGenome-predicted SSU closely matched the observed canonical and cryptic splicing events in the designed introns, with predicted RNA expression diverging more appreciably (**Figures 3H, S10, S11**, and **S12**; **Data S1**).

We further measured normalized mScarlet expression by fluorescence microscopy for the top-performing ProtoIntrons in each design direction (**Figures 3I,J** and **S13**). ProtoIntrons generally drove significantly higher normalized mScarlet fluorescence in the target cell line relative to the off-target line across design directions, suggesting that differential splicing translated into differential payload expression at the protein level. We also provide further evidence for differential splicing via nanopore long-read sequencing and gel electrophoresis in line with both NGS and fluorescence measurements (**Figure S14**).

Notably, while the Proto program only specified SSU-based constraints at the intron boundaries, we observed that the designed sequences also leveraged cryptic splice usage within the designed intron to achieve differential gene regulation, which was an emergent mechanism that we did not directly specify (**Figures S10**, **S11**, and **S12**). For example, many successful designs introduced intermediate cryptic splice sites that diverted SSU away from the expected splice sites in the non-target cell line (**Figure S15A–D**). These cryptic events contain hallmarks of genuine splice acceptors and were enriched at positions of elevated AG-dinucleotide density (**Figure S15E–H**). Cryptic splice junctions also coincided with AlphaGenome-predicted splice sites while remaining largely independent of canonical-site strength (**Figure S15I,J**). These results demonstrate how Proto optimization can generate sequences that fulfill the user-specified high-level design goals via emergent mechanisms.

Lastly, we asked how well our *in-silico* metrics predicted experimental success. AlphaGenome had largely accurate predictions, placing 85% of observed junctions within 20 base pairs of a predicted junction (**Figure S16A**). However, no single constraint metric separated differentially spliced ProtoIntrons from non-differentially spliced generations (**Figure S16B–D**). This suggests that the combination of constraints, rather than a single metric, facilitated differential splicing design, consistent with the multi-objective optimization at the core of Proto. Although the HepG2-mis-spliced/K562-spliced design objective did not produce any designs with significant differential splicing (**Figure S8**), we were able to achieve low-*N* success in the three other design conditions without high-throughput screening of thousands of sequences as done previously (Chen et al., 2025). Composing complementary signals from predictive models of regulatory sequences with Proto provides a practical route to cell-line-specific splicing design.

### 2.4. Designing synthetic promoter-repressor interactions

We next asked whether Proto could co-design interacting components across modalities. Promoter-repressor pairs provide a compact test case for this challenge, requiring coordinated design of a DNA regulatory element and a protein that recognizes it to modulate transcription. Their function depends on sequence-specific protein-DNA recognition and, in many natural systems, cooperative protein-protein interactions (Lewis et al., 1996). Such pairs are also important building blocks for synthetic regulatory systems, including toggle switches, oscillators, and layered logic circuits (Elowitz and Leibler, 2000; Gardner et al., 2000; Nielsen et al., 2016). However, many existing systems draw from a limited repertoire of natural repressors, motivating approaches that can generate new promoter-repressor interactions with target-biased recognition and reduced reliance on natural part libraries.

We first designed synthetic *σ*70 promoters, which we term ProtoPromoters, using a two-stage Proto program (**Figure 4A**). Using Evo 2 to generate sequence proposals, we leveraged rejection sampling to select candidates likely to be strong promoters in *E. coli* using constraints based on Promoter Calculator activity predictions (LaFleur et al., 2022), biologically matched −35 and −10 boxes from *σ*70 promoters, and orthogonality to native transcription-factor motifs. These initial promoter sequences were then further optimized using a simple uniform mutation proposal and MCMC, with an additional constraint being added to introduce palindromic operator sites directly blocking the −35 box, −10 box, or transcription start site in selected candidates for downstream repressor binding.

**Figure 4.**
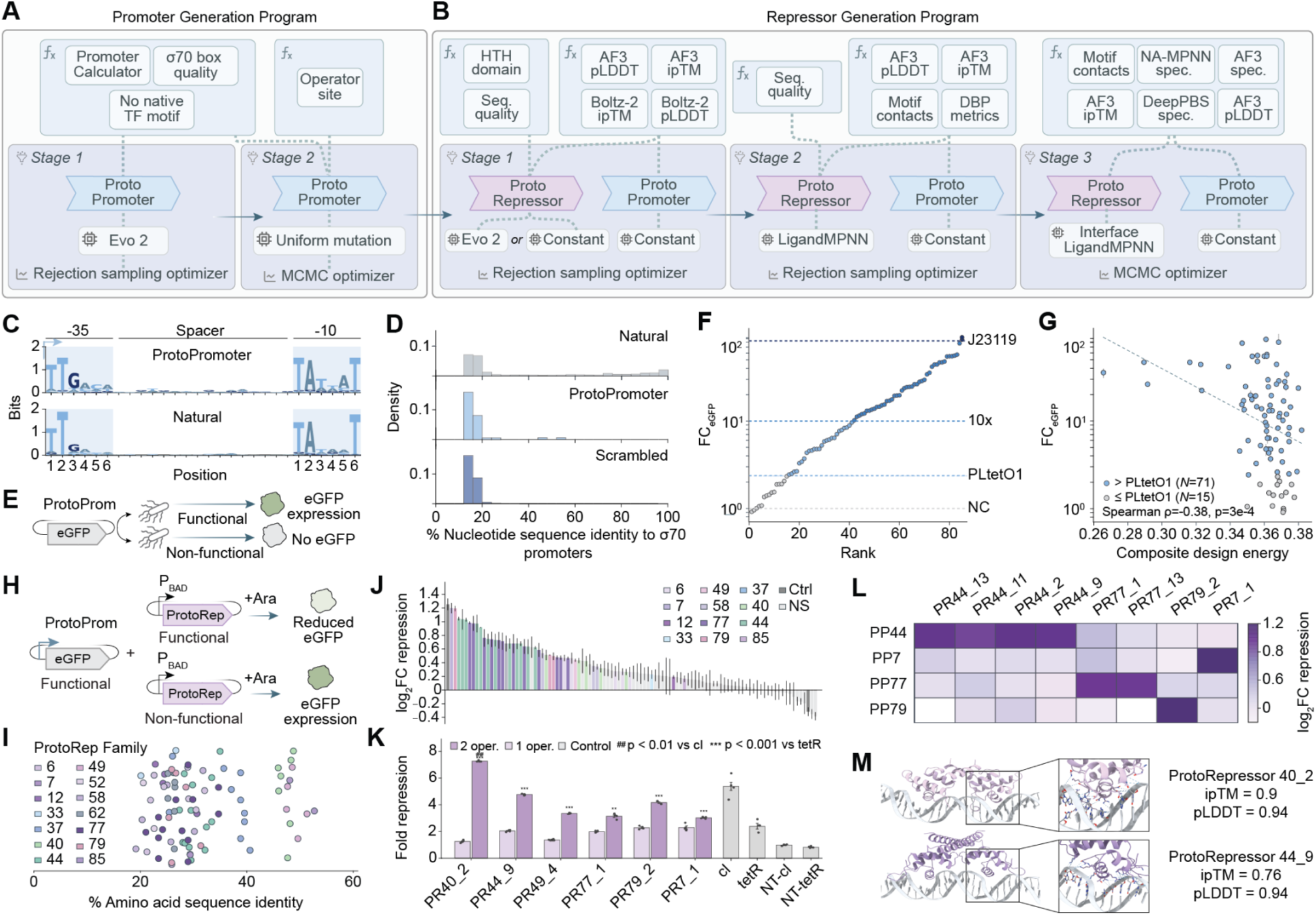
| Design of synthetic promoter-repressor pairs with Proto. (**A**) ProtoPromoter generation pipeline. Stage 1 uses Evo 2 with rejection sampling to generate candidate promoter sequences filtered for predicted promoter activity, *σ*70 box quality, and absence of native transcription factor motifs to optimize for strength and orthogonality. Stage 2 applies uniform mutation with an MCMC optimizer to refine selected sequences and introduce palindromic operator sites. (**B**) ProtoRepressor generation pipeline. Stage 1 uses Evo 2 and natural sequences with rejection sampling to select for initial repressor sequences showing predicted binding to ProtoPromoters as measured by AlphaFold 3 (AF3) and Boltz-2 pLDDT/ipTM. Stage 2 applies LigandMPNN with rejection sampling, scoring motif contacts and Rosetta-based DNA-binding protein metrics. Stage 3 refines the binding interface with LigandMPNN and MCMC optimization to promote specificity, incorporating NA-MPNN specificity, DeepPBS specificity, and AF3 specificity metrics. (**C**) Sequence logos of candidate ProtoPromoters showing conservation at the −35 and −10 boxes for designed ProtoPromoters versus natural *σ*70 promoters. (**D**) Distribution of percent sequence identity to *σ*70 promoters for natural, ProtoPromoter, and scrambled sequences. ProtoPromoters are highly diverse and share limited sequence identity to natural *σ*70 promoter sequences. (**E**) Schematic of the bacterial eGFP reporter assay for ProtoPromoter functional screening. ProtoPromoters are cloned upstream of eGFP, reading out transcriptional activity as fluorescence. (**F**) Ranked eGFP fold change for designed ProtoPromoters; dashed lines indicate the measured strength of the J23119 constitutive promoter, a 10× fold change threshold, PLtetO1, and a no-promoter control (NC). ProtoPromoters are generally highly active, with many exceeding a 10-fold change in eGFP expression and the strongest matching or approaching the constitutive promoter J23119. (**G**) Composite design energy has a moderate inverse correlation with eGFP fold change and generally selects for transcriptionally active promoters, validating the multi-objective scoring function. Error bars; standard error of mean. (**H**) Schematic of the ProtoRepressor screening assay. eGFP expression driven by a functional ProtoPro-moter (ProtoProm) serves as the reporter, and a candidate ProtoRepressor (ProtoRep) is induced from a separate arabinose-dependent promoter. Functional ProtoRepressors repress the ProtoPromoter and reduce eGFP, while non-functional variants leave eGFP expression intact. (**I**) Distribution of highest BLAST amino-acid sequence identities (horizontal axis) of experimentally screened Pro-toRepressors across 14 ProtoPromoters indicates high sequence novelty of generated ProtoRepressors. The vertical axis corresponds to random jitter. ProtoRep Family, ProtoRepressor candidates designed to target a unique ProtoPromoter. (**J**) log_2_ fold change (log_2_FC) repression across ProtoPromoter-ProtoRepressor combinations as measured by flow cytometry, with colors indicating repressor family. ProtoRepressors across diverse promoter families show significant repression over negative controls. Error bars, standard error of mean. Ctrl, tetR positive and non-targeting control; NS, non-significant over fold change = 1. (**K**) Fold repression for selected top ProtoRe-pressor candidates configured with a single operator site (1 oper.) or two operator sites (2 oper.). Addition of a second operator site increases repression strength across candidates, compared with a constitutive lambda repressor control (cI), tetR, and a non-targeting tetR control (tetR NT). ##, *P* < 0.01 versus cI; ***, *P* < 0.001 versus tetR. Bars show mean; circles, individual replicates. (**L**) Cross-repression heatmap showing log_2_ fold change repression of candidate repressors across cognate and non-cognate ProtoPromoter targets. Strongest ProtoRepressor candidates generally demonstrate specificity toward their target ProtoPromoters. (**M**) AF3-predicted structures of protein-DNA complexes for ProtoRepressor 40_2 and ProtoRepressor 44_9, with insets showing the recognition helix inserted into the major groove with base and backbone contacts.

Designed promoters retained the expected information content at the −35 and −10 boxes (**Figure 4C**) yet were highly diverse, sharing limited sequence identity with natural *σ*70 promoters and approaching scrambled controls in their identity distribution (**Figures 4D** and **S17**). Having established that ProtoPromoters preserved canonical

promoter grammar without simply recapitulating natural sequences, we next evaluated their functional activity. Each candidate sequence was cloned upstream of eGFP and assayed for induced transcriptional activity in *E. coli* by measuring resultant fluorescence (**Figure 4E**; **Data S2**). Overall, ProtoPromoters exhibited strong activity, with 45 of 86 candidates driving eGFP expression more than 10-fold above the no-promoter control and 71 of 86 producing higher activity than a strong existing promoter, PLtetO1 (**Figure 4F**). Consistent with this enrichment for active promoters, composite design energy was moderately inversely correlated with promoter strength, as measured by eGFP fold change (Spearman *ρ* = −0.38, two-sided *t*-distributed *P* = 3.0 × 10^−4^; **Figure 4G**).

We next designed cognate ProtoRepressors against operator sites from ProtoPromoters spanning a variety of activity levels. Against a panel of ProtoPromoters that were held constant during generation, we implemented a three-stage repressor generation program that jointly optimized protein sequence, structural confidence, DNA binding, and promoter-specific recognition (**Figure 4B**). In the first stage, candidate helix-turn-helix repressor backbones were generated against ProtoPromoter operator sites, with designs filtered by sequence quality and structure-prediction metrics. In the second stage, repressor sequences were further optimized using LigandMPNN and evaluated for predicted motif contacts, protein-protein interaction strength, and DNA-binding features. Finally, interface residues were refined against each target operator using MCMC, incorporating specificity metrics from NA-MPNN, AlphaFold 3 (AF3), and DeepPBS to select repressors predicted to bind the intended promoter while minimizing off-target recognition (**Methods**). Across these *in-silico* metrics, selected ProtoRepressors matched or exceeded natural helix-turn-helix proteins on binding confidence, interface contact richness, and predicted specificity while preserving native-like packing (**Figure S18**).

To evaluate repression, we co-transformed *E. coli* with the previously designed functional ProtoPromoter-eGFP constructs and a separate plasmid expressing each ProtoRepressor under an arabinose-inducible promoter, with functional repressors expected to reduce eGFP fluorescence compared to scrambled repressor controls (**Figures 4H** and **S19; Data S2**). The screened ProtoRepressors were highly diverse, with most designs sharing less than 50% sequence identity to natural proteins across targets from 14 ProtoPromoters (**Figure 4I**). Flow cytometry of the ProtoRepressor library revealed a high functional hit rate, with 46% of designs significantly reducing eGFP expression relative to non-targeting controls across 12 ProtoPromoter targets, with just 2 ProtoPromoters failing to have any successful designs. Although most active candidates produced only modest repression, a subset achieved stronger activity, with 9% producing more than 1.5-fold reduction in eGFP expression (cf. 2.38 fold repression for tetR positive control; **Figure 4J**). As operator valency can tune transcriptional repression, we next tested whether adding a second palindromic operator site directly downstream of the transcription start site (TSS) would improve repressor performance. The two-operator architecture increased repression across candidates relative to the single-site design, with the strongest designs exceeding strong constitutive cI-mediated repression and approaching or surpassing tetR-mediated repression (ProtoRepressor 40_2, *P* = 9.8 × 10^−3^ versus cI; ProtoRepressor 44_9, *P* = 4.5 × 10^−7^ versus tetR by one-sided *t*-test; **Figure 4K**).

Notably, the strongest ProtoRepressors preferentially repressed their target ProtoPromoters across the tested cognate and non-cognate pairings, consistent with target-biased recognition rather than purely generic DNA binding (**Figure 4L**). For example, ProtoRepressor 44_9 exhibited no significant repression against ProtoPromoter 7, 77, or 79 while exhibiting 2.0 fold repression against designed target ProtoPromoter 44. Consistent with this, AF3 models of top complexes (ProtoRepressor 40_2 and 44_9) positioned the recognition helix within the DNA major groove with both base-specific and backbone contacts (**Figure 4M**).

As in the intron designs, no single *in-silico* metric cleanly separated functional from non-functional repressors, suggesting that repressor activity was better captured by a composite of structure-prediction-confidence, interface-contact, and specificity signals (**Figure S20**; **Table S2**). Notably, these repressors were designed without a task-specific generative model trained to produce DNA-protein binders. Instead, Proto integrated general-purpose generative models, structure predictors, and biophysical scoring functions into a consensus workflow that prioritized designs with plausible folding, DNA-binding geometry, and target specificity. Proto yielded functional promoter-repressor pairs with favorable success rates compared to prior efforts in sequence-specific protein-DNA design, where the highest reported success rates range from 3-13% against a lower fraction of DNA targets (Glasscock et al., 2025; Sehgal et al., 2026), albeit using different assays. Moreover, achieving functional repressors not only requires DNA binding, but also realizing other complex objectives such as protein-protein cooperativity. Together, the joint design of active promoters and cognate repressors demonstrates that Proto can coordinate diverse generative models, biophysical models, and structural predictors to generate experimentally testable interacting regulatory components after low-*N* screening.

### 2.5. Agentic programming of complex biological systems

An emerging programming paradigm leverages AI coding agents that pair a reasoning large language model with command-line tool use to perform increasingly sophisticated software engineering tasks from natural language instructions alone (Karpathy, 2025; Chen et al., 2021; OpenAI, 2026; Anthropic, 2026). Because Proto may have a nontrivial learning curve for many researchers, we likewise hypothesized that AI agents could enable these researchers to write Proto programs through simple natural language instruction and steering. Moreover, Proto’s standardized tool API removes much of the environment-configuration burden on which agents often fail or expend substantial effort (Luebbert, 2026), and Proto’s compositional primitives help facilitate more complex, multi-objective design tasks.

We therefore sought to demonstrate these capabilities by using AI coding agents to write Proto programs at the scale of tens or hundreds of genes and with tens or hundreds of functional constraints, which would be difficult even for a human expert to program by hand (**Figure 5A**). We began with the task of protein diversification in which generative models sample variants of natural proteins with similar structures and functions. Diversification of individual proteins, such as GFP (Hayes et al., 2025) and Cas9 (Ruffolo et al., 2025), has been an important proof-of-concept that generative models can productively explore protein sequence space. Language-model-guided diversification can in turn improve the efficiency of directed evolution campaigns (Hie et al., 2024; Shanker et al., 2024).

**Figure 5.**
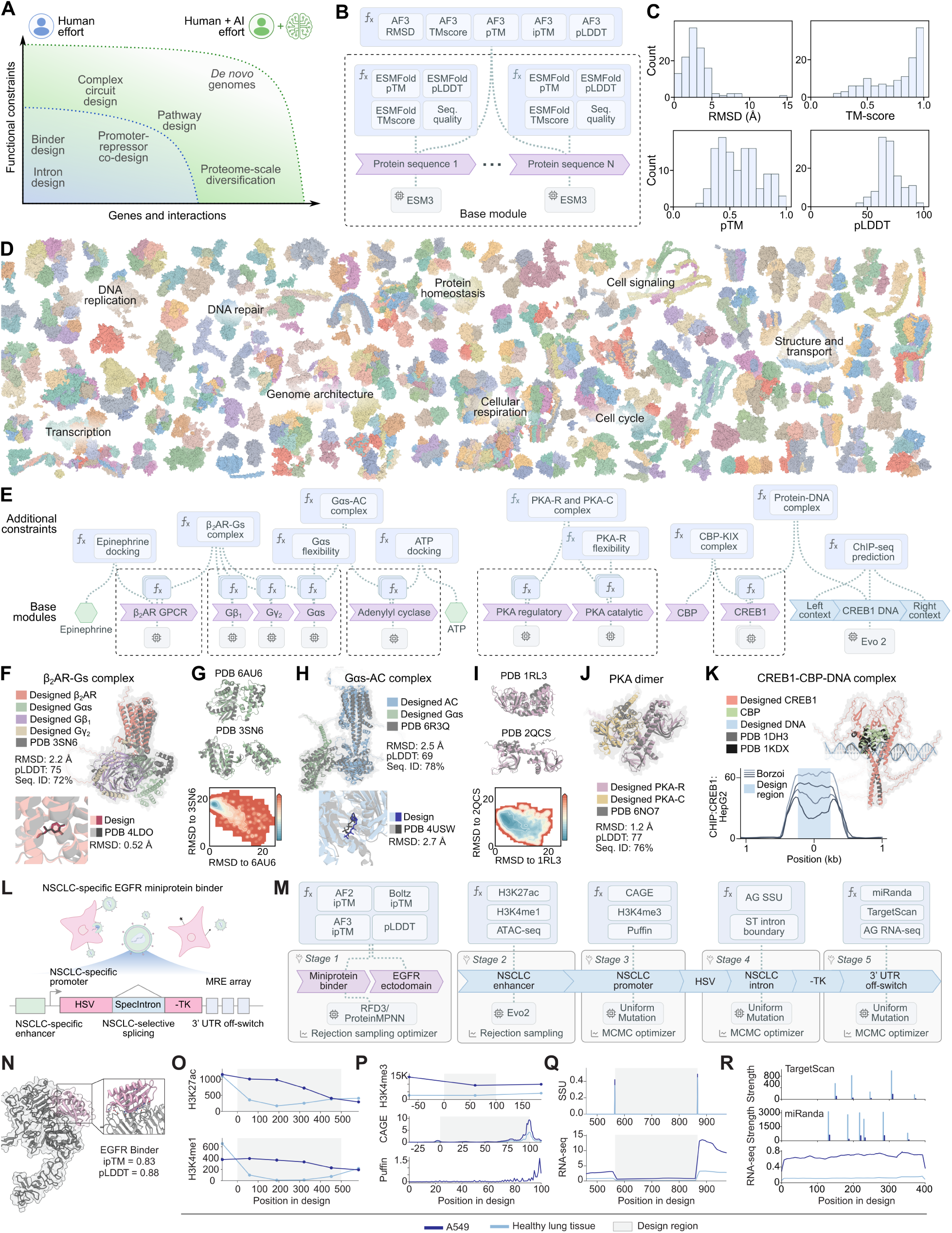
| AI agents facilitate complex design specification with Proto. (**A**) A general-purpose AI coding agent with human interaction can translate natural-language steering instructions into a Proto program, which both enables users with limited domain expertise to write basic programs and also extends the frontier of what expert users are able to program. (**B-D**) Proteome-scale complex diversification. (**B**) Schematic of the diversification module in which wild-type human sequences seed each subunit, ESM3 proposes variants, and per-subunit constraints on structural confidence (ESMFold pLDDT/pTM), fold consistency (TM-score, RMSD to native), and sequence complexity drive a joint MCMC optimizer, with each complex rescored as a multimer by AF3. (**C**) For designed complexes with a known experimental structure, these distributions plot RMSD of the AF3-predicted, ESM3-diversified complex to the native structure (median 1.9 Å), AF3 TM-score to native (median 0.80), as well as AF3 structural confidence metrics pTM (median 0.65) and pLDDT (median 71). (**D**) Representative AF3-predicted complexes, along with broad functional category labels, illustrating the scale of this diversification campaign. (**E-K**) Signaling-pathway redesign of the *β*2-adrenergic axis. (**E**) Pathway schematic spanning *β*2AR, G*αβγ*, adenylyl cyclase, PKA, CREB1, and a CREB-responsive DNA element, with function-specific constraints layered onto the base ESM3/AF3-diversification module. Elements that are shown without attached generators (epinephrine, ATP, CBP, and the CREB DNA contexts) were held constant during optimization. AF3-predicted designed complexes of *β*2AR-Gs (**F**, top), epinephrine docked in *β*2AR (**F**, bottom), G*α*s-adenylyl cyclase (**H**, top), adenosine triphosphate (ATP) docked in adenylyl cyclase (**H**, bottom), and the regulatory and catalytic subunit heterodimer of PKA (**J**) aligned to their experimental structures. BioEmu conformational-ensemble analysis showing that designed G*α*s (**G**) and the PKA regulatory subunit (**I**) are predicted to populate two known native conformations of each protein. (**K**) The top shows ESM3-diversified CREB1, Evo 2-generated DNA, and native CBP as predicted in a complex by AF3, compared to native structures of the CREB1-DNA and CREB1-CBP interactions. The bottom shows Borzoi-predicted CREB1 ChIP-seq signal over the Evo 2-generated DNA element. (**L-R**) Non-small cell lung cancer (NSCLC)-selective therapeutic payload. (**L**) Design of a multi-layer lentivirus-based gating strategy in which an EGFR-targeting miniprotein binder biases cell entry toward NSCLC cells, an NSCLC-selective enhancer and promoter drive HSV-TK transcription, an embedded intron enables NSCLC-selective splicing of the HSV-TK coding sequence, and a 3^′^ UTR microRNA-response-element array suppresses residual expression in off-target contexts. (**M**) Proto program spanning five design stages: (i) miniprotein binder design against the EGFR ectodomain, (ii) NSCLC-selective enhancer design, (iii) NSCLC-selective promoter design, (iv) NSCLC-selective intron design, and (v) 3^′^ UTR off-switch design, with constraints on protein-structure confidence, epigenomic activity, promoter-associated signals, splice-site usage, and miRNA-mediated repression. (**N**) AF3-predicted structure of the designed EGFR-binding miniprotein in complex with the EGFR ectodomain shows high confidence structure and several interface contacts. (**O**) AlphaGenome predicted H3K27ac and H3K4me1 signal across the designed enhancer in A549 and healthy lung tissue in GAPDH integration context, showing A549-biased enhancer-associated activity relative to healthy lung tissue. (**P**) AlphaGenome predicted H3K4me3, CAGE, and Puffin promoter activity across the designed promoter in A549 and healthy lung tissue shown in GAPDH integration context. CAGE and Puffin predict a TSS at the end of the designed promoter, with reduced activity in healthy lung tissue. (**Q**) AlphaGenome predicted splice-site usage and RNA-seq signal across the designed NSCLC-selective HSV-TK intron shows a modest preference toward intron excision in A549 relative to healthy lung tissue. (**R**) Designed 3^′^ UTR off-switch with stronger TargetScan- and miRanda-predicted miRNA binding-site activity in healthy lung tissue, avoidance of NSCLC-high miRNA targets, and AlphaGenome-predicted low RNA-seq signal at selected MREs.

While these previous studies diversified single proteins, many cellular functions are carried out by protein complexes for which diversification would require jointly optimizing all constituent subunits in the complex. We therefore asked whether Proto could scale this approach from a single protein to hundreds of protein complexes spanning the human proteome (**Figure 5B-D**). Using a combination of manual and AI-based curation, we assembled a set of 249 human protein complexes comprising 797 individual genes (**Data S3**). We then used an AI agent to automatically generate a Proto program for one or more complexes, each encoding the diversification of all unique subunits (**Figure 5B**). Within each program, wildtype human sequences serve as initial seeds, ESM3 proposes sequence variants for each subunit, and subunit-level constraints on structural confidence, fold consistency, and sequence complexity guide an MCMC optimizer. For each complex, the agent used Proto to sample at least five sets of sequences, scored each set as a multimer with AF3, and selected the highest-confidence set by pLDDT (**Methods**).

Across the designed complexes, AF3 predicts structures with low RMSD to native references (median 1.9 Å) and high TM-scores (median 0.8) despite substantial sequence divergences from wildtype (median of 80% sequence identity) (**Figures 5C,D** and **S21A,B**), similar to those reported in previous diversification campaigns (Ruffolo et al., 2025). Together, these results show that the agent, using Proto, reproduced the structural fidelity of single-protein diversification while diverging substantially in sequence, now across hundreds of complexes simultaneously. AF3 prediction with high confidence and similarity to the native structure are standard *in-silico* acceptance criteria used in diversification campaigns prior to experimental testing (Ruffolo et al., 2025); we emphasize, however, that structure-prediction-based metrics only serve as plausibility filters rather than a guarantee of function.

To go beyond structure-preserving diversification, we then used Proto to encode more complex functional constraints across an entire intracellular signaling cascade (**Figure 5E-K**). This task requires designing multi-modal interactions involving small-molecule ligands or nucleic acid elements, as well as the protein conformational dynamics that mediate signaling interactions.

We chose the canonical *β*2-adrenergic signaling axis, spanning the *β*2-adrenergic receptor (*β*2AR), the heterotrimeric G protein G*αβγ*, adenylyl cyclase (AC), protein kinase A (PKA), the transcription factor CREB1, and a CREB-responsive DNA element. Importantly, we encoded function-specific constraints relevant to signal transduction on top of the base module of ESM3 diversification and AF3 structural scoring (**Figure 5E**). These constraints include epinephrine binding at the receptor; conformational flexibility of G*α*s between its GDP- and GTP-bound states; ATP binding by adenylyl cyclase; regulatory-subunit flexibility of PKA; and transcription-factor dimerization, coactivator recruitment, and DNA binding by CREB1 (**Figure S21C-G**). A separate optimization stage also uses Evo 2 to generate a genomic region with high Borzoi-predicted CREB1 ChIP-seq signal.

Despite sequence diversification, AF3 predicts that the designed pathway retains its native interaction geometries across all states, with the *β*2AR-Gs complex aligning to experimental structures at 2.2 Å RMSD and 72% average sequence identity and the G*α*-adenylyl cyclase complex at 2.5 Å RMSD and 78% average sequence identity (**Figures 5F,H,J** and **S21C,D**). BioEmu analysis further predicts that the designed G*α*s and PKA regulatory subunits populate the functional conformations observed for their wildtype counterparts (**Figures 5G,I** and **S21E,F**), and Borzoi predicts a strong CREB1 ChIP-seq signal centered on the designed DNA element (**Figures 5K** and **S21G**).

Beyond diversification, coupling agents with Proto may provide a powerful framework for specifying synthetic systems that can potentially achieve novel, therapeutically relevant functions, a process that requires exploration across many possible design strategies. We therefore used Proto to specify cell-type-selectivity of a therapeutic payload that targets non-small cell lung cancer (NSCLC) (**Figure 5L**). The core therapeutic mechanism is a herpes simplex virus thymidine kinase (HSV-TK) transgene delivered by lentiviral vector. HSV-TK converts the prodrug ganciclovir into a toxic metabolite that induces cell death (Moolten, 1986). The design challenge is to ensure that functional HSV-TK is produced only in NSCLC cells, so that ganciclovir administration selectively kills tumor cells while sparing healthy tissue.

Our Proto program enforces this selectivity through five optimization stages, each controlling a distinct layer of regulation. The first stage designs a miniprotein binder against the EGFR ectodomain, a tumor-associated surface receptor frequently expressed in NSCLC, to bias lentiviral entry toward tumor cells (Hirsch et al., 2003; Sharma et al., 2007). The second and third stages design an enhancer and a promoter, respectively, that preferentially drive transcription in NSCLC cells. The second stage generates the enhancer with Evo 2 and the third stage generates the promoter with uniform mutation, optimizing both against cell-type-specific epigenomic and transcriptomic signatures predicted by Borzoi and AlphaGenome; the promoter was also optimized with the Puffin transcription-initiation predictor (Avsec et al., 2026; Dudnyk et al., 2024; Linder et al., 2025). The fourth stage designs an intron embedded within the HSV-TK coding sequence with NSCLC-selective splicing such that correct excision would reconstitute a functional HSV-TK open reading frame, whereas alternative splicing would disrupt the reading frame and diminish expression. The fifth stage designs a 3^′^ UTR off-switch composed of microRNA response elements that silence any residual transcript in non-NSCLC contexts (**Figure 5M**; **Methods**).

Across these layers, the program converged on a coherent predicted NSCLC-selective design in which each subprogram contributed a distinct regulatory bias. At the cell-entry layer, Proto produced a 110-aa EGFR binder with consistent interface confidence across three independent structure-prediction families, with ipTM values ranging from 0.81-0.83 (**Figures 5N** and **S22A,B**). At the transcriptional layer, the enhancer and promoter subprograms each recovered strong A549-selective regulatory signatures. The designed 500-nt enhancer showed approximately 9-fold higher H3K27ac and 1.8-fold higher accessibility in A549 than in healthy lung, consistent with predicted enhancer activity (**Figures 5O, S22C,D**, and **S23A-C**). The promoter subprogram likewise produced a strong A549-biased chromatin profile as measured by Borzoi and Puffin, showing moderate Puffin-predicted promoter activity with retained A549 selectivity, including 2.7-fold higher H3K4me3 (**Figures 5P, S22E,F**, and **S23D,E**).

Both the predicted CAGE signal and Puffin showed a peak toward the 3^′^ end of the design, suggesting the presence of a transcription start site. Interestingly, these predictions, while optimized using AlphaGenome, were generally directionally concordant with a separate sequence-to-function model, Borzoi, that was not used during optimization, increasing confidence that the predicted A549 selectivity may reflect genuine signal rather than overfitting to a single predictor (**Figure S22**). At the RNA-processing layer, the designed 301-nt intron contributed a smaller but complementary bias, with NSCLC-selective splice-site usage up to 1.3-fold higher in A549 than in healthy lung tissue (**Figures 5Q** and **S23F-H**). Finally, at the post-transcriptional layer, the 3^′^ UTR off-switch introduced five healthy-high miRNA target sites in the best candidate, giving 7.1-fold higher predicted repression in healthy lung than A549 while avoiding tumor-high miRNA sites (**Figures 5R** and **S23I**).

In total, the full NSCLC program consists of five optimization stages, three distinct generators (RFdiffusion3 and ProteinMPNN, Evo 2, and uniform mutation) (Butcher et al., 2025; Dauparas et al., 2022; Brixi et al., 2026), two optimizers (rejection sampling and MCMC), and over a dozen constraint functions spanning structure prediction, epigenomic track prediction, splice-site modeling, and miRNA targeting (**Figure S23J**). Several of the final subprograms were identified only after iterative exploration with an agentic interface (**Methods**), which evaluated alternative generators, optimizers, and constraints to prioritize those most likely to succeed for each design objective. Together, these three examples illustrate how Proto can compose programs with diverse generative models (such as autoregressive generation of DNA, uniform mutation of RNA, and diffusion-based diversification of proteins), predictive models (of epigenomic activity, ligand binding, protein structure, or conformational dynamics), and biological tasks (diversification of protein multimers and signaling pathways, cell-type-specific gene regulation) at the scale of tens or hundreds of genes. The structured grounding provided by Proto combined with natural language steering of increasingly capable general-purpose AI coding agents can expand both the accessibility and the scale of generative biology.

## 3. Discussion

In this work, we have distilled the methodologies and best practices developed by the biological design community into a unifying programmatic framework. Proto allows researchers to express complex design tasks by representing powerful biological AI models as simple primitives, which can in turn be composed and reused within a shared language. We experimentally validated designs spanning all three modalities of the central dogma (RNA splicing, DNA-protein interactions) in prokaryotic and eukaryotic systems, achieving functional synthetic sequences after low-throughput testing of only tens of candidates in each case. We release Proto as an open resource, including software infrastructure and a user interface, at https://proto.evodesign.org/.

The development of Proto required substantial engineering effort. The current ecosystem of biological AI models and computational tools was not designed for general-purpose composition; models and tools are commonly released as independent artifacts, each with distinct software dependencies, hardware requirements, model weights, databases, and input/output conventions that often conflict. This lack of compatibility and standardization presents a major barrier for design campaigns that compose multiple tools in the same workflow. Design pipelines that do integrate different methods (Boyd et al., 2025; Hopf et al., 2026; Mille-Fragoso et al., 2025; Ovchinnikov et al., 2025; Pacesa et al., 2025; Roney and Ovchinnikov, 2022) are often limited to a small set of mutually compatible components and are tailored to a specific problem, such as binder design. Proto provides standardized and open-source infrastructure for over 120 tools (**Figure S2**), including model weight and database provisioning, environment configuration, resource-aware execution, and standardized data exchange, which can be explored at https://proto.evodesign.org/tools.

Proto is part of a growing line of work that leverages inference-time search and multi-objective composition instead of monolithic, task-specific models. Many state-of-the-art design tools, such as BindCraft (Pacesa et al., 2025), Germinal (Mille-Fragoso et al., 2025), and Protein Hunter (Cho et al., 2025), achieve their performance not by training a single bespoke model, but by combining a suite of existing models at inference time. Proto generalizes this pattern beyond *de novo* protein design to compose arbitrary models, modalities, and objectives. Moreover, while many biological design campaigns to date have optimized for a single property, such as enzyme activity or binding affinity, Proto natively performs multi-objective optimization.

Related efforts within biology, like Cello, SBOL, and Eugene, have developed a programmatic specification for genetic circuits through simple, composable rules defined on curated part libraries. While important conceptual milestones, they primarily operate over predefined genetic parts and circuit abstractions and are limited in their ability to generalize to new sequences or span design tasks across scales (Bilitchenko et al., 2011; Galdzicki et al., 2014; Jones et al., 2022; Nielsen et al., 2016). We also note that the sequence generation, scoring, and optimization functionality of tools like Cello can be incorporated into Proto.

Proto also fits within a broader history of programming language design. A Proto program specifies a probability distribution over sequences conditioned on desired function, and compilation performs approximate inference of the modes of *p*(sequence | function). Proto is therefore an instance of a probabilistic programming language (Bingham et al., 2018; van de Meent et al., 2021; Tran et al., 2017) specialized to biological design (**Methods**). Proto’s graphical interface also parallels visual programming languages that enable users to specify logical operations without writing traditional code (Maloney et al., 2010).

While Proto is built around tools and models commonly available to the scientific community, greater usability and composability can themselves change how these tools and models are used. We developed Proto to enable safe and responsible research and education. By implementing an API and web application for usability, we are also able to monitor usage that goes outside of these boundaries, and Proto’s hosted agentic interface includes safeguards to reduce the risk of misuse, in addition to being built on top of LLMs that have their own biological safeguards. As biological AI improves, safety practices will evolve alongside it, which will include usage monitoring and updating safeguards as necessary as Proto continues to evolve (Carter et al., 2023).

The requirement to test Proto-generated designs experimentally remains an important bottleneck and caveat underlying all of Proto’s generations. As such, Proto is complementary to existing methods commonly used in biological engineering, such as large-scale screening or directed evolution. Further improvements in technologies for building and testing biological designs at the scale of multi-gene systems or complete genomes will be required to test the most complex Proto programs (Robinson et al., 2026).

We anticipate that improvements in generative models of biological sequences, predictive sequence-to-function models, and AI agents built on reasoning large language models will increase the scope, scale, and reliability of biological functions that are designable by Proto. Future improvements will also arise from developing more efficient optimizers beyond simulated annealing or gradient descent. As technologies for generative biological programming like Proto continue to mature, we envision a future in which biological design is limited not by natural parts lists but by human creativity.

## 4. Methods

### 4.1. Theoretical and methodological preliminaries

When designing a language for generative biology, we had the following desiderata:

**Generality.** Any unifying framework must accommodate a diversity of biological design tasks, modalities (DNA, RNA, protein), generative models (autoregressive language models, diffusion models), and scoring functions (neural-network predictors, biophysical models, simple heuristics, experimental measurements).
**Compositionality.** Complex biological systems typically involve interacting components and multiple design objectives. A desirable framework must allow simple primitives to be combined into larger programs without compounding the algorithmic complexity.
**Efficiency.** The time to generate biologically relevant sequences must be tractable, especially given the combinatorially large theoretical search space that makes naive enumeration of all possible sequences impractical.
**Design success.** Ultimately, generated designs must actually be functional, ideally after experimentally testing only tens of designs, without large-scale screening. This requires the ability to combine rich generative priors (so candidate sequences are biologically plausible) and strong scoring functions (so candidates have the desired function) (King et al., 2025; Merchant et al., 2025; Mille-Fragoso et al., 2025; Pacesa et al., 2025).

To realize these criteria, it is helpful to formalize aspects of biological design within a theoretical framework.

Fundamentally, biological designs are encoded by sequences. We denote a sequence as *x* ∈ X^*L*^, where *X* is the vocabulary (typically the four DNA bases, four RNA bases, or twenty canonical amino acids) and L is the sequence length.

We also need to encode to what extent a sequence is desirable. A highly flexible way of doing so is by specifying a constraint function *f* : X^*L*^ → ℝ that maps a discrete sequence to a scalar score. More generally, we can also admit a constraint function that is defined over K sequences

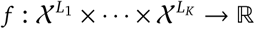

and maps these sequences to a scalar score. In this study, we take the convention that lower scores are more desirable.

We also need a way of producing sequences. Formally, we can define sequence production as sampling from a probability distribution *ρ* over ***χ***^*L*^. A central observation is that any function from sequences to scalars can be turned into a probability distribution. Specifically, given an arbitrary scoring function *E*(*x*), the distribution

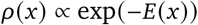

is well-defined (up to a partition function) and concentrates on sequences with low *E*. This is the energy-based model formulation (LeCun et al., 2006), and it provides a bridge from constraints (scalar functions of sequences) to distributions (objects from which we can sample). Therefore, the EBM formulation helps convert a scoring function into a generative model.

The EBM formulation also makes constraints naturally compositional (Du and Kaelbling, 2024). For example, suppose a user has three constraints over four sequences *x*_1_, *x*_2_, *x*_3_, *x*_4_, where *f*_*A*_ acts on *x*_1_, *f*_*B*_ acts on *x*_2_, and *f*_*C*_ acts on (*x*_3_, *x*_4_). Each constraint individually defines an EBM:

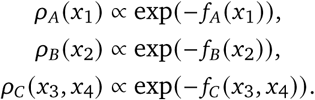

Treating these three distributions as independent factors of a joint distribution—that is, taking a product of experts (Hinton, 2002)—the joint distribution over all four sequences is

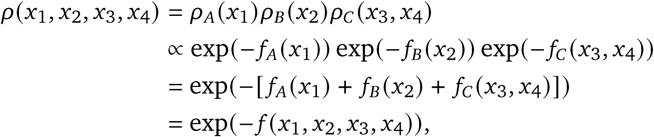

where *f* = *f*_*A*_ + *f*_*B*_ + *f*_*C*_. The combined distribution is itself an EBM, with energy equal to the sum of the individual energies. More generally, note that any collection of constraints 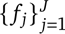 acting on (possibly overlapping) subsets of program sequences combines into an aggregate constraint

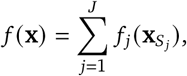

where **x**_*S*_*j*__ denotes the sequences on which *f* _*j*_ acts.

### 4.2. Generative sequence priors and design targets

A remaining consideration is how to efficiently sample from this distribution. A naive Metropolis-Hastings (MH) sampler with a uniform proposal (i.e., randomly sampling a single mutation uniformly over the vocabulary and uniformly over the sequence positions on each step of the optimization loop) would explore sequence space too slowly to be useful, especially under a complex set of constraints, as typically most of X^*L*^ has negligible mass under a biologically meaningful *ρ*. As noted by Du and Kaelbling (2024) and as done in many biological design campaigns in practice (Hayes et al., 2025; King et al., 2025; Merchant et al., 2025; Nguyen et al., 2024; Ruffolo et al., 2025), we can leverage unsupervised generative models trained on large collections of biological sequences (and optionally conditioned on some property, such as a protein backbone) to serve as proposal distributions. We will denote such a generative model as *p*(*x*). Not only do these models admit efficient sampling, but they also serve as powerful biological priors that substantially improve design success rates upon experimental testing (Brixi et al., 2026; Dauparas et al., 2022; Hie et al., 2024; Ingraham et al., 2023; Madani et al., 2023).

Then, assuming a single sequence *x*, a generator distribution *p*(*x*), and a constraint *f* (*x*), we can now define the design target as

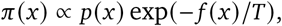

where the prior *p*(*x*) acts as a base distribution over biologically plausible sequences, the constraint *f* (*x*) essentially acts as a negative log-likelihood favoring designs that satisfy the criteria of interest, and the temperature *T* > 0 controls the trade-off between fidelity to *p* and minimization of *f*. As *T* → 0, samples from *π* concentrate on the modes of *p* that simultaneously minimize *f* (as *T* → 0, the − *f* (*x*)/*T* term dominates, so mass concentrates on the minimizers of *f*, with the prior *p* breaking ties among them). Various optimization algorithms can be used to obtain these samples.

Note that when producing multiple sequences, generators can also naturally compose. A program with *K* generators 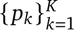, each acting on disjoint subsets of sequences, induces a factorized prior

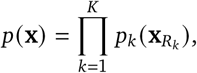

where **x** denotes the full set of design sequences and **x**_*R*_*k*__ denotes the subset proposed by *p*_*k*_. This factorization enables modular sampling, as most generative models in biology operate over a single modality, allowing complex programs to naturally combine multiple generators.

### 4.3. Proto: Full factorization, graphical representation, and interface

All of these components form the primitives used in the Proto design language. Proto supports genetically encoded sequences, i.e., canonical DNA, RNA, and proteins. Functions that encode the desirability of a set of sequences are referred to as constraints. In Proto, a constraint may correspond to the output of a neural-network predictor (e.g., AlphaFold pLDDT, Borzoi track prediction) or a simple statistic (e.g., coding density, GC content). Models that sample new sequences are referred to as generators. Generators can be implemented by large autoregressive neural networks, or simpler methods such as uniform sampling of sequence space. Finally, algorithms for sampling from a design target distribution *π* are referred to as optimizers, which we discuss at length in the next section.

Combining the constraint and generator decompositions into our EBM formulation, the full design target for a program with a set of sequences *x*, *K* generators, and *J* constraints is

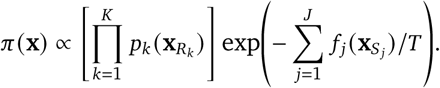

**x**_*Rk*_ denotes the subset of sequences proposed by *p*_*k*_ and **x**_*Sj*_ denotes the subset of scored sequences provided as input to the scoring function *f* _*j*_, where these can be any subset of **x**.

The factorized structure of *π* admits a natural graphical representation as a factor graph (Kschischang et al., 2001), a bipartite graphical model in which one set of nodes corresponds to sequence variables {*x*_*i*_} and another to factors *p*_*k*_ and *f* _*j*_, with edges connecting each factor to the sequences on which it depends (**Figure S1A,B**). The factor graph captures the full structure of a Proto program where each factor node corresponds to a generator or a constraint, each variable node corresponds to a designed sequence, and edges encode the dependencies that determine how generation and scoring couple across the program. These factor graphs enable a principled, graphical representation of the components of a generative design campaign.

The user interface for Proto is largely based on the factor graph representation (**Figure S1C**), while also taking inspiration from visual programming languages that enable adding and connecting core components within a graphical environment (for example, in a web application). The Python API for Proto directly enforces abstractions at the level of constraints, generators, and optimizers. These abstractions relate to each other based on the connectivity of the factor graph; generators produce sequences (represented as segments), constraints score sequences, and the optimizer controls sequence sampling based on the full set of generators and constraints.

The distributional structure encoded by Proto is a hallmark of probabilistic programming languages, which separate the specification of a probabilistic model from the inference procedure used to query it (van de Meent et al., 2021). In a typical probabilistic programming language, sampling statements define a generative prior while observe or factor statements re-weight that prior by likelihood or soft-evidence terms, after which a generic inference engine (for example, MCMC or variational inference) approximates the induced posterior. Proto instantiates this same separation for biological design in which generators play the role of sampling statements that define the prior *p*(**x**), each constraint plays the role of a factor statement contributing a log-density term − *f* _*j*_ (**x**_*Sj*_)/*T*, and optimizers play the role of the inference engine that queries the resulting target *π*(**x**). Proto further specializes generic probabilistic programming in two ways. First, rather than user-specified parametric priors, Proto’s generators are typically large pretrained neural models that supply strong, learned priors over biologically plausible sequences. Second, because the objective of design is to obtain high-scoring sequences rather than to characterize uncertainty, Proto’s optimizers seek the modes of *π* (via simulated annealing, gradient descent, or related procedures) rather than producing samples representative of the full distribution.

### 4.4. Optimization with Metropolis-Hastings and other algorithms

An important remaining consideration is the kind of algorithm used to sample from *π*. As alluded to above, in the most general case, we can use a Metropolis-Hastings (MH) algorithm in which, at each step, we (i) select a generator *p*_*k*_ (e.g., uniformly at random, or in a fixed cycle), (ii) sample a candidate update from *p*_*k*_ while holding the remaining sequences fixed, and (iii) accept the candidate with probability determined by the change in the aggregate constraint. Because the proposal is drawn from a factor of the prior, the prior contribution to the MH ratio cancels. To see this, let **x** denote the current state and **x**^′^ the proposed state, with proposal density 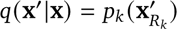 for the updated sequences. The standard MH ratio,

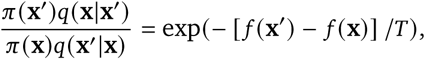

reduces to the constraint-only form because the *p*_*k*_ factor in *π* cancels against the proposal *q*, and all other prior factors are identical in numerator and denominator (their sequences are unchanged). The acceptance probability is therefore

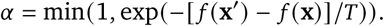

In practice, only those constraints whose value depends on the updated sequences contribute to *f* (**x**^′^) − *f* (**x**), so the per-step cost of evaluation is bounded by the local connectivity of the factor graph rather than by the total number of constraints in the program. An example algorithm is provided below:

#### Algorithm 1 Illustrative optimization with Metropolis-Hastings MCMC

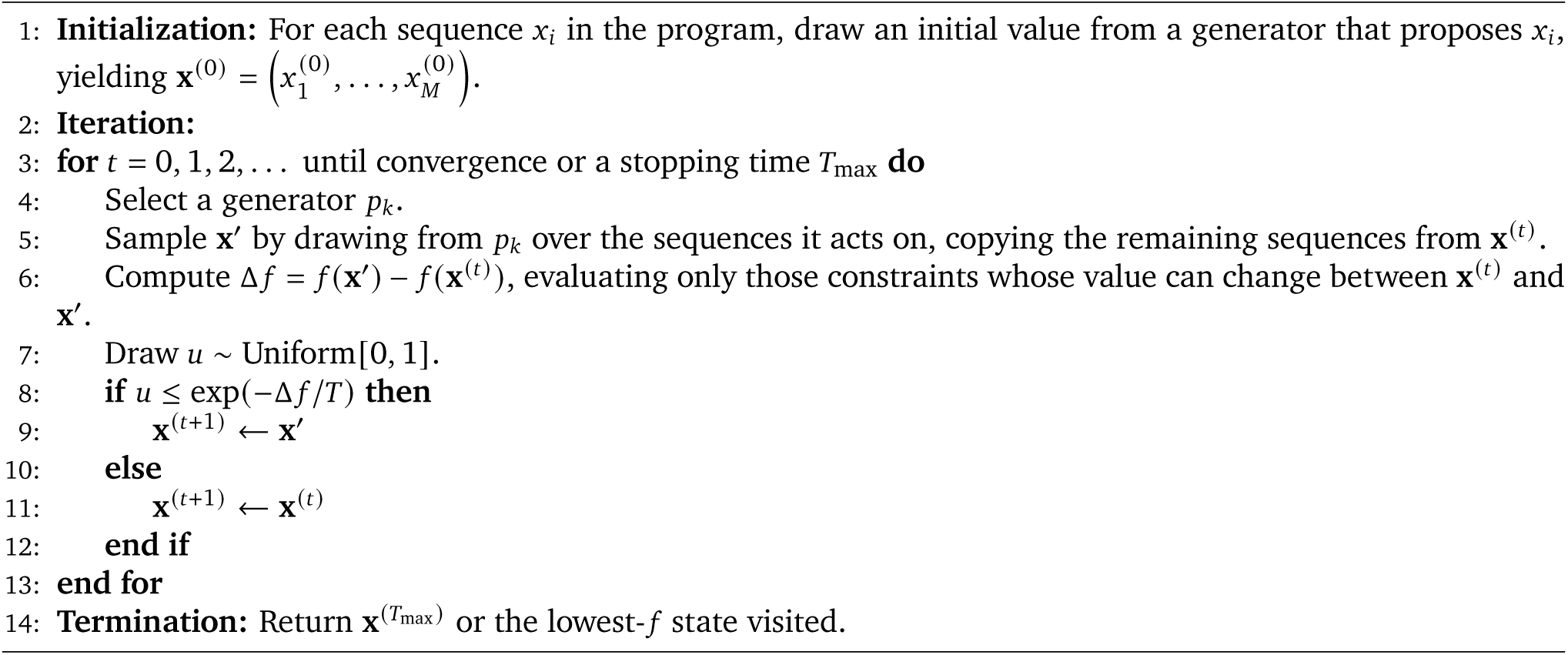

In the *T* → 0 limit the procedure reduces to greedy energy descent. Proto also supports (and defaults to) simulated-annealing schedules in which *T* decreases over the course of optimization.

While the MH algorithm is the most general optimizer in Proto (and also serves as a useful explanatory example), certain program structures can admit more efficient alternatives than MCMC. Notably, all of these sampling algorithms are only meant to approximately sample from the target distribution *π* and are not guaranteed to produce the globally minimal sequences under the aggregate constraint function. However, these approximations are often sufficient to obtain good design results in practice.

We describe the currently implemented optimizers aside from MCMC below (**Figure S4**), but note that the language can flexibly accommodate future algorithms as needed.

#### Rejection sampling optimizer

When the prior *p*(**x**) is sufficiently informed and constraints take the form of bioinformatic filters, rejection sampling (i.e., drawing from *p* and discarding any sequence for which *f* exceeds a threshold) is an effective and embarrassingly parallel optimization strategy. This is the optimizer used to filter candidate CRISPR-Cas operons generated by fine-tuned Evo 1 (Nguyen et al., 2024).

#### Gradient-based optimizer

When all constraints are differentiable with respect to a continuous relaxation of the sequence (e.g., logits or one-hot embeddings), gradients of *f* can be backpropagated through the constraint stack to update the sequence directly. This is particularly efficient when the program contains a small number of high-information constraints, e.g., AlphaFold-based scores in the Germinal antibody design protocol. Proto supports both straight-through estimators for discrete tokens and continuous relaxations followed by argmax decoding.

#### Cycling optimizer

When two generators or generator-constraint pairs alternate (e.g., a structure predictor and a structure-conditioned autoregressive sequence model), the optimizer can cycle deterministically between them. This is the strategy used in the Protein Hunter protocol (Cho et al., 2025), which alternates Boltz-2 structure prediction with ProteinMPNN sequence redesign for efficient *de novo* sequence-structure codesign.

#### Beam search optimizer

When the dominant generator is autoregressive and constraints can be evaluated on partial sequences, beam search retains the top-few prefixes by aggregate constraint at each decoding step and extends only those prefixes, with constraint scores accumulating along each beam. This is the optimizer used in the Morse-code chromatin-accessibility design (Brixi et al., 2026), in which Evo 2 decodes DNA autoregressively while Enformer and Borzoi score the resulting accessibility profile.

In each case, the optimizer’s choice is driven by the structure of the program, including which generators and constraints are used, whether constraints decompose along sequence prefixes, and whether constraints are differentiable with respect to the input. The four primitives and the EBM target distribution *π* remain unchanged across optimizers, only the procedure for (approximately) sampling from *π* varies.

### 4.5. Proto tools Python library

The core functionality of Proto depends on the efficient composition of the large and growing collection of programs and models that the computational biology community has developed to generate, score, and select biological sequences. These tools are typically released as independent artifacts, each requiring its own virtual environment on top of frameworks such as PyTorch and JAX that carry their own, frequently incompatible, dependencies. Deploying several such tools within a single codebase is therefore difficult as their dependencies often conflict and cannot be co-installed in a shared environment. The Proto tools python library resolves these conflicts by installing each wrapped tool in its own isolated environment while exposing every tool through a single, uniform interface, so that end users can compose heterogeneous models without managing installation, dependency resolution, or environment configuration themselves.

Each tool is wrapped as a self-contained module that declares its software dependencies as well as any required model weights and references databases. The underlying infrastructure of the library then provisions these resources automatically upon the first use of the tool, and caches them for reuse across subsequent tool calls. Every call to a given tool spawns a subprocess that is executed within that tool’s dedicated environment, with inputs and outputs passed through a standardized data-exchange format so that sequences, structures, and metrics can move between tools irrespective of their native input/output conventions.

Beyond environment isolation, the library manages accelerator hardware on behalf of the user so that design campaigns composing many models can run efficiently without manual scheduling. A device manager tracks every model allocation, places models onto free GPUs, and offloads models when all devices are occupied. For batched workloads, the library fans computation out across any available devices, and multiple models can optionally be packed onto a single device as space permits. Partitioning uses cost-aware scheduling based on a per-tool cost estimate so that workers with heterogeneously sized inputs finish at approximately the same time, and identical inputs are deduplicated so that repeated items are computed only once.

Through this execution layer, the Proto tools library provides standardized access to over 120 tools spanning generative sequence models (such as ESM3, ProGen3, Evo 2) (Hayes et al., 2025; Bhatnagar et al., 2025; Brixi et al., 2026), inverse-folding models (ProteinMPNN, LigandMPNN) (Dauparas et al., 2022, 2025), structure predictors (ESMFold, Boltz-2, Chai-1, and AlphaFold 3) (Lin et al., 2023; Passaro et al., 2025; Chai Discovery et al., 2024), sequence-to-function models (AlphaGenome, Borzoi, Enformer, Malinois, and SpliceTransformer) (Avsec et al., 2026, 2021; You et al., 2024; Linder et al., 2025; Gosai et al., 2024), antibody and protein language models (AbLang, ESM2) (Olsen et al., 2022; Lin et al., 2023), structure design models (RFdiffusion3) (Butcher et al., 2025), and standard bioinformatic utilities (such as ORFipy, Prodigal, PyHMMER, MMseqs2, BLAST, and MAFFT) (Hyatt et al., 2010; Steinegger and Söding, 2017; Cock et al., 2015; Katoh et al., 2002; Larralde and Zeller, 2023; Singh and Wurtele, 2021). We release the Proto tools library as open-source software on GitHub under an MIT license, enabling researchers to install and compose this body of tools with minimal configuration.

### 4.6. Proto language Python library

The Proto language abstractions are implemented as core classes in a Python library for expressing and optimizing a biological design. The fundamental unit of design is a Sequence, which wraps a biological string of a declared type (DNA, RNA, protein, or ligand) with type and character validation and stores a metadata record that tracks constraint scores and optimization history. Sequences are organized and stored within Segments, which are contiguous regions (e.g., a promoter or coding sequence) that are populated and managed during optimization. An ordered collection of Segments forms a Construct, which represents a complete biological design (e.g., a gene composed of a promoter, ribosome-binding site, coding sequence, and terminator). Together, the Sequence, Segment, and Construct constitute the scaffold of a design on which generators, constraints, and optimizers operate.

Optimization is driven by three operators that act on Segments. A Generator supplies the proposal distribution for populating a Segment through a unified interface, so that random samplers, masked and autoregressive sequence models, structure-conditioned inverse-folding models, and diffusion-based backbone generators can be used interchangeably. A Constraint scores how well a candidate satisfies a design objective, returning a weighted scalar by default or an accept/reject filter when given a threshold, and may additionally expose a differentiable backward pass that supplies gradients for gradient-based optimization. Constraints can be aggregated to define an arbitrary overall objective function. An Optimizer is the search strategy that drives a design loop. Each Segment maintains two pools of Sequences: the proposals being explored and the results retained as the best found so far. At each iteration the Generator fills the proposal pool, the Constraints score those candidates, and the Optimizer updates the result pool to retain the lowest-energy candidates, which seed the next iteration. A Program chains Optimizers into sequential stages over shared Constructs, with the outputs of each stage seeding the next, so that different generators, constraints, and optimization strategies can act on a single design in succession.

### 4.7. Proto cloud API server

The backend API server was implemented in Python using the FastAPI framework and connected to a serverless cloud platform for remote computation. Proto Programs are expressed in JSON format for standardized data serialization and pipelining. Program optimizations and model computations are dispatched by the API server to the serverless cloud platform with an isolated environment and the necessary compute resources. Per-iteration results are streamed to the user interface so scoring trajectories and intermediate designs are surfaced as they are produced. State and metadata are persisted in a cloud database for persistent storage.

### 4.8. Proto user interface

The user interface was implemented using React and TypeScript within the Next.js framework to visualize and compose Proto primitives. The interactive drag-and-drop canvas can be used to assemble segments, constraints, and generators through connectable nodes to create complete visual programs. These primitives can be composed into multi-stage optimization pipelines and dispatched to cloud infrastructure for execution. Results are streamed in real time, with score trajectories plotted across optimization iterations as they are produced. Every node exposes typed inputs and parameters that are customizable for a design objective and directly maps to the inputs and parameters provided to the Python API. Completed designs produce interactive sequence and structure viewers and per-component energy score breakdowns. The user interface includes a chatbot assistant that educates and guides users through Proto components and composing programs.

### 4.9. Symmetric homo-oligomer design

Symmetric homo-oligomeric protein backbones were designed by Metropolis-Hastings simulated annealing over a single protomer amino acid sequence, scored against an ESMFold-derived objective that simultaneously rewards monomer fold quality and oligomer-level ring symmetry and compactness (Lin et al., 2023). At every step of the chain, the current protomer sequence was replicated *in silico* into N identical copies to form a candidate homo-N-mer; the resulting complex was folded with ESMFold; and four scalar terms (oligomer pLDDT, pTM, ring-symmetry deviation, and globularity) were combined into a single energy that the chain sought to minimize.

The optimizer was a single-trajectory Metropolis-Hastings chain run for a fixed budget of *M* steps with one proposal per step and exponential temperature annealing from *T*_max_ = 1.0 to *T*_min_ = 1 × 10^−4^, with the temperature at step *k* given by *T* (*k*) = *T*_max_ · (*T*_min_/*T*_max_)^(*k*−1)/(*M*−1)^. At each step, a single amino acid substitution in the protomer was proposed drawn uniformly over the 20-amino-acid alphabet and uniformly over the sequence positions and the proposal was accepted with probability *α* = min(1, exp(−(*E*_new_ − *E*_old_)/*T*)).

The energy minimized by the chain was a weighted sum of four ESMFold-based terms. The first two are based on predicted confidence as 1 − the mean per-residue pLDDT and a global-quality term defined as 1 − pTM. For the ring-symmetry term, the protomer is duplicated N times, the resulting complex is folded with ESMFold, one centroid is computed per chain from backbone coordinates, and the standard deviation *σ* of all pairwise inter-centroid distances is computed and clipped as min(1.0, *σ*/10.0) so that lower values correspond to more symmetric oligomers. For the globularity term, the same replicated complex is folded, the standard deviation *g* of distances from every backbone atom to the global complex centroid is computed, and the term is clipped as min(1.0, *g*/20.0) so that lower values correspond to more compact oligomers. The total energy was computed as

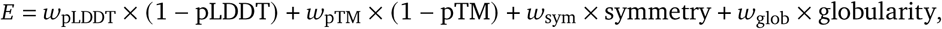

with respective weights (*w*_pLDDT_, *w*_pTM_, *w*_sym_, *w*_glob_) = (1, 1, 1, 5). ESMFold for multimer scoring was run with 25-residue poly-glycine linker inserted between concatenated chains and a residue-index offset of 512 separating each new protein chain.

In our design sweep, protomer length was swept across 50 and 100 residues, oligomer order across *N* = 2 to 8, and chain length across step budgets of 5,000 and 10,000 steps, defining 28 (length, *N*, steps) cells that were each populated by a small number (∼10) of jobs. Designs were scored based on their final ESMFold pTM and pLDDT scores.

### 4.10. Protein Hunter monomer design

Protein Hunter (Cho et al., 2025) is a *de novo* single-chain protein design loop that alternates between structure prediction and backbone-conditioned protein sequence design. Each design trajectory was initialized as a single chain of fixed length *L* (of 200, 300, or 400 residues, depending on the sweep) seeded with an all-unknown sequence, that is, *L* copies of the wildcard residue `X`. The structure was then iteratively refined for a fixed number of cycles. At each cycle, the current sequence was folded with the chosen structure-prediction model (Boltz-2 or Chai-1) (Passaro et al., 2025; Chai Discovery et al., 2024), the predicted backbone was passed to ProteinMPNN to sample a single new sequence (Dauparas et al., 2022), and the sampled sequence replaced the current one and was carried forward into the next cycle. After the final cycle, one additional structure prediction was performed on the final sequence.

Sequence resampling at each cycle was performed with ProteinMPNN with sampling temperature 0.1 and cysteine exclusion. A single sequence was sampled per cycle. Structure prediction at each cycle was performed by either Boltz-2 or Chai-1. For each predicted model we recorded pLDDT and pTM indexed by cycle, where cycle 0 corresponds to the initial structure prediction on the all-`X` seed, and cycle *N* = 5 corresponds to the final structure prediction on the final designed sequence. Each structure predictor was run across design lengths of 200, 300 and 400 residues with MSAs disabled and 5 design cycles, with 20 independent trajectories per (backend, length) cell, for a total of 120 trajectories.

### 4.11. CRISPR-Cas generation

Genomic loci were generated with the fine-tuned Evo 1 genome language model using the evo-1-8k-crispr checkpoint. Each autoregressive generation had a two-token prompt consisting of the Cas9 class token followed by a single nucleotide (“‘`A`”) and was extended to a fixed length of 8,000 nt. Sampling was performed across a 3 × 2 grid of decoding hyperparameters comprising sampling temperatures of 0.1, 0.3 and 0.5 and top-*k* values of 2 and 4, with top-*p* fixed at 1.0. Two thousand sequences were drawn per (temperature, top-*k*) combination, yielding 12,000 candidate loci per sampling job. The full design campaign comprised four independent sampling jobs and therefore 48,000 candidate loci in total.

Each generated locus was passed through an eight-stage cascade of orthogonal filters applied in order of increasing computational cost such that failure at any stage short-circuited subsequent evaluation. First, open reading frames were predicted on both strands with ORFipy under a minimum length of 3,000 nt; the longest predicted ORF on either strand was translated and retained as the candidate Cas9 protein for all downstream protein-level analyses (Singh and Wurtele, 2021). Second, the translated protein was searched against a Cas9 profile HMM using PyHMMER’s hmmsearch, requiring at least one hit at sequence-level *E* ≤ 1 × 10^−3^. This profile HMM was built using Cas9 sequences from the fine-tuning dataset (Larralde and Zeller, 2023). Third, the locus was scanned for CRISPR arrays with MinCED requiring at least three repeats of length ≥ 23 nt (Bland et al., 2007). Fourth, protein identity to the training distribution was assessed by MMseqs2 search (using default mmseqs easy-search wrapper params) of the candidate against a combined reference database of 130,972 Cas-family training proteins; candidates were required either to return no hit or to have a top-hit protein sequence identity below 90% (Steinegger and Söding, 2017). Fifth, the candidate was pairwise-aligned to its nearest training-set hit with MAFFT, and the Gini coefficient of the gap distribution along the alignment was required to be below 0.1, which suppresses candidates whose nearest match is dominated by a small number of long indel blocks (Katoh et al., 2002). Sixth, domain architecture was verified by a PyHMMER scan against a curated Cas9 domain HMM library at domain-level *E* ≤ 1×10^−10^, with all four canonical Cas9 domains (RuvC_1, RuvC_2, RuvC_3 and HNH) required to be detected (Larralde and Zeller, 2023). Seventh, tracrRNA association was tested with CRISPRtracrRNA in type-II mode (`model_type=“I”`), requiring both a predicted tracrRNA and an IntaRNA-supported anti-repeat interaction (Mitrofanov et al., 2022). Eighth, three-dimensional structural plausibility of the translated protein was assessed by AF3 monomer prediction with multiple sequence alignments enabled using a locally installed ColabFold MSA pipeline, and the resulting model was required to satisfy mean pLDDT ≥ 75.0, radius of gyration < 45.0 Å and longest *α*-helix < 50 residues (Abramson et al., 2024). The four sampling runs in our sweep returned 8, 11, 8 and 13 candidates, for a total of 40 passing designs from 48,000 sampled loci.

### 4.12. Chromatin accessibility beam search

DNA sequences were designed by autoregressive beam search over a fixed genomic locus consisting of a left genomic flank, a variable target segment of length 21,120 bp, and a fixed right genomic flank. Following Brixi et al. (2026), generated DNA sequences replace the native sequence at chrX: 52,051,929-52,123,468 in the mm39 mouse genome, with surrounding context given as flanks. Both flanks were held constant throughout optimization; only the target segment was modified by the design procedure. The target encoded a Morse-code accessibility pattern (“PROTO” or “`.--. .-.`”) in which dot and dash symbols mark positions intended to have high accessibility, separated by gap regions intended to remain inaccessible. Dots were 384 bp wide, dashes were 1,152 bp wide, intra-letter gaps were 384 bp, and inter-letter gaps were 1,152 bp; the first symbol began at position 0 of the target window, and under these settings the full pattern spanned the entire 21,120-bp target. The autoregressive prompt for Evo 2 was taken as the final 8,192 bp of the left flank.

At beam search step t, scoring was only performed on the partially designed sequence. Beam search maintained *K* = 2 active beams. At each iteration, every beam generated *N* = 18 candidate continuations of length *L* = 128 bp, yielding *K* × *N* = 36 candidate children per beam-search step. Because the target length was 21,120 bp and each step added 128 bp, optimization proceeded for exactly 21,120 / 128 = 165 beam steps. Evo 2 sampling used temperature 1.0, top-*k* = 4 and top-*p* = 1.0. Generation was performed with key-value caching enabled to enable efficient sampling. All *K* beams were initialized from the same Evo 2 prompt. At each beam step, candidate proposals were generated independently from each parent beam, scored under the accessibility objective, and ranked across the union of all *K*×*N* children. The top *K* children were retained as parents for the next step. Pruning was performed using the score from only the most recent step. Importantly, candidate scoring was performed on the full accumulated prefix, not on the most recent 128-bp continuation in isolation.

Each candidate prefix was independently scored by two accessibility predictors, Borzoi and Enformer, and the total beam energy was the sum of the two model losses (Linder et al., 2025; Avsec et al., 2021). For Borzoi, the model input window was 524,288 bp and predictions were evaluated at 32-bp resolution; for Enformer, the input window was 196,608 bp and predictions were evaluated at 128-bp resolution. In both cases, the model window was extracted from the full construct so that the designed target remained aligned to its intended genomic coordinates. When required, the right flank was truncated once at setup to the minimum length needed to satisfy both model windows.

Borzoi and Enformer designs used the DNase hypersensitivity tracks in 129 ES-E14 cells. Borzoi was run as a four-replicate ensemble and reduced to a single per-bin track by lower confidence bound, defined bin-wise as the mean minus the standard deviation across replicates; Enformer outputs were reduced by mean across the selected tracks. The total accessibility objective was

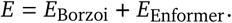

The base objective for each model was a pattern-matching ℓ_1_ loss. The reduced model output was first restricted to bins overlapping the currently generated target prefix and then normalized by the global maximum of the model output across the full input window, yielding a normalized signal *s̃*_*m*,*i*_ for model *m* and bin *i*. A binary Morse target *y*_*i*_ ∈ {0, 1} was constructed at each model’s native output resolution by setting bins that overlapped any dot or dash window to 1 and all other bins to 0; even partial overlap with a single positive window was sufficient to mark a bin as positive. The base loss for model *m* was

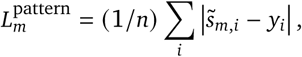

where the sum runs over the *n* bins covered by the currently generated prefix. No loss term was applied to unsampled positions of the target.

To shape candidate ranking beyond bin-wise matching, the pattern loss for each model was supplemented with four auxiliary terms reflecting global properties of the predicted accessibility signal. Let *s̅*_high_ and *s̅*_low_ denote the mean raw (un-normalized) signal across Morse high windows and Morse low windows respectively, *a* denote the raw amplitude max(*s*) − min(*s*) over the target region, *T* (·) = log1p(·) denote an elementwise transform, and *δ* denote a contrast margin. The per-model loss was

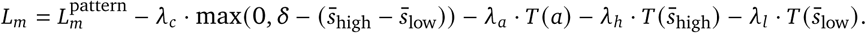

We used *δ* = 0.2, contrast weight *λ*_*c*_ = 1.0, raw-amplitude weight *λ*_*a*_ = 0.2 with the amplitude evaluated over the target region, high-window weight *λ*_ℎ_ = 0.4, and low-window weight *λ*_*l*_ = 0.4, with log1p applied to the amplitude and to both window means. Lower values of *L*_*m*_ remained preferable.

### 4.13. *De novo* antibody CDR design with Germinal

We reimplemented Germinal (Mille-Fragoso et al., 2025), a gradient-based pipeline for *de novo* design of antibody complementarity-determining regions (CDRs), to generate binders against programmed death-ligand 1 (PD-L1) in both single-domain (VHH) and single-chain variable fragment (scFv) backbone formats. In Proto, the antibody hallucination procedure that proposes CDR sequences serves as the generator, AlphaFold 2 (AF2) confidence and antibody-antigen interface geometry together with the AbLang antibody language model serve as constraints (Olsen et al., 2022; Jumper et al., 2021), and gradient descent over a continuous sequence relaxation followed by semi-greedy Markov chain Monte Carlo (MCMC) serves as the optimizer. The PD-L1 target was held fixed as chain A of a 115-residue antigen derived from the PD-L1–nanobody complex (RCSB PDB 5JDS, residues 18–132 renumbered to 1–115), with interface hotspot residues A37, A39, A41, A96, and A98, while only the CDR positions of the binder were allowed to vary. For VHH designs, the CDR positions (zero-indexed) were 25-35, 53-60, and 99-116 within a 131-residue nanobody scaffold; for scFv designs, the CDR positions were 25-32, 50-57, 96-108, 150-155, 173-178, and 212-220 within a 242-residue VH-linker-VL scaffold, both taken from the Germinal repository (https://github.com/SantiagoMille/germinal).

Each design trajectory proceeded through a staged optimization schedule. AF2 was first used to hallucinate CDR sequences by backpropagating an antibody-antigen design loss into a continuous representation of the binder sequence, beginning with 60 logit-space optimization steps, followed by 35 steps over a softmax-relaxed sequence, and concluding with 10 steps of semi-greedy MCMC; three structural recycles were used per AF2 evaluation. Confidence gates were applied between stages, first on pLDDT and ipTM and subsequently incorporating the interface predicted aligned error (ipAE), so that low-confidence trajectories were pruned before incurring the cost of later stages.

The AF2 hallucination loss was a weighted sum of structural and interface terms, comprising pLDDT (weight 1.0), interface pLDDT (1.0), intra-chain predicted aligned error (0.1), inter-chain predicted aligned error (0.5), ipTM (0.7), intra-chain contacts (0.1), inter-chain contacts (0.2), radius of gyration (0.1), helix content (0.1), *β*-strand content (0.1), and a distogram categorical cross-entropy term (0.01). Intra-chain contacts were defined with a 14.0 Å cutoff and a target of two contacts, and inter-chain contacts with a 20.0 Å cutoff and a target of ten contacts. To bias hallucinated CDRs toward natural antibody sequence likelihood, the AbLang antibody language model was applied with a sampling temperature of 0.1 and a redesign bias weight of 10.0, and search-stage proposals used a per-residue mutation rate of 0.05.

Trajectories that passed the confidence gates were redesigned and filtered against orthogonal structure-prediction oracles. Each candidate passed through hallucinated-structure checks, an external cofold of the antibody-antigen complex followed by FastRelax refinement in PyRosetta, and an initial set of Germinal filters. CDR positions were then resampled with the antibody-specific inverse-folding model AbMPNN (40 samples per design at sampling temperature 0.2, retaining the top five candidates for VHH designs and the top four for scFv designs), and each redesigned candidate was cofolded a final time with AF3, relaxed, and scored by ipSAE and pDockQ2 (Basu and Wallner, 2016; Dreyer et al., 2023). A single live MMseqs2/ColabFold homology search was performed per campaign to construct a target-only multiple sequence alignment that was reused across cofolding (Ovchinnikov et al., 2025). Final acceptance required pDockQ2 > 0.23 and ipSAE ≥ 0.6, together with external structure-prediction metrics of pLDDT ≥ 0.8, ipTM ≥ 0.75, pTM ≥ 0.8, and predicted aligned error < 8 Å. Confidence thresholds during hallucination differed by backbone format; VHH trajectories required pLDDT ≥ 0.75, ipTM ≥ 0.68, and ipAE ≤ 0.30, whereas scFv trajectories required pLDDT ≥ 0.80, ipTM ≥ 0.67, and ipAE ≤ 0.28. The VHH scaffold was parameterized with framework lengths of 25, 17, 38, and 14 residues and CDR lengths of 11, 8, and 18 residues; the scFv scaffold used VH framework lengths of 25, 17, 38, and 11 residues with VH CDR lengths of 8, 8, and 13, a 15-residue linker, and VL framework lengths of 26, 17, 33, and 10 residues with VL CDR lengths of 6, 6, and 9. With this program, we obtained PD-L1 binders bearing *de novo* CDRs in both VHH and scFv backbones that achieved high AF2 confidence metrics (pLDDT > 0.8, ipTM > 0.6).

### 4.14. Computational design of alternatively spliced introns

Introns with cell-line-specific alternative splicing were designed by Metropolis-Hastings simulated annealing over the internal intron sequence within a split fluorescent reporter. Each design embedded a candidate intron into a representative synthetic plasmid construct—specifically, an mScarlet-IRES-eGFP reporter at nucleotide position 477 (i.e., amino-acid position 159)—where a 1-kb target window centered on the intron was constructed for splice-site evaluation. The splice donor and acceptor dinucleotides were held fixed throughout optimization such that every intron began with GT and ended with AG, where the generator mutated only the internal intron core between these boundaries.

Two initialization strategies were used. The first strategy seeded each trajectory with a fixed 379-bp chimeric HBB intron 2 sequence (Nieuwenhuis et al., 2021), while the second strategy seeded each trajectory with a random DNA intron of either 151 or 301 bp carrying the fixed GT and AG boundaries. In all cases, candidate updates were proposed by a uniform nucleotide mutation generator, and each trajectory was optimized for 5,000 Metropolis-Hastings steps with one proposal per step and exponential temperature annealing from *T*_max_ = 1 × 10^−2^ to *T*_min_ = 1×10^−3^, retaining a single final sequence per trajectory. AlphaGenome and SpliceTransformer were both evaluated on GPU, with AlphaGenome restricted to positive-strand tracks (Avsec et al., 2026; You et al., 2024).

Each proposal was scored by a weighted sum of constraint scores. A SpliceTransformer boundary constraint evaluated the concatenated left flank, intron core, and right flank with 4-kb left and right genomic contexts, with the final score based on the predicted donor and acceptor probabilities at the fixed splice-site positions and combined as 1 − 0.5 × ( *p*_donor_ + *p*_acceptor_) to favor strong, correctly placed splice sites. The optimization optionally applied a SpliceTransformer tissue-specific splice-site usage penalty at the donor and acceptor positions, with the tissue channels following the AlphaGenome objective direction: the “BRAIN” channel for neural-proxy conditions, the “LIVER” channel for HepG2 conditions, and the “BLOOD” channel for K562 conditions.

Cell-line specificity was enforced primarily through AlphaGenome splice-site-usage (SSU) predictions evaluated in genomic safe-harbor contexts. For a target cell line the constraint was computed as 1 − SSU (a maximization objective), whereas for an off-target cell line the constraint was simply set equal to the SSU (a minimization objective). Introns were designed in both directions of each cell-pair comparison: on in neural and off in K562, off in neural and on in K562, on in HepG2 and off in K562, and off in HepG2 and on in K562. The following cell type ontology terms were used to index the AlphaGenome tracks: a neural proxy for SH-SY5Y (CL:0002319, CL:0011012, CL:0011020, CL:0000047), HepG2 (EFO:0001187), and K562 (EFO:0002067). All recovered constraint weights were unity (SpliceTransformer boundary, SpliceTransformer specificity, and the AlphaGenome target and off-target terms each weighted 1.0), so the aggregate energy minimized by the chain was

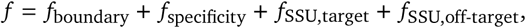

with the specificity term included only when splice specificity was enabled.

All designs were optimized under diverse genetic contexts such that splice-site predictions were averaged across delivery and integration contexts. The SpliceTransformer and AlphaGenome constraint functions leveraged plasmid contexts containing either a CMV, EF1*α*, or SFFV promoter. Additionally, 16-kb sequence context was used as input to AlphaGenome in which we further integrated this plasmid context into one of the AAVS1, CCR5, CLYBL, or hROSA26 genomic safe-harbor contexts during AlphaGenome scoring; the AlphaGenome input contained a 4-kb plasmid/gene left context + 1-kb SpliceTransformer target (described above) + 4-kb plasmid/gene right context, with this whole 9-kb construct centered within the 16-kb input sequence with the safe harbor sequence providing the remaining left- and right-flank sequences. The constraint scores were then summed across all relevant plasmid and genomic locus contexts with equal weight. In total, the focused sweeps yielded 2,872 designs across both directions of the neural-versus-K562 and HepG2-versus-K562 comparisons, spanning the chimeric HBB intron 2 and random initializations, 151-, 301-, and 379-bp intron lengths, and runs with and without SpliceTransformer specificity (**Table S1**).

### 4.15. Cloning of ProtoIntron reporter plasmid constructs

Sequences encoding generated ProtoIntrons, along with the right exon of mScarlet appropriate overhangs to facilitate integration into the reporter plasmid, were synthesized as eBlocks. To clone the synthesized introns into the reporter plasmid, parent plasmid pAM_BP_016_1, containing Ef1*α*-eGFP-CMV and a portion of the leftmost mScarlet exon, was digested with BstEII-HF (New England Biolabs) according to the manufacturer’s recommendations and subsequently gel purified to remove undigested product (Qiagen). Synthetic intron sequences were then inserted into the digested backbone using Gibson Assembly with NEBuilder HiFi DNA Assembly Master Mix (New England Biolabs) according to the manufacturer’s recommendations. Assembled products were then individually transformed into Stellar *E. coli* HST08 Competent Cells (Takara Bio), and positive clones were selected on LB agar plates containing 50 *μ*g/ml carbenicillin. Plasmid sequences were then confirmed by nanopore sequencing (Quintara Biosciences), before being miniprepped according to the manufacturer’s guidelines (Qiagen) for subsequent testing in mammalian cells.

### 4.16. Nucleofection of pooled intron retention reporter constructs

Designed intron reporters were initially tested as pooled libraries in K562, SH-SY5Y, and HepG2 cells. Constructs were grouped according to their design objective and pooled within library; for example, libraries designed for high retention in SH-SY5Y and low retention in K562 were nucleofected into both SH-SY5Y and K562 cells, whereas libraries designed for high retention in HepG2 and low retention in K562 were nucleofected into both HepG2 and K562 cells. For each cell line, 1 × 10^6^ cells were nucleofected with a pooled plasmid library containing 50 ng of each construct using the Lonza 4D-Nucleofector system and SF Cell Line Solution according to the manufacturer’s guidelines. Briefly, cells were collected, counted, and pelleted before being resuspended in SF nucleofection solution, mixed with the pooled plasmid library, and transferred to a Lonza 4D Nucleocuvette. Cells were nucleofected using cell-line-specific 4D-Nucleofector programs: K562, FF-120; SH-SY5Y, CA-137; and HepG2, EH-100. Immediately after nucleofection, cells were transferred to a 6-well plate containing 2 mL complete growth medium and cultured for 48 hours prior to RNA extraction. All nucleofections were repeated in triplicate, with each replicate coming from distinct subcultures.

### 4.17. ProtoIntron RNA extraction, library generation, and NGS

At 48 hours post-nucleofection, total RNA was extracted using the Zymo Quick-RNA Miniprep Kit (Zymo). For suspension K562 cultures, cells were collected by centrifugation, washed with PBS, and lysed directly in RNA lysis buffer. For adherent SH-SY5Y and HepG2 cultures, medium was removed, cells were washed with PBS, and RNA was extracted by direct lysis in the culture vessel. RNA was purified according to the manufacturer’s miniprep protocol. To reduce residual plasmid and genomic DNA, samples were initially treated with DNase during column cleanup. Eluted RNA was then subjected to a second DNase treatment using TURBO DNase (Thermo Fisher Scientific). RNA concentration was subsequently measured by Nanodrop and normalized across samples.

Normalized RNA was reverse transcribed using SuperScript IV reverse transcriptase (Thermo Fisher Scientific) and a library-specific reverse primer downstream of the construct barcode (primAM105) containing a 6-nt unique molecular identifier (UMI) according to the manufacturer’s guidelines. Resultant cDNA was then used as template for targeted amplification of the reporter region spanning the designed splice junction and construct-specific barcode. Initially, the number of PCR cycles was determined by qPCR of cDNA using SYBR Green Universal Master Mix (Applied Biosystems). Amplicons were generated using NEBNext High-Fidelity 2× PCR Master Mix using the following conditions: 98 °C for 30s for one cycle; 98 °C for 10s, 65 °C for 30s, and 72 °C for 45s for Ct + 3 cycles; and a final extension of 72 °C for 3 minutes. Amplicons were subsequently purified by double-sided SPRIselect bead size selection using a 0.5× right-side selection followed by a 0.9× left-side selection. Amplicon libraries were then quantified using a Qubit 1X dsDNA High Sensitivity Assay Kit (Thermo Fisher) before being sent to Quintara Biosciences for paired-end 2 × 300 bp sequencing using an Illumina NovaSeq X.

### 4.18. NGS data processing of splicing amplicon sequencing

Raw paired-end FASTQ files were processed to quantify splicing outcomes for each reporter construct. Construct-identifying barcodes and molecular UMIs were extracted from Read 2 (R2) using umi_tools (v1.1.6) with a regular expression pattern anchored to the invariant downstream constant sequence present in R2, extracting the 8-nucleotide construct barcode and the 6-nucleotide molecular UMI flanking this anchor. Barcodes were matched against a whitelist of known construct sequences, allowing a single mismatch only when the assignment was unambiguous; unmatched reads were discarded.

Splicing outcome was classified from Read 1 (R1) by locating a 22-nucleotide invariant anchor corresponding to the last 22 nucleotides of the upstream exon, ending immediately before the intron start. The 20-nucleotide window immediately downstream of this anchor was then matched against two reference sequences: the invariant downstream exon start sequence (correctly spliced, ≤2 mismatches) and the assigned construct’s designed intron sequence (intron-retained, ≤2 mismatches). Reads whose window sequence matched neither the canonical exon junction nor the start of the assigned construct’s intron and were spliced to a non-canonical acceptor site within the intron or downstream exon were classified as cryptically spliced. Reads in which the anchor was not found were discarded.

PCR duplicates were collapsed by grouping reads by their construct barcode and molecular UMI pair, with each unique molecule being assigned its splicing outcome by majority vote across all PCR copies. Chimeric contamination was handled in two steps. First, chimeric molecules whose R1 carried a different construct’s intron sequence were identified by intron mismatch and excluded from counting entirely. Second, to account for chimeric molecules that escaped this filter, namely those whose R1 carried correctly-spliced sequence from another construct, which is undetectable because all constructs share the same downstream exon, a per-sample chimeric rate *C* was estimated from the HBB2c positive control. As HBB2c is designed to splice constitutively, it should show ∼0% mis-splicing after the first exclusion step; any residual apparent mis-splicing is therefore attributed to chimeric contamination and used to estimate *C*. The corrected mis-spliced count was computed as:

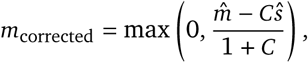

where *m*^ and ^*s* are the observed chimeric-excluded mis-spliced and correctly-spliced molecule counts, respectively (X. D. Chen et al., 2025). The correctly-spliced count was left unchanged.

Differential mis-splicing between target cell lines (SH-SY5Y or HepG2) and matched K562 controls was assessed by Fisher’s exact test on pooled corrected mis-spliced and correctly-spliced counts across all replicates. Benjamini-Hochberg FDR correction was applied within each comparison. Constructs with |Δmis-spliced| ≥ 5 percentage points and FDR < 0.05 were called as differentially mis-spliced; constructs with fewer than 20 total deduplicated reads in either condition were excluded.

### 4.19. Nucleofection of differentially spliced constructs

Designed intron reporters that showed differential splicing by pooled NGS screening were subsequently tested individually in the corresponding cell lines. For each construct and cell line, 2 × 10^5^ cells were nucleofected with 300 ng of plasmid DNA using the Lonza 4D-Nucleofector system, SF Cell Line Solution, and a 16-well Nucleocuvette strip according to the manufacturer’s guidelines. Briefly, cells were collected, counted, and resuspended in SF nucleofection solution, mixed with plasmid DNA for an individual reporter construct, and transferred to a well of the 16-well Nucleocuvette strip. Cells were nucleofected using cell-line-specific 4D-Nucleofector programs: K562, FF-120; SH-SY5Y, CA-137; and HepG2, EH-100. Immediately after nucleofection, cells were transferred to complete growth medium and cultured for 48 hours prior to downstream analysis. All nucleofections were repeated in triplicate, with each replicate derived from distinct subcultures.

### 4.20. Fluorescence imaging of differentially spliced constructs

Individual intron reporter constructs that exhibited differential splicing in pooled NGS screening were evaluated by fluorescence microscopy 48 hours post-transfection. Cells were imaged 48 hours post-transfection on a Thermo Fisher Scientific EVOS M5000 inverted microscope using a 10× objective (AMEP4981, NA 0.3). Separate 16-bit TIFF images were captured in the GFP, TRANS, and TX Red channels for each construct in its corresponding target and non-target cell-line conditions. Exposure and gain settings were kept constant across matched samples within each experiment to allow qualitative comparison of reporter expression across cell lines.

Fluorescence images were processed in Python prior to single-cell quantification. To correct for uneven illumination, each fluorescence channel was flatfield-corrected by estimating a low-frequency illumination field using Gaussian smoothing (*σ* = 96 pixels), normalizing this field to its median value, clipping the resulting gain map to [0.5, 2.0], and dividing the raw image by the clipped gain map. Background fluorescence was estimated separately for each imaging channel and cell line as the median pixel intensity of non-cell regions in untransfected control images and subtracted prior to segmentation and fluorescence quantification.

Transfected cells were segmented using the GFP channel, which served as a transfection marker. Background-subtracted GFP images were smoothed with a Gaussian filter (*σ* = 1 pixel), and cell masks were generated by applying Otsu thresholding to background-subtracted non-zero pixels. Masks were refined by binary morphological closing and hole filling, and adjacent cells were separated by watershed segmentation on the Euclidean distance transform using a minimum peak distance of 8-12 pixels. Segmented objects were filtered to exclude debris, poorly segmented objects, and aggregates using an area range of 100-22,000 pixels^2^. For each remaining GFP-positive object, mean and median fluorescence intensities were extracted from both the GFP and TX Red channels. mScarlet/eGFP ratios were calculated per cell as

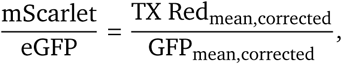

where TX Red_corrected_ corresponds to the background corrected TX red fluorescence across the segmented cell area and GFP_corrected_ corresponds to the background corrected GFP fluorescence across the segmented cell area. Per cell mScarlet/eGFP values were averaged across each cell to get overall mScarlet/eGFP ratios.

### 4.21. Nanopore sequencing of differentially spliced constructs

Following individual nucleofection of differentially spliced intron reporter constructs, cells were collected 48 hours post-nucleofection, and total RNA extraction, DNase treatment, reverse transcription, and targeted PCR amplification of the reporter region spanning the designed splice junction and construct-specific barcode were performed as described in **ProtoIntron RNA extraction, library generation, and NGS**. Purified PCR amplicons for each construct across their corresponding cell lines were normalized to 20 ng/*μ*L using a NanoDrop (Thermo Fisher) and sent to Quintara Biosciences for nanopore sequencing.

### 4.22. Computational design of *E. coli σ*70 ProtoPromoters

ProtoPromoter designs were produced through a multi-stage generation and optimization pipeline: autoregressive promoter generation with a genome language model, an initial energy-based rejection-sampling filter to enrich for promoter-like candidates, Metropolis-Hastings MCMC refinement against a composite energy function and an operator-site presence selection to enrich for designs with a repressor operator positioned to occlude the core promoter. During the first optimization stage, candidates were initially generated with the Evo 2 7B genome language model. Each sequence was produced by conditioning the model on a native RegulonDB *σ*70 promoter of length 80 nt (Salgado et al., 2023), supplied as a prompt, and extending autoregressively to a fixed length of 100 nt. Evo 2 sampling used temperature 0.9, top-*k* = 4 and top-*p* = 1.0. This procedure produced 804,500 candidate promoters.

Each candidate was then scored under a composite energy function. The composite energy was the weighted mean of three penalties. The first was a promoter-strength penalty from Promoter Calculator (LaFleur et al., 2022), evaluated as the plus-strand dGtotal. The second was a *σ*70 box-quality penalty from a RegulonDB-derived position-weight model of the −35 and −10 boxes, with consensus TTGACA and TATAAT and an optimal 17-bp spacer. The third was a transcription-factor-site penalty from a MEME/FIMO scan against 97 *E. coli* TF motifs, which penalized any cryptic regulatory site. The Promoter Calculator constraint was weighted 5× that of the *σ*70 box-quality and MEME/FIMO constraints. This step accepted 4,295 of the 804,500 generated candidates and rejected the remaining 800,205, enriching the pool for promoter-like sequences before optimization.

The surviving candidates were then further optimized against the same composite energy with a Metropolis-Hastings MCMC sampler and addition of an operator site presence filtering constraint. Briefly, each promoter was scanned for inverted-repeat operator candidates, defined as a pair of half-sites of ≥ 7 bp separated by a spacer of ≤ 1 bp with at most one mismatch per half-site, where mismatch was scored as the Hamming distance between half-site 1 and the reverse complement of half-site 2. A promoter was kept only if at least one such operator overlapped the −10 box, the −35 box, or the TSS by ≥ 3 bp such that it was positioned to sterically occlude RNA-polymerase recruitment or initiation. At each step, a proposal was drawn from a uniform single-nucleotide mutation generator, a position was chosen uniformly along the sequence and substituted by a base drawn uniformly over A, C, G, T, and the mutated sequence was rescored under all constraints. The proposal was accepted or rejected by the Metropolis criterion, comparing the proposal’s composite energy to the current sequence’s energy. Improvements were always accepted, and energy-increasing moves were accepted with probability *e*^(−Δ*E*/*T*)^. The chain was annealed with a geometric temperature schedule, where

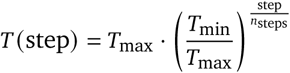

Following optimization, 93 final candidates for experimental validation were selected based on lowest energy and confirmed presence of an operator site occluding the −35 box, −10 box, or TSS.

### 4.23. Experimental validation of *σ*70 promoters

Sequences encoding generated *σ*70 promoter candidates, along with overhangs corresponding to an upstream multiple cloning site and downstream backbone context to facilitate integration into the reporter plasmid, were synthesized as eBlocks. To clone the synthesized promoters into the reporter plasmid, parent plasmid pAM002_BP_pZE21-_tetR_GFP_kanR was digested with KpnI-HF and SalI-HF (New England Biolabs) according to the manufacturer’s recommendations and subsequently gel purified to remove undigested product (Qiagen). Candidate promoter sequences were then inserted into the digested backbone using Gibson Assembly with NEBuilder HiFi DNA Assembly Master Mix (New England Biolabs) according to the manufacturer’s recommendations. Assembled products were plated overnight on LB agar plates containing 50 *μ*g/ml kanamycin. Individual colonies were then selected, sequence confirmed via nanopore sequencing (Quintara Biosciences), and miniprepped (Qiagen) for subsequent fluorescence characterization. Of the 93 ordered candidates, 86 were successfully synthesized and cloned.

50 ng of sequence-confirmed plasmid DNA was transformed into Stellar *E. coli* HST08 Competent Cells (Takara Bio). Transformants were grown overnight in LB containing 50 *μ*g/ml kanamycin. The following day, 4 *μ*l of overnight culture was transferred to 200 *μ*l of LB containing 50 *μ*g/ml kanamycin and grown for 8 hours with shaking. After 8 hours of growth, cultures were normalized to an optical density of 1.0. eGFP fluorescence was then measured using monochromator-based excitation at 485 nm with a 20 nm bandwidth and emission at 535 nm with a 20 nm bandwidth, with 9 measurements taken per well in a 3 × 3 circular pattern. All fluorescence measurements were assayed in triplicate using independent transformation replicates.

For experiments testing the effect of an additional operator site directly downstream of the TSS, each Proto-Promoter backbone was digested with KpnI-HF (New England Biolabs) according to the manufacturer’s recommendations and subsequently gel purified to remove undigested product (Qiagen). Oligos containing an extra operator site and appropriate backbone overhangs were ordered from IDT, before being annealed and inserted into the digested backbone using Gibson Assembly with NEBuilder HiFi DNA Assembly Master Mix (New England Biolabs) according to the manufacturer’s recommendations.

### 4.24. Computational design of ProtoRepressors

Sequence-specific ProtoRepressors were designed against the operator sites of sixteen ProtoPromoters (ProtoPromoter 6, 7, 8, 12, 33, 37, 40, 44, 49, 52, 56, 58, 62, 77, 79, and 85) through a three-stage generation-and-optimization pipeline: a sourcing-and-rejection-sampling stage (Stage 1) that enriched for scaffolds with predicted operator binding; a sequence-design stage (Stage 2) that imposed motif-contact and biophysical rejection sampling; and a Metropolis-Hastings MCMC refinement stage (Stage 3) that optimized the binding interface for predicted operator specificity.

In Stage 1, candidates were drawn from two complementary pools and filtered by rejection sampling for predicted operator binding. In the first pool, *de novo* candidate transcription factors were generated with the Evo 2 7B genome language model (evo2_7b). Each generation was prompted with the ProtoPromoter sequence followed by 290 nt of sequence immediately upstream of the coding sequence of a known *E. coli* helix-turn-helix (HTH) DNA-binding protein, and autoregressive nucleotide-level sampling then continued downstream into candidate coding regions at temperature 1.0 with top-*k* = 4, generating 2,500 nt per prompt for 12,355 sampled sequences in total. Open reading frames were called with Prodigal in metagenomic mode (`-meta`) and translated prior to filtering (Hyatt et al., 2010).

Translated candidates were scored with a weighted constraint objective combining a DNA-binding-domain term (weight 2.5) and a protein-quality term (weight 1.5). The DNA-binding-domain term was computed with PyHMMER hmmsearch (Finn et al., 2011) against Pfam-A profiles for HTH-, AraC-, Cro-, and LacI-like families at a sequence E-value threshold of 5 × 10^−3^. The protein-quality term favored candidates 80-200 amino acids in length with low complexity ≤ 0.3, repetitiveness ≤ 0.3, and diversity ≥ 0.3. Candidates with an energy score greater than 0.3, or without an HTH-domain hit passing the E-value threshold, were removed, leaving 434 sequences. In parallel, a curated natural HTH seed pool was assembled by querying InterPro for classical and winged HTH, MarR, IclR, Crp/CAP, Cro/cI, GntR, TetR, AsnC, LuxR, MerR, *λ*-cI, AraC, and LysR family proteins; after clustering at 70% sequence identity with MMseqs2 (Steinegger and Söding, 2017), 156 natural HTH sequences were retained.

The combined pool was screened against all sixteen operators in an all-against-all design. Each candidate protein-operator pair was modeled as a homodimer bound to the operator with 10 bp of flanking context on each side and co-folded with AF3 and Boltz-2 (Abramson et al., 2024; Passaro et al., 2025). Retention was determined by hard filters requiring Boltz-2 overall ipTM, protein-DNA ipTM, and protein-protein ipTM each ≥ 0.5 and protein pLDDT ≥ 70. Pairs passing these filters were ranked by a weighted structural-confidence composite combining protein-DNA ipTM (weighted 3× for each model), protein-protein ipTM (2×), overall ipTM (1×), and protein pLDDT (1×), used for downstream prioritization. This yielded 624 candidate ProtoRepressor-operator pairs.

Surviving pairs were positioned on their operators by template-guided superposition, providing the design-ready geometry used for sequence design. For each promoter, idealized B-form operator DNA containing the operator sequence plus 10 bp of ProtoPromoter flanking sequence on each side was generated with 3DNA/DSSR-X3DNA (Lu and Olson, 2008). Twenty natural HTH-DNA crystal structures (PDB 1QPI, 1R8D, 2KEI, 2OR1, 2VZ4, 2XRO, 2ZHG, 3BDN, 3ZQL, 4EGY, 4EGZ, 4L62, 4PXI, 4WLS, 5D8C, 5YEJ, 6JGW, 7TEA, 7TEC, and 8SVD) were used as docking templates. For each template, the template DNA was aligned to the synthetic operator by DNA backbone-atom superposition, and the candidate ProtoRepressor was placed into the major groove by C*α* superposition of its predicted HTH recognition helix onto the corresponding recognition helix in the template complex. This was repeated across all passing operator-template combinations to generate template-guided starting models.

In Stage 2, sequence-specific DNA-binding proteins were designed onto these starting models with a LigandMPNN (Dauparas et al., 2025) workflow adapted from Glasscock et al. (2025). Following that work, a prefilter calibration step was applied: a maximum-likelihood predictor over interface ddG and contact molecular surface (CMS) defined an inexpensive pass/fail filter (filter_eq/filter_cut) that triaged sequences before full Rosetta evaluation. LigandMPNN sequences were then generated with the operator DNA held as fixed context. Designs were accepted only if they satisfied a motif-contact requirement (≥ 1 protein-DNA base contact at a 4.0 Å heavy-atom cutoff) together with the calibrated Rosetta DNA-binding metrics (interface ddG, CMS, shape complementarity, buried-unsatisfied polar atoms, and base/bidentate hydrogen-bond terms, see **Table S2**) (Chaudhury et al., 2010; Glasscock et al., 2025).

Design proceeded in three increasingly deep production rounds: round 1 at one sequence per structure and decoding temperature 0.1 across all templates; round 2 at four sequences per structure and temperature 0.2 over the top ≤ 64 templates per promoter (ranked by the structural-confidence composite); and round 3 at fifty sequences per structure and temperature 0.1. If a round produced fewer than 20 passing designs for a promoter, it was repeated with the decoding temperature increased by 0.03.

The workflow was then extended with an AF3 co-folding and Rosetta-rescoring cascade. Each surviving design was co-folded as a protein-operator complex with AF3 and retained only if it passed complex-confidence filters: protein-DNA ipTM ≥ 0.6, overall ipTM ≥ 0.6, protein-protein ipTM ≥ 0.7, and protein pLDDT ≥ 70. Surviving models were rescored in PyRosetta, requiring CMS ≥ 225 Å^2^, interface_sc ≥ 0.60, packstat ≥ 0.55, base_score ≥ 10, bidentate_score ≥ 1.0, and buried-unsatisfied polar atoms ≤ 2.0. This design, co-fold, rescore, template-reselection loop was iterated for three cycles, each cycle using the strongest predicted binders, ranked by the same composite score, to seed the next; after the final cycle, up to the top 100 candidates per promoter advanced to Stage 3.

In Stage 3, candidate ProtoRepressors were optimized against a fixed operator using an interface-focused Metropolis-Hastings MCMC loop in which both AF3 complex confidence and operator specificity were active optimization signals. As before, the operator was held constant while LigandMPNN acted as the mutation-proposal generator. Refinement ran over a 500-step trajectory in a single interface-focused configuration: residues within 8 Å of the DNA were variable, while residues beyond 8 Å were fixed. At each step, ten sequences were generated per structure at decoding temperature 0.1, and the best-scoring (lowest-*S*) proposal was selected; it was accepted if it lowered the objective (Δ*S* < 0) and otherwise accepted with probability *e*^(−Δ*S*/*T*)^.

Each proposal was scored by a composite objective *S*, computed as a weighted sum of per-term scores that the framework scales to [0, 1] with 0 denoting the best value and 1 the worst (for example, a pLDDT of 100 maps to 0). *S* therefore behaves as a cost that the MCMC minimizes:

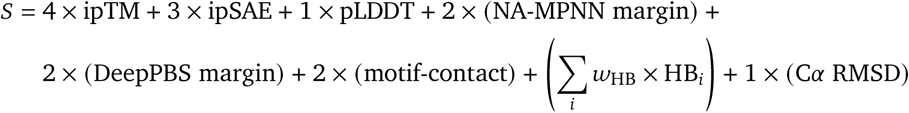

where every term is the framework-scaled [0, 1] score (0 is best), C*α* RMSD is measured against the template-guided starting model, HB_*i*_ are AF3 phosphate- and base-contact hydrogen-bond terms (weights 0.5 and 1.5), and all weights were fixed across promoters.

Off-target sets were fixed for each design before MCMC and comprised six operators: up to three same-length palindromic operators drawn from the ProtoPromoter panel, and three variants generated by random, non-palindrome-preserving substitution of the cognate operator at three positions (Hamming distance 3). AF3 contributed off-target specificity terms computed by evaluating each ProtoRepressor against the off-targets and taking the maximum off-target protein-DNA ipTM and ipSAE and the on-target-minus-off-target ipTM and ipSAE margin.

NA-MPNN specificity was evaluated with the dedicated specificity checkpoint `s_70114.pt` (Kubaney et al., 2025). Operators were scored as the sum of per-position log-probabilities under the predicted matrix, using a dyad-aware orientation-maximization procedure in which one half-site was scored against the forward motif and the other against the reverse complement, taking the maximum across orientations; half-sites were defined from the ProtoPromoter generation process. The NA-MPNN specificity margin was the cognate-operator score minus the highest-scoring off-target, with an analogous half-site margin also computed. DeepPBS specificity was computed as the difference between the highest scoring binding motif minus the desired binding motif from the input AF3 structure (Mitra et al., 2024). The two readouts were combined into a consensus operator-readout specificity score, *C*_consensus_ = *z*_NA-MPNN_ + *z*_DeepPBS_, with each *z*-score computed relative to the distribution of readouts across natural and scrambled HTH sequences; higher *C*_consensus_ values indicated stronger predicted discrimination of the cognate operator from off-targets.

Proposed designs were additionally selected to preserve direct motif engagement through a base-specific contact gate: beyond requiring a protein-DNA contact, designs were optimized for at least one unique contacting protein residue and at least one unique contacted DNA base position, defined at a 4.0 Å heavy-atom cutoff to DNA base atoms. The best-scoring candidates per promoter (up to 14) were selected for synthesis and experimental testing, with the number per promoter depending on how many reached satisfactory optimization and 2 promoters producing no passing candidates.

### 4.25. Cloning of ProtoRepressor candidates

Sequences encoding candidate repressors were synthesized as eBlocks with appropriate homology arms to facilitate Gibson assembly into a pBAD-derived expression backbone. To generate the repressor expression backbone, pAraCas9_+Sp2+Sp6+_I-SceI was PCR-amplified using AraC backbone extraction primers to remove the Cas9 coding sequence and guide RNA while preserving the pBAD/AraC regulatory context. The PCR product was treated with FastDigest DpnI according to the manufacturer’s recommendations (Thermo Scientific) to remove template plasmid before being gel purified to remove any additional residual template and nonspecific products (Qiagen).

Repressor sequences were inserted into the purified backbone using Gibson Assembly with NEBuilder HiFi DNA Assembly Master Mix (New England Biolabs) according to the manufacturer’s recommendations. Assembled products were plated overnight on LB agar plates containing 50 *μ*g/ml spectinomycin. Individual colonies were selected, sequence confirmed, and miniprepped according to manufacturer guidelines for subsequent characterization.

### 4.26. Experimental screening of ProtoRepressor candidates

Sequence-confirmed repressor plasmids and corresponding ProtoPromoters were transformed into Stellar *E. coli* HST08 Competent Cells (Takara Bio) using 50 ng of plasmid DNA each. Transformants were grown overnight in LB containing 50 *μ*g/ml kanamycin and spectinomycin. The following day, 4 *μ*l of overnight culture was transferred to 200 *μ*l of LB containing 50 *μ*g/ml kanamycin and spectinomycin with 2 mg/ml arabinose for ProtoRepressor induction and grown for 8 hours with shaking. After 8 hours of growth, 10 *μ*L of each culture was transferred to a 96-well round bottom plate, pelleted, and resuspended in PBS for subsequent analysis by flow cytometry.

Flow cytometry was performed with an Attune NxT flow cytometer equipped with an autosampler. Samples were acquired from 96-well plates using identical detector settings, with forward scatter, side scatter, and BL1 detector settings held fixed throughout acquisition.

To assess orthogonality of ProtoRepressor activity, the highest-performing repressor candidates from each of the four strongest ProtoPromoters were selected, co-transformed, and measured as described above, with each of the other tested ProtoPromoter reporter plasmids. All experiments were conducted in at least triplicate, with each replicate corresponding to an independent transformation.

### 4.27. Analysis of repression using flow cytometry

Raw FCS files were processed using a multi-step gating pipeline. First, events with non-finite or non-positive FSC-A or SSC-A values were excluded. To remove instrument start-up and end-of-acquisition artifacts, the first 250 and last 100 temporally-ordered valid events per well were discarded. The remaining cells were subjected to a Gaussian mixture model (GMM) gate on the FSC-A vs. SSC-A plane: a single-component GMM was fit to log-scaled FSC-A and SSC-A values using Cytoflow’s GaussianMixtureOp (`num_components=1`, `sigma=2`), and events falling outside the 2*σ* boundary of the fitted Gaussian were excluded (Teague, 2022). Doublets were then removed by filtering on the FSC-A/FSC-H ratio, with events deviating more than 2 × 1.4826 × MAD from the per-well median ratio excluded. GFP fluorescence (BL1-A channel) was quantified as the median of all cells passing both gates.

Fold repression for each replicate pair was computed as the ratio of the median eGFP signal in uninduced wells to that in induced wells. Per-candidate geometric mean fold repression was calculated across replicate pairs as the exponent of the mean of the natural-log-transformed fold values. Significance was assessed using one-sided one-sample t-tests on the log_2_ fold repression values against null hypotheses of fold 1×, 1.25×, 1.5×, and in the case of the additional operator experiments, induced tetR and cI. Multiple testing correction was applied across all candidate designs using the Benjamini-Hochberg procedure.

### 4.28. Human multimer diversification with Proto

Diversification programs for human protein complexes were generated automatically by an AI coding agent (initial development with Claude Code with Opus 4.5/4.6 and refinement via Codex CLI with GPT-5.4/-5.5) from a curated set of major human pathways and complexes. The agent performed the initial curation with manual steering to ground the curation by leveraging existing databases, namely the Protein Data Bank (PDB) and UniProt.

For each complex, the agent emitted one Proto program and one configuration file, yielding 93 generated programs that together span 797 unique HUGO Gene Nomenclature Committee (HGNC) gene symbols and 249 complex definitions (150 hetero-complexes, 50 monomers, 27 homo-families, 15 hetero-families, and 7 homomers), of which 76 carried at least one representative PDB structure (**Data S3**). Each program encoded the diversification of all unique subunits of the complex.

Wild-type protein sequences were obtained by mapping each HGNC symbol to its UniProt accession and retrieving the corresponding amino-acid sequence, with each gene instantiated as a single protein segment within a single-segment construct. Complex stoichiometry was resolved through a combination of manual curation and agent-automated literature search and structure database cross references, where stoichiometry was specified explicitly for 43 complex entries and inferred for 206. The expanded stoichiometry was used for downstream AF3 complex scoring.

Within each program, every protein subunit was assigned an ESM3 generator (`esm3_sm_open_v1` checkpoint) with a sampling temperature of 0.3 and a masking strategy that proposed substitutions at 25% of sequence positions (Hayes et al., 2025). All subunits of a complex were optimized jointly by a single Metropolis-Hastings MCMC optimizer with one retained result and one proposal per step, under simulated annealing from *T*_max_ = 1 × 10^−1^ to *T*_min_ = 1 × 10^−2^, for a total of n_genes × n_steps_per_generator steps (with n_steps_per_generator = 3 by default).

Each diversified subunit was scored against five shared constraints. Two were ESMFold confidence terms (pLDDT and pTM). A TM-score similarity constraint compared the diversified subunit to its native ESMFold structure, applied as a threshold (retaining designs of native-like TM-score ≥ 0.7) only when the native model reached a target pLDDT of at least 0.6. An RMSD similarity constraint likewise compared the design to the native ESM-Fold structure, transformed to take values between 0 and 1 with a sigmoid inflection at 5.0 Å and slope 3.0 and the same minimum native pLDDT of 0.6. Finally, a base protein-quality constraint disabled length, diversity, and balanced-amino-acid checks but enforced low-complexity and repetitiveness limits.

After diversification, each complex was scored with AF3 with a local ColabFold MSA search, expanding subunits by stoichiometry and reporting pLDDT, pTM, ipTM, and PAE (the latter rescaled by 31.75 Å to provide a value between 0 and 1). For complexes with a representative PDB structure, the experimental coordinates were retrieved and the final AF3 prediction was compared against them by TM-score similarity (pLDDT threshold 50) and by all-atom aligned RMSD similarity (transformed to be within 0 and 1 with a sigmoid inflection of 5.0 Å and a slope of 3.0). For each complex, the program was run five times, each run being one MCMC trajectory that retained a single set of subunit sequences, with the highest-confidence set selected by AF3 pLDDT.

### 4.29. Human signaling pathway diversification with Proto

The *β*2-adrenergic signaling pathway program was generated from the same agent template as the human complexes above and then extended to encode additional ligands, static sequences, and conformational-ensemble constraints (initial development with Claude Code with Opus 4.5/4.6 and refinement via Codex CLI with GPT-5.4/-5.5). The pathway comprised eight human protein genes: ADRB2 (*β*2-adrenergic receptor), GNAS (G*α*s), GNB1 (G*β*), GNG2 (G*γ*), ADCY9 (adenylyl cyclase), PRKACA (PKA catalytic subunit), PRKAR1A (PKA regulatory RI*α* subunit), and CREB1. These were augmented components that form important interactions relevant to signal transduction: epinephrine ligand (binds to *β*2-adrenergic receptor), an ATP ligand (binds to adenylyl cyclase), two CREBBP KIX coactivator domains, and a CREB transcription-factor DNA motif together with its reverse complement. These were assembled into six scored complexes: the ligand-bound receptor monomer (ADRB2 + epinephrine; reference PDB 4LDO), the receptor-Gs complex (ADRB2, GNAS, GNB1, GNG2; reference PDB 3SN6), the G*α*-adenylyl cyclase complex (GNAS, ADCY9; reference PDB 6R3Q), the ligand-bound adenylyl cyclase monomer (ADCY9 + ATP; reference PDB 4USW), the PKA RI*α* holoenzyme (PRKACA ×2, PRKAR1A ×2; reference PDB 6NO7), and the CREB dimer (CREB1 ×2, CREBBP KIX ×2, and the forward and reverse-complement CREB DNA motifs; reference PDBs 1DH3, 1KDX). Subunits present in multiple copies were represented as repeated segments driven by a single tied generator.

The program was executed in three stages. In the first stage, the CREB-responsive DNA element was designed before any protein diversification. A 512-bp variable DNA segment was placed between fixed left and right genomic flanks to form a full-length Borzoi input of 524,288 bp (with each flank spanning 261,888 bp) and generated with an Evo 2 generator (checkpoint evo2_7b; top-*k* = 4, top-*p* = 1.0, temperature = 0.5) under rejection sampling over 300 proposals, prompted from a CREB design sequence (Brixi et al., 2026; Linder et al., 2025). The DNA objective maximized Borzoi-predicted activity on the human CREB1 ChIP-seq tracks in HepG2 (activity threshold 200.0). The central 50 bp of the resulting 512-bp design were then added to the CREB complex in both forward and reverse-complement orientations.

In the second stage, the eight pathway proteins were diversified from their wild-type sequences. Each gene was assigned a single ESM3 generator (Hayes et al., 2025) (`esm3_sm_open_v1`, temperature 0.3) tied across all stoichiometric copies of that subunit, with a masking strategy proposing substitutions at 25% of positions. The eight protein constructs were optimized jointly by a Metropolis-Hastings MCMC optimizer with one retained result and one proposal per step under simulated annealing from *T*_max_ = 1 × 10^−1^ to *T*_min_ = 1 × 10^−2^ for 40 steps. Each protein was scored on its primary copy by ESMFold pLDDT, ESMFold pTM, and an overall protein-quality constraint (threshold 0.15; low-complexity fraction ≤ 0.12, repetitiveness ≤ 0.08, minimum repeat length 2), for 24 protein-level constraints in total.

In the third stage, the diversified sequences were rescored using a rejection-sampling optimizer drawing from the existing results. Conformational dynamics were evaluated with four BioEmu (Lewis et al., 2025) structure-ensemble RMSD constraints: two probing the G*α*s (GNAS) GDP- versus GTP-bound states against the inactive (PDB 6AU6, chain A, residues 85-394) and nucleotide-exchange (3SN6, chain A, residues 85-394) references at 3,000 samples each, and two probing PKA regulatory-subunit (PRKAR1A) flexibility against homodimer (PDB 1RL3, chain A, residues 119-379) and tetramer (PDB 2QCS, chain B, residues 119-379) references at 1,000 samples each.

Each BioEmu ensemble constraint reported the minimum RMSD to each of the reference PDBs across the entire ensemble, where the RMSD was transformed to be between 0 and 1 with a sigmoid inflection at 3.0 Å with slope 3.0. BioEmu was sampled with default parameters and a batch size of 100. As a control, we also obtained BioEmu ensemble predictions of the native sequence or of synthetic sequences in which we used ProteinMPNN to sample a single backbone conditioned sequence for each of the reference PDBs (2 references for G*α*s and 2 references for the PKA regulatory-subunit) with temperature 0.1, followed by the same BioEmu ensemble analysis on the resulting sequence.

Whole-complex geometry was re-scored with AF3 using local ColabFold MSA search, evaluating pLDDT, pTM, ipTM, and PAE for each of the six complexes (24 complex-level constraints) (Abramson et al., 2024). The full pathway program therefore comprised nine generators (one Evo 2, eight ESM3) and 53 constraints across the three stages. Additional downstream metrics computed on the designed sequences and their AF3-predicted structures included the all-atom aligned RMSD values to the experimental structures (using pymol’s cealign algorithm), pocket-aligned RMSD of ligands (computed on the ligand atoms and using pymol’s cealign algorithm on the residues within 10 Å of the ligand in both the predicted and experimental structures), and the multimeric TM-score defined with respect to the experimental structures (using `USalign <designed_pdb> <experimental_pdb> -mm 1 -ter 1`) (Schrödinger, LLC, 2015; Zhang et al., 2022).

### 4.30. NSCLC circuit design with Proto

To generate an NSCLC-specific gating system for HSV-TK delivery, we constructed a program consisting of five optimization stages and five designed sequences: an EGFR binder, an enhancer, a promoter, an HSV-TK split intron, and a 3^′^ UTR off-switch. Each component was optimized separately, with regulatory-cassette components scored in the context of the previous design step and hypothetical integration site. The final cassette components were defined as a variable 5^′^ genomic locus mimicking potential lentivirus integration sites, a 500 bp enhancer, a 100 bp promoter, HSV-TK exon 1 (first 566 bp of HSV-TK), a synthetic intron of 301 bp, HSV-TK exon 2 (final 569 bp of HSV-TK), a 400 bp 3^′^UTR, and a variable 3^′^ genomic locus mimicking natural integration context. Cell-type selectivity was defined throughout as A549 lung adenocarcinoma (AlphaGenome ontology EFO:0001086) versus healthy lung (AlphaGenome UBERON:0002048). Regulatory optimization steps were scored inside the gene bodies of broadly/highly-expressed integration loci: GAPDH, ACTB, EEF1A1, and FTL, plus a known lentiviral integration spot HMGA2. Flanking context was obtained as the gene-body midpoint ± 8,192 bp from NCBI (GRCh38).

The first optimization stage designed a de-novo miniprotein binder against the EGFR ectodomain (UniProt P00533, AlphaFold model residues 25-645, chain A) to bias lentiviral entry toward EGFR-high tumor cells, targeting the domain-III epitope (hotspot residues 384, 408, 409, 443, 465, 467, 468 in native EGFR numbering). Backbones were generated by epitope-centered RFdiffusion3 and sequences by ProteinMPNN (inverse folding) (Butcher et al., 2025; Dauparas et al., 2022), under a rejection-sampling optimizer that drew 48 candidates per run and retained the top 8. Each candidate was scored in-loop on a consensus of two independent structure-prediction oracles, Boltz-2 and AlphaFold 2 (Passaro et al., 2025; Jumper et al., 2021). Interface ipTM and pLDDT were evaluated on both oracles—each entered as a separate maximization term (interface ipTM weight 1.0 per oracle; pLDDT weight 0.1 per oracle), summed in the energy objective rather than combined as their minimum.

Interface ipAE (weight 1.0) and an interface-contact term (≥ 2 intra-chain contacts at 14 and ≥ 2 inter-chain contacts at 20 Å; weight 0.25) were evaluated on AlphaFold 2. AlphaFold 2 predictions used a single recycle, while Boltz-2 used its default recycling. As a final filtering constraint, surviving candidates were re-scored with AlphaFold 3 and any with interface ipTM < 0.7 were rejected.

The second optimization stage designed a 500-bp enhancer with an Evo 2 generator (checkpoint evo2_7b; 2,048-bp prompt derived from upstream context of natural enhancer sequences) under rejection sampling with 500 samples per run and 10 retained. Designs were scored on cell-type-specific epigenomic tracks with Al-phaGenome, contrasting A549 against lung across the integration loci: the objective maximized the active-enhancer marks H3K4me1 (weight 4.0) and H3K27ac (weight 4.0) and chromatin accessibility (DNase/ATAC; weight 1.0 and 3.0 respectively) while minimizing the promoter-associated marks CAGE (weight 2.0) and H3K4me3 (weight 1.0) to push the design away from a promoter-like signature. A total of 7500 candidates were sampled.

The third optimization step designed a 100-bp promoter placed between the upstream enhancer (top 5 seeds from stage 2) and the downstream HSV-TK coding sequence to drive NSCLC-selective transcription. Candidate promoters were optimized and assessed in the aforementioned genomic context embedded immediately 3^′^ of the upstream enhancer such that each design was scored in the presence of its upstream regulatory element and host-locus chromatin. Promoters were initialized from natural-promoter sequence templates. Candidate updates were proposed by a uniform-mutation generator introducing three substitutions per proposal, and each trajectory was optimized by Metropolis-Hastings MCMC for 400 steps, with exponential temperature annealing from *T*_max_ = 1.0 to *T*_min_ = 1×10^−3^, retaining ten designs. Each proposal was scored by a weighted sum of an AlphaGenome promoter-mark term and a Puffin promoter-initiation term (Dudnyk et al., 2024; Avsec et al., 2026). The AlphaGenome term (weight 6.0) evaluated A549-versus-lung signal over the promoter interval, maximizing the A549-versus-lung margin for the initiation/active marks CAGE, H3K4me3, H3K27ac, and DNase under a per-mark weighting that prioritizes the initiation marks CAGE and H3K4me3 (weights CAGE 2.0, H3K4me3 2.0, H3K27ac 1.0, DNase 0.5), while minimizing the enhancer-associated mark H3K4me1 (weight 0.5) to keep the design promoter-like. In addition to AlphaGenome, a Puffin promoter-activity term (weight 6.0) was included as a weighted optimization term that scores designs toward a minimum activity threshold of 0.08 and sharpness threshold of 0.15.

The fourth optimization step designed a 301-bp synthetic intron embedded between two HSV-TK coding exons, such that selective intron excision in NSCLC cells would reconstitute a functional HSV-TK open reading frame. Candidate introns were optimized in the context of the full therapeutic cassette, including HSV-TK exon sequence flanking the intron, the top five promoter-enhancer combinations selected in the preceding step, and the specified genomic safe-harbor contexts. For each design, a 1-kb splice-evaluation window centered on the intron was embedded within a 16-kb AlphaGenome input containing the surrounding cassette and genomic context. Introns were initialized as random 301-bp sequences with fixed canonical GT donor and AG acceptor dinucleotides. Candidate updates were proposed by a uniform-mutation generator introducing three substitutions per proposal within the internal intron sequence, and each trajectory was optimized by Metropolis-Hastings MCMC for 150 steps with exponential temperature annealing from *T*_max_ = 1 × 10^−2^ to *T*_min_ = 1 × 10^−3^. Each proposal was scored by a weighted sum of AlphaGenome and SpliceTransformer constraints (Avsec et al., 2026; You et al., 2024). Al-phaGenome splice-site-usage predictions were evaluated on positive-strand tracks within a 3-nt peak-search radius at the fixed donor and acceptor sites, maximizing SSU in A549 and minimizing SSU in healthy lung with matched weights of 8.0. Scores were summed with equal weight across the top five promoter-enhancer cassette contexts and genomic integration contexts. A low-weight SpliceTransformer boundary term evaluated the concatenated left-flank, intron, and right-flank sequence with 4-kb left and right contexts to preserve correctly placed splice-site geometry, using the score 1 − 0.5 × ( *p*_donor_ + *p*_acceptor_), where *p*_donor_ and *p*_acceptor_ are the predicted donor and acceptor probabilities at the fixed splice boundaries.

The fifth subprogram designed a 400-bp 3^′^ UTR off-switch with a uniform-mutation generator (3 substitutions per proposal, seeded from a natural-3^′^UTR template) optimized by Metropolis-Hastings MCMC over 8000 steps with exponential temperature annealing from *T*_max_ = 1 × 10^−2^ to *T*_min_ = 1 × 10^−3^. The objective installed microRNA response elements for drivers abundant in healthy lung and depleted in NSCLC (miR-486-5p, miR-144-3p, miR-197-3p, miR-451a, let-7d-5p) under a consensus driver term of total weight 3.0, distributed across drivers in proportion to their TCGA-LUAD lung-versus-A549 abundance (log2 RPM ratio). Each driver was required to be called concordantly by both miRanda and TargetScan (John et al., 2004; McGeary et al., 2019) and contributed two equally-weighted sub-terms (half the per-driver weight each): miRanda sites used an alignment-score threshold of 140 and a duplex-energy threshold of −20 kcal/mol with a saturating repression threshold of 0.8, while TargetScan sites required a canonical ≥ 7-mer seed match (6-mer sites excluded) with a saturating threshold of 1.0. Tumor-high oncomiR sites (miR-21-5p, miR-210-3p, miR-31-5p) were escaped under both callers, each as a separate minimization term of weight 1.0. Sequence realism was enforced against measured natural 3^′^ UTR statistics: dinucleotide-composition match (weight 0.6), GC content 35-60% (weight 0.4), and maximum homopolymer length ≤ 12 (weight 0.4). A low-weight AlphaGenome RNA-seq contrastive prior, maximizing the A549-versus-lung expression margin, was added as one term of weight 0.15 per genomic context and summed across three integration loci (HMGA2, GAPDH, EEF1A1).

Much of the program’s final composition was determined not *a priori* but through iterative, agent-driven exploration: Claude Code and Codex CLI were used to author, execute, score, and revise candidate Proto optimization stages, comparing alternative generators, optimizers, scoring models, and biological targets for each design objective and retaining only the choices that empirically performed best. For the EGFR binder, the agents evaluated several published binder-design pipelines: BindCraft, Germinal, and RFdiffusion3 + ProteinMPNN, and selected RFdiffusion3 + ProteinMPNN under Boltz-2/AlphaFold-2 consensus selection with AF3 as a final filtering oracle, after BindCraft and Germinal both produced zero accepted designs out of 40 in a 12-hour run on this epitope and single-oracle optimization repeatedly yielded binders that one structure predictor scored highly but the others rejected. For the 3^′^UTR off-switch, the agents derived a data-grounded NSCLC versus healthy miRNA target set from TCGA-LUAD tumor-versus-normal miRNA abundance, then narrowed it to the sites called concordantly by both miRanda and TargetScan while screening out tumor-high oncomiRs. Comparable exploration comparing distributions of natural enhancers and promoters versus scrambled sequences tuned the per-mark and per-context objective weights used in these optimization stages.

### 4.31. Statement on AI use

We used AI coding agents (Claude Code and Codex CLI) to assist in developing the software infrastructure, user interfaces, and in conducting some of the Proto program designs; the role of agentic workflows in design specification is itself described as part of the methodology of this study. We used AI-based web tools (ChatGPT, Claude.ai, and Gemini) to assist with manuscript writing, including drafting and editing of prose. We did not use AI tools to generate any figures. All authors take full responsibility for the entire content of the manuscript, including any portions developed with the assistance of AI tools, and have reviewed and verified all text, code, designs, and results.

## 5. Data, code, and materials availability

We make the following open-source repositories available:

- Implementation and Python API for the Proto language: https://github.com/evo-design/proto-language
- Implementation and Python API for the Proto tools layer: https://github.com/evo-design/proto-tools

A web interface for Proto is available at https://proto.evodesign.org/. Sequencing data will be made available in public repositories upon publication of the manuscript. Experimental materials will be made available upon reasonable request to the corresponding author.

We also make the following supplementary data files available:

- **Data S1:** A ZIP file containing computational scores for intron designs, AlphaGenome predictions visualized as genome tracks, and plasmid and primer sequences used for experimental screening of designed introns.
- **Data S2:** An Excel spreadsheet containing sequences and scores for all tested promoter and repressor designs and plasmid and primer sequences related to repressor and promoter design experiments.
- **Data S3:** An Excel spreadsheet containing the human proteins and protein complexes subjected to structure-preserving diversification.

## 6. Acknowledgments

We acknowledge Garyk Brixi, Joseph Caputo, Drew Endy, Xiaojing Gao, Yukun Alex Hao, Brian Kang, David Li, Niko McCarty, Luis Santiago Mille-Fragoso, Jean-Sebastien Paul, April Pawluk, Chiara Ricci-Tam, Ivan Specht, Talal Widatalla, Xiaowei Zhang, and members of the Laboratory of Evolutionary Design for helpful discussions and assistance with manuscript preparation. We acknowledge Richard Bao, Nikash Chhadia, Nahum Maru, and Rachel Park for feedback on engineering tasks.

A.T.M. acknowledges funding support from the Knight-Hennessy Graduate Scholarship Fund and National Science Foundation Graduate Research Fellowship. B.V. acknowledges funding support from the National Library of Medicine (NLM) and the MAC3 Data Science and Precision Medicine Fellowship. S.H.K. acknowledges funding support from the National Science Foundation Graduate Research Fellowship. B.L.H. acknowledges funding support from Schmidt Sciences AI2050, the Gates Foundation, the Chan Zuckerberg Initiative, Arc Institute, Stanford Center for Digital Health, and Stanford Human-Centered Artificial Intelligence (HAI) Hoffman-Yee Research Grants.

## 7. Author contributions

A.T.M., D.G., and B.L.H. conceived the study. B.L.H. supervised the study. B.L.H. developed the initial specification of the Proto language. D.G. and B.V. developed the initial version of the software infrastructure. A.T.M., D.G., B.V., L.B.-A., P.Y., S.H.K., and B.L.H. contributed to the software infrastructure. A.T.M., D.G., and B.V. developed the initial version of the tools infrastructure and documentation. A.T.M., D.G., B.V., L.B.-A., P.Y., S.H.K., and B.L.H. contributed to the tools infrastructure and documentation. T.M. and E.H. developed the initial version of the user interface. A.T.M., D.G., B.V., L.B.-A., E.H., S.H.K., and B.L.H. contributed to the user interface. D.G. and B.L.H. conducted the reimplementation of existing design campaigns in Proto. A.T.M. and B.L.H. conducted ProtoIntron design and sampling. A.T.M. conducted the experimental testing of the ProtoIntrons. A.T.M. conducted ProtoPromoter and ProtoRepressor design and sampling. A.T.M. conducted the experimental testing of the ProtoPromoters and ProtoRepressors. B.L.H. conducted the proteome-scale diversification and signaling pathway diversification design tasks. A.T.M. and B.V. conducted the NSCLC circuit design task. E.A.A. and B.L.H. acquired funding for the project. A.T.M., D.G., B.V., S.H.K., and B.L.H. wrote the initial version of the manuscript. All authors wrote the final version of the manuscript.

## Supplementary Material

### A. Supplementary figures and tables

**Figure S1.**
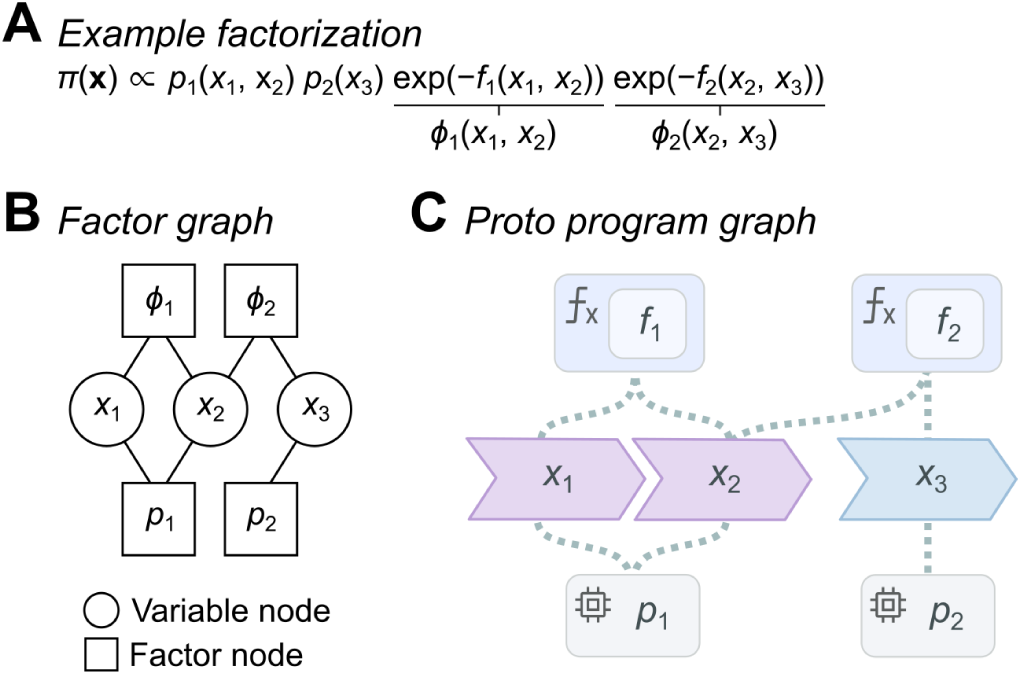
| Example factorization and graphical representation. Proto optimizes an energy-based model with a design target distribution *π*(*x*). (**A**) An example factorized form of *π*(*x*) containing three sequences *x*, two generative priors *p*, and two constraint functions *f* that each define a Boltzmann factor *φ*. (**B**) A factor graph representation of the factorized distribution in (**A**) in which circular nodes indicate the random variables (sequences) and the square nodes indicate the factors (generators and constraints). (**C**) This factor graph representation forms the basis of the stylized Proto program graph; compare to Figure 1E.

**Figure S2.**
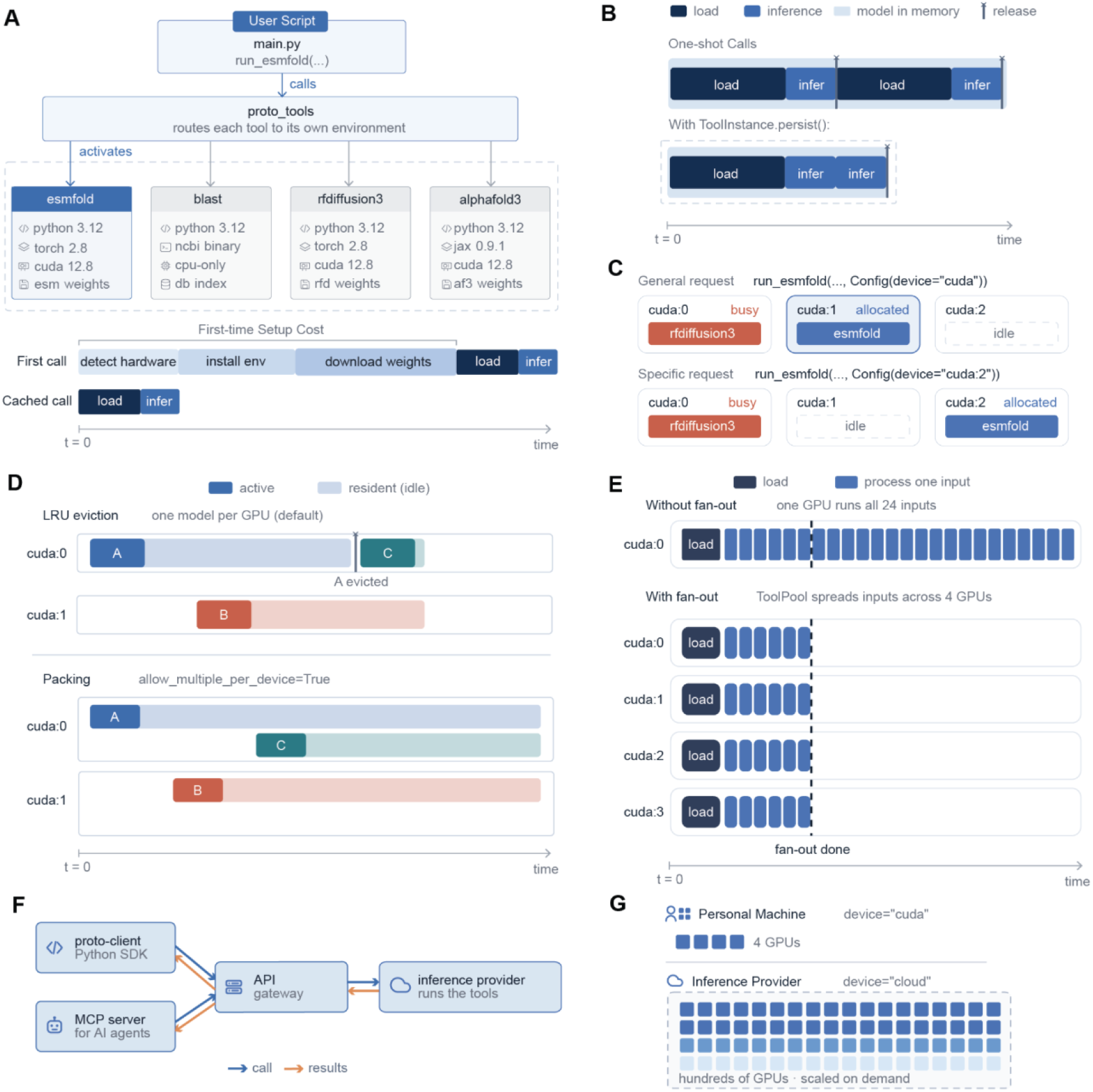
| Illustration of the architecture for Proto tools. (**A**) Environment isolation system. The primary process from the user’s main script dispatches a subprocess inside individual virtual environments for each tool. The first time the tool is called, the virtual environment is installed, and weights are downloaded. All subsequent calls on a system avoid this set-up cost. (**B**) Tool subprocesses can operate in one-shot or persistent execution modes. In one-shot mode, the model is loaded in, performs inference, and then immediately exits. If subsequent calls to a tool are being made, persistence allows the user to control how long the process lives to avoid the time costs of reloading the model. (**C**) Automatic device allocation. When a user calls a model, the infrastructure layer automatically routes the model to an available device. We also enable full placement customization; the user can specify a specific device they would like the model to be loaded onto. (**D**) Device allocation behaviors are fully customizable. If all devices are busy, the least recently used model will be offloaded to make room for the next model call. Users can also pack multiple models on the same device if space allows. (**E**) Proto tools enables GPU fan out, which spreads jobs out across all available local GPUs to maximize the utilization of available local hardware. (**F**) Proto also supports an MCP for AI agents and API gateway for cloud-based inference. (**G**) The API gateway will allow users to scale up their compute utilization to hundreds of GPUs.

**Figure S3.**
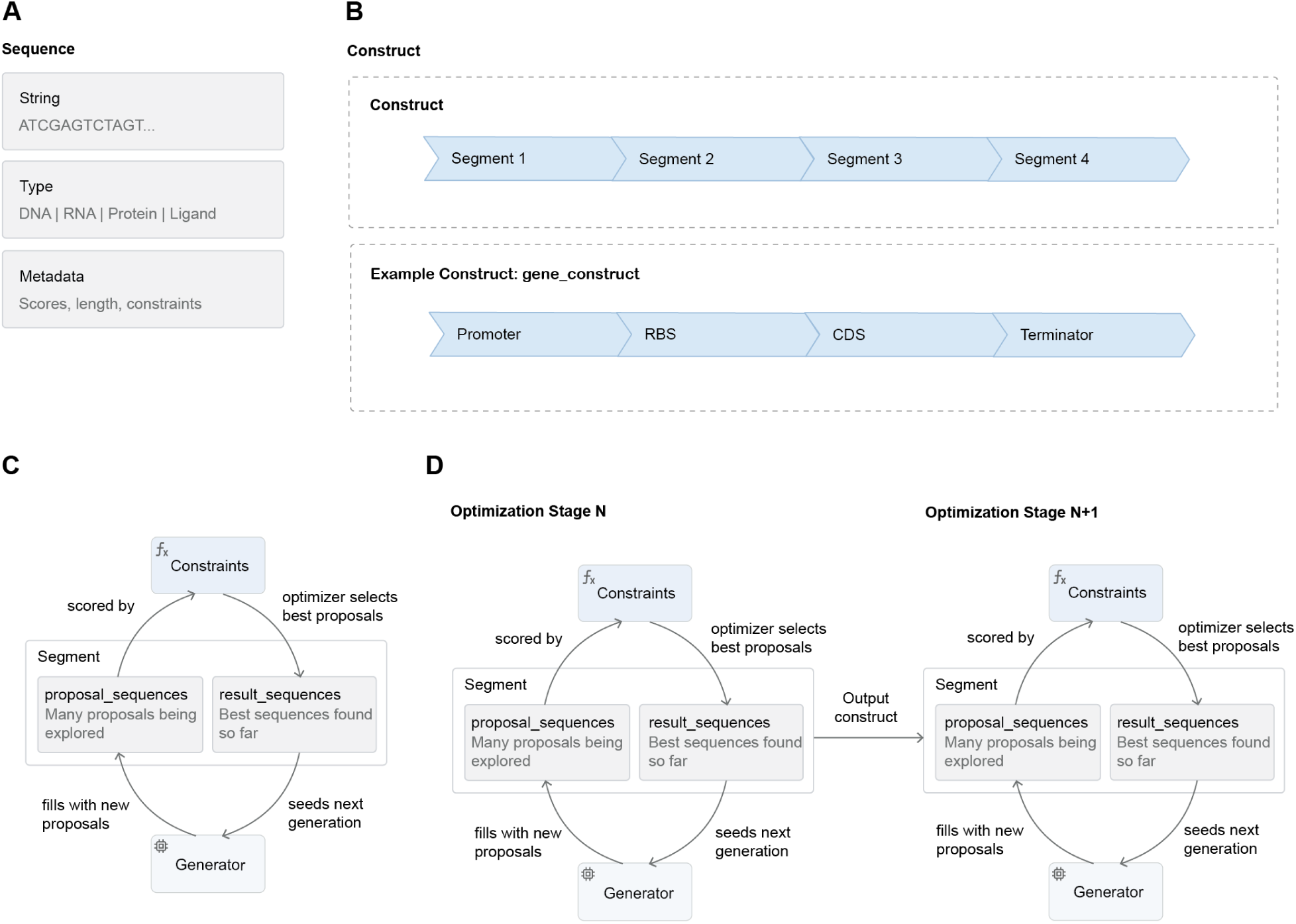
| Components and overall workflow of the Proto language software implementation. (**A**) Proto implements a Sequence object with the actual sequence value, the sequence type, and associated metadata (computed by constraints or the generator). (**B**) In addition to the Sequence class, we define two additional classes: Segments and Constructs. A Segment wraps a Sequence, but also keeps track of a batch dimension (where multiple sequence variations could correspond to the same Sequence variable). A Construct is defined on a list of Segments and is meant to indicate that Segments are part of the same contiguous sequence, and will therefore concatenate multiple Segments in a user-defined order as part of logging and visualization. However, the implementation keeps Segments logically separated even in a Construct. (**C**) Segments contain two internal variables: `proposal_sequences` and `result_sequences`, where the first contains generated sequences prior to any scoring by the program’s constraints and the second contains post-scored sequences. Optimizers manage how these variables are populated and which sequences from `proposal_sequences` get preserved in `result_sequences`. (**D**) For multistage programs containing multiple Optimizers, the sequence handoff between these stages involves the final sequences produced by an optimizer at stage *N* seeding the `proposal_sequences` variable in stage *N* + 1.

**Figure S4.**
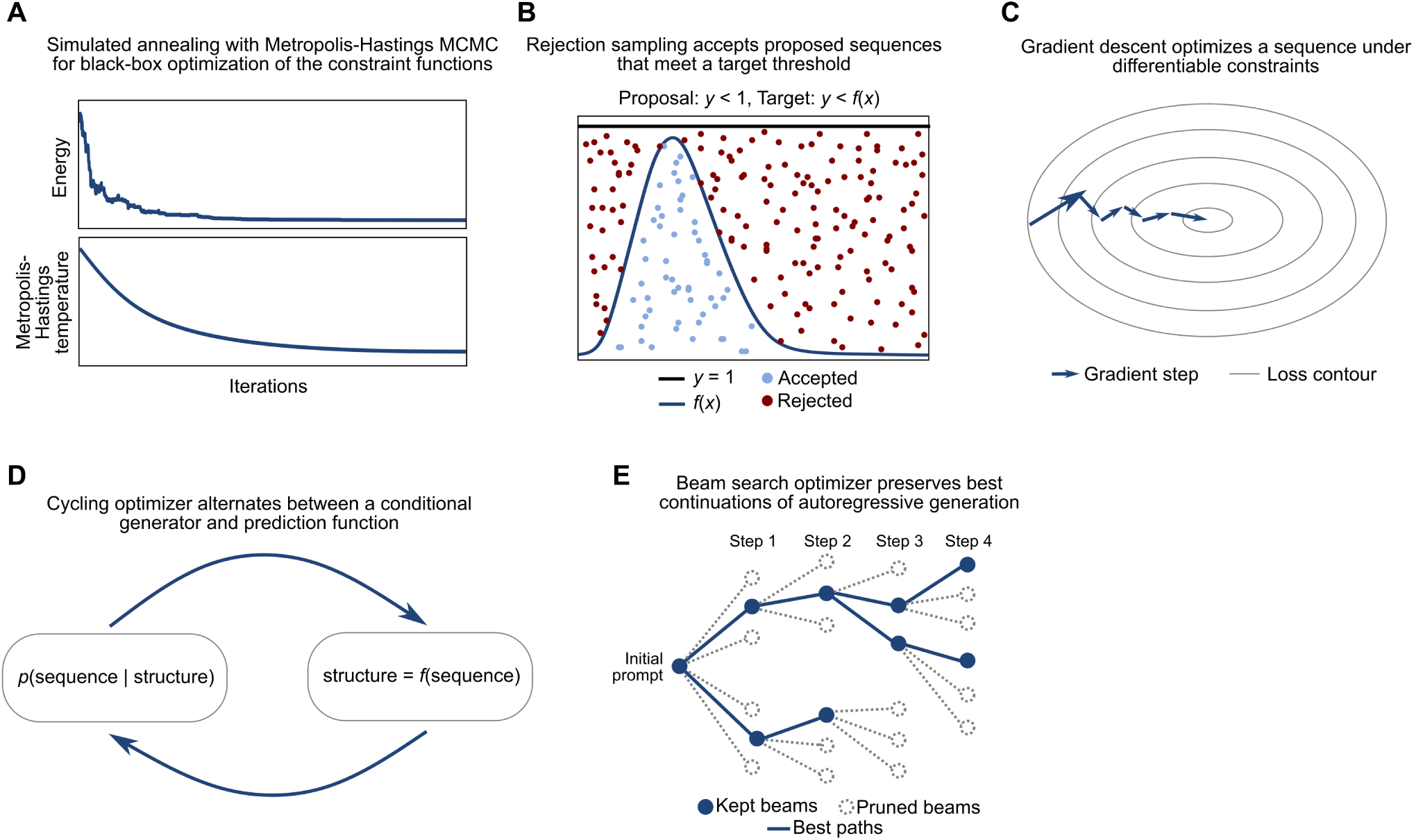
| Graphical visualizations of different Proto optimizers. (**A**) Simulated annealing is implemented with a Metropolis-Hastings MCMC algorithm with a temperature decay that focuses designs on local minima under a user-provided energy function. (**B**) A rejection sampling optimizer pairs a generative model and a set of scoring functions with user-defined thresholds, accepting sequences that pass all filters and rejecting those that fail at least one filter. (**C**) The gradient-based optimizer minimizes a differentiable loss function (in this case, the combined constraint function) by modifying the sequence on each iteration of gradient descent. (**D**) A cycling optimizer requires a special generator in the form of *p*(*x* | *y*) and is defined according to a function *f* such that *y* = *f* (*x*), where *x* is the sequence. Sequences are generated by alternating between samples from *p* and computation of *f*, where *y* could be a complex structured variable. The main instantiation of this in Proto is the alternation between a structure prediction model (Boltz-2) and a structure-conditioned sequence design model (ProteinMPNN) as in the Protein Hunter method. (**E**) A beam search optimizer is specific to autoregressive sequence generation with constraints that admit partial-sequence scoring. Multiple autoregressive continuations of a prompt are sampled, scored, and a subset are preserved for future continuations.

**Figure S5.**
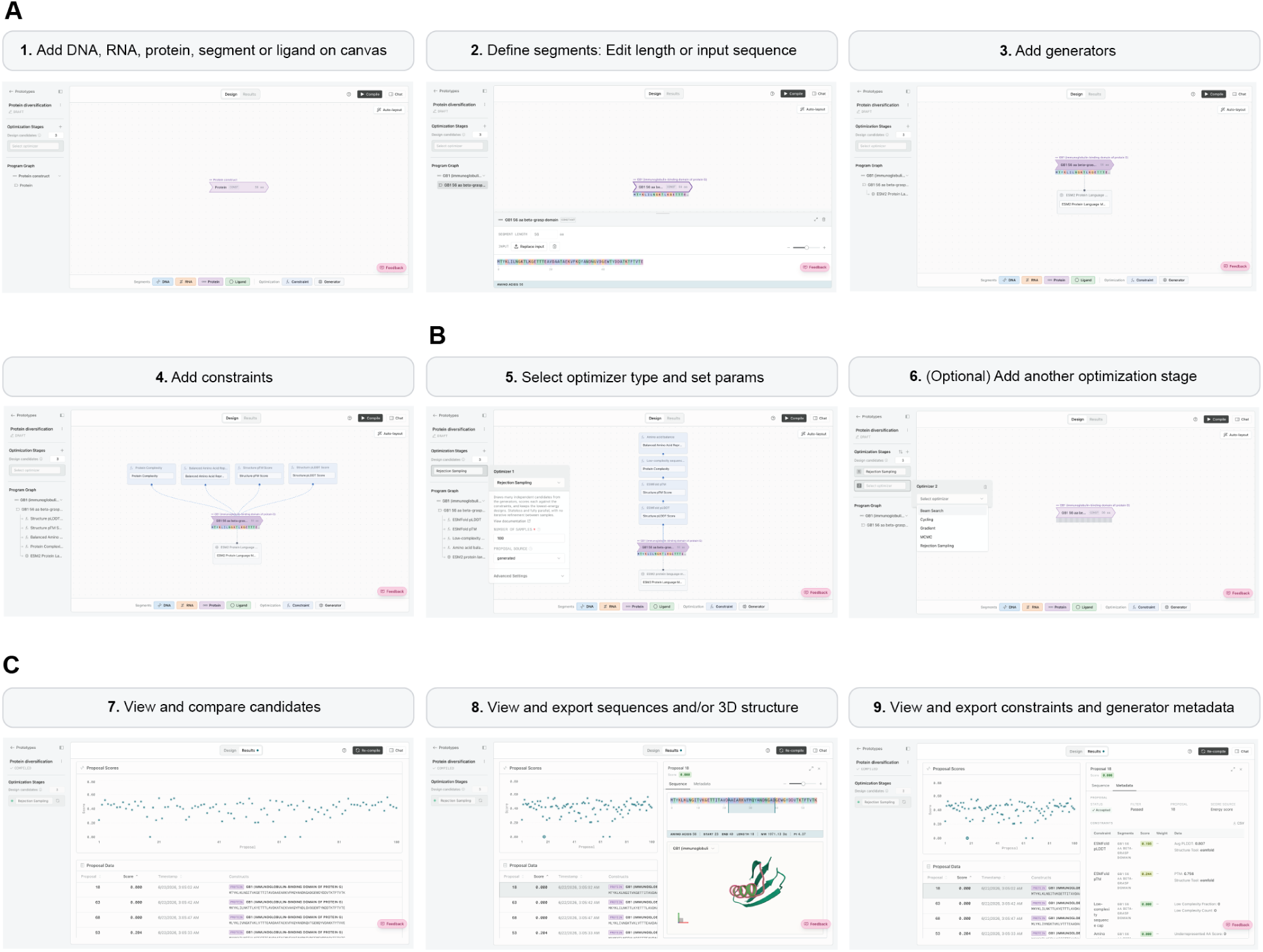
| Workflow in the Proto web application. (**A**) In the web interface, a user first defines sequences and adds generators and constraints. (**B**) The user then selects the optimizer, as well as adds additional optimization stages, and then compiles the program, assuming no configuration or compiler errors. (**C**) The optimizer progress and final results are then visualized alongside tools for inspecting the designed sequences and associated metadata, such as protein structures.

**Figure S6.**
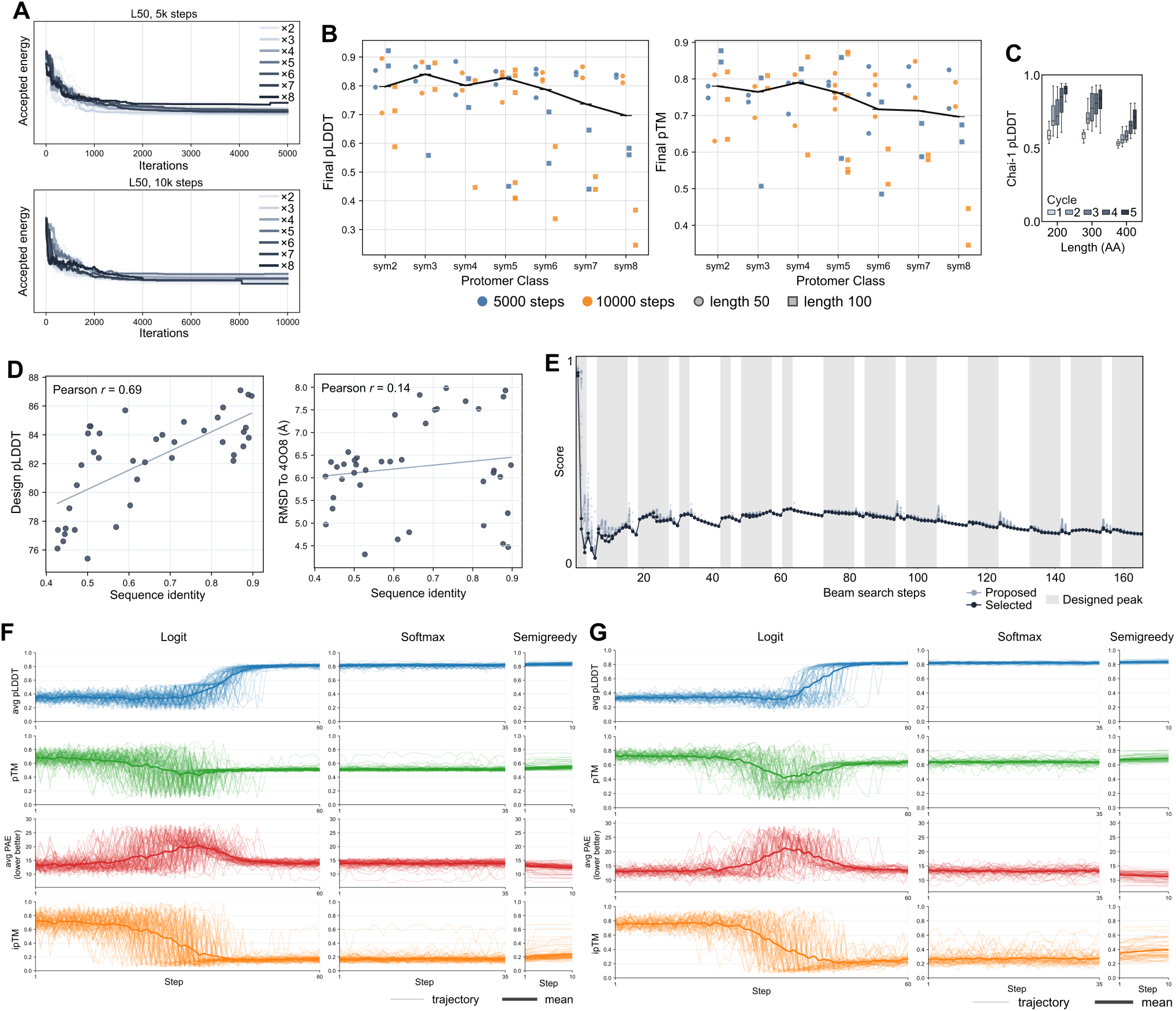
| Additional data on replicating previous design programs. (**A**) Optimization trajectories for symmetric protein simulated annealing grouped by the number of protomers and with separate plots for one set of runs with 5,000 steps and another set of runs with 10,000 steps. The 5,000-step plot is also shown in Figure 2B. (**B**) Jitter plots showing the number of protomers on the horizontal axis and the final ESMFold pLDDT or pTM on the vertical axis. The mean values are also plotted and connected by a black line. (**C**) Protein Hunter was also run with Chai-1 instead of Boltz-2; compare to Figure 2E. (**D**) Scatter plot, where each point corresponds to a designed Cas9 sequence that passes all rejection sampling filters. The protein sequence identity of the designed Cas9 sequence to the closest sequence in the Evo 1 CRISPR-Cas fine-tuning set is plotted on the horizontal axis, with the AF3 pLDDT or the all-atom aligned RMSD between the AF3-predicted Cas9 protein and the native experimental structure on the vertical axis. (**E**) The number of beam search iterations is plotted on the horizontal axis, the Borzoi- and Enformer-defined score is plotted on the vertical axis. Proposed sequences are plotted as light gray dots and the accepted sequence values are plotted as dark gray dots. Lines connect a given iteration to its parent, i.e., corresponding to the sequence from which beam search continued. (**F, G**) Various Germinal trajectories are plotted with AF2-multimer-defined confidence metrics on the vertical axis and the iteration of gradient descent on the horizontal axis. Lighter lines indicate an individual sampling trajectory and the darker line indicates the mean across all trajectories. Sampling runs are shown for VHH (**F**) and scFv (**G**) binders.

**Figure S7.**
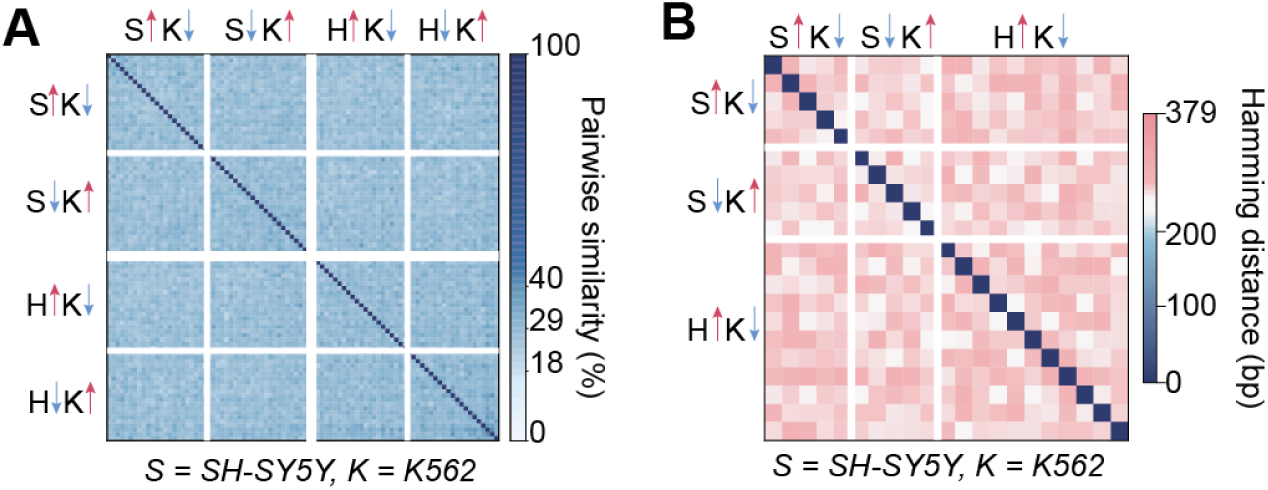
| Pairwise similarities between all selected and differentially spliced ProtoIntrons. (**A**) Pairwise sequence similarity across selected ProtoIntrons grouped by desired differential splicing direction between SH-SY5Y, K562, and HepG2. Darker blue indicates higher sequence similarity. ProtoIntrons remain highly diverse across and within design groups. (**B**) Pairwise Hamming distance across differentially spliced ProtoIntrons. S, SH-SY5Y; K, K562; H, HepG2.

**Figure S8.**
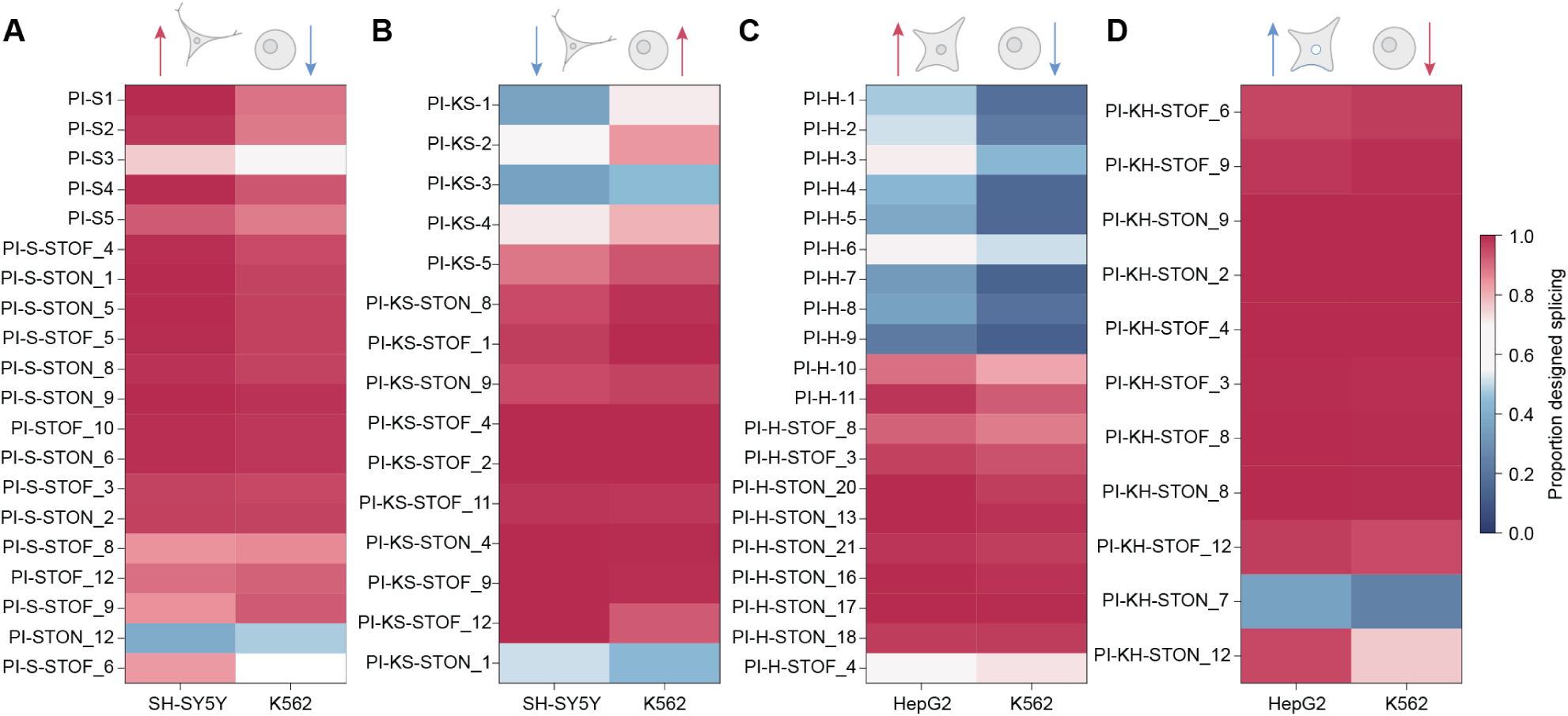
| NGS-derived proportion of designed splicing values for all experimentally screened constructs in SH-SY5Y, HepG2, and K562. Proportion of designed splicing for ProtoIntrons selected to increase designed splicing in (**A**) SH-SY5Y relative to K562, (**B**) K562 relative to SH-SY5Y, (**C**) HepG2 relative to K562, and (**D**) K562 relative to HepG2. 3 out of 4 design campaigns have candidates showing significant differential splicing in the intended design direction.

**Figure S9.**
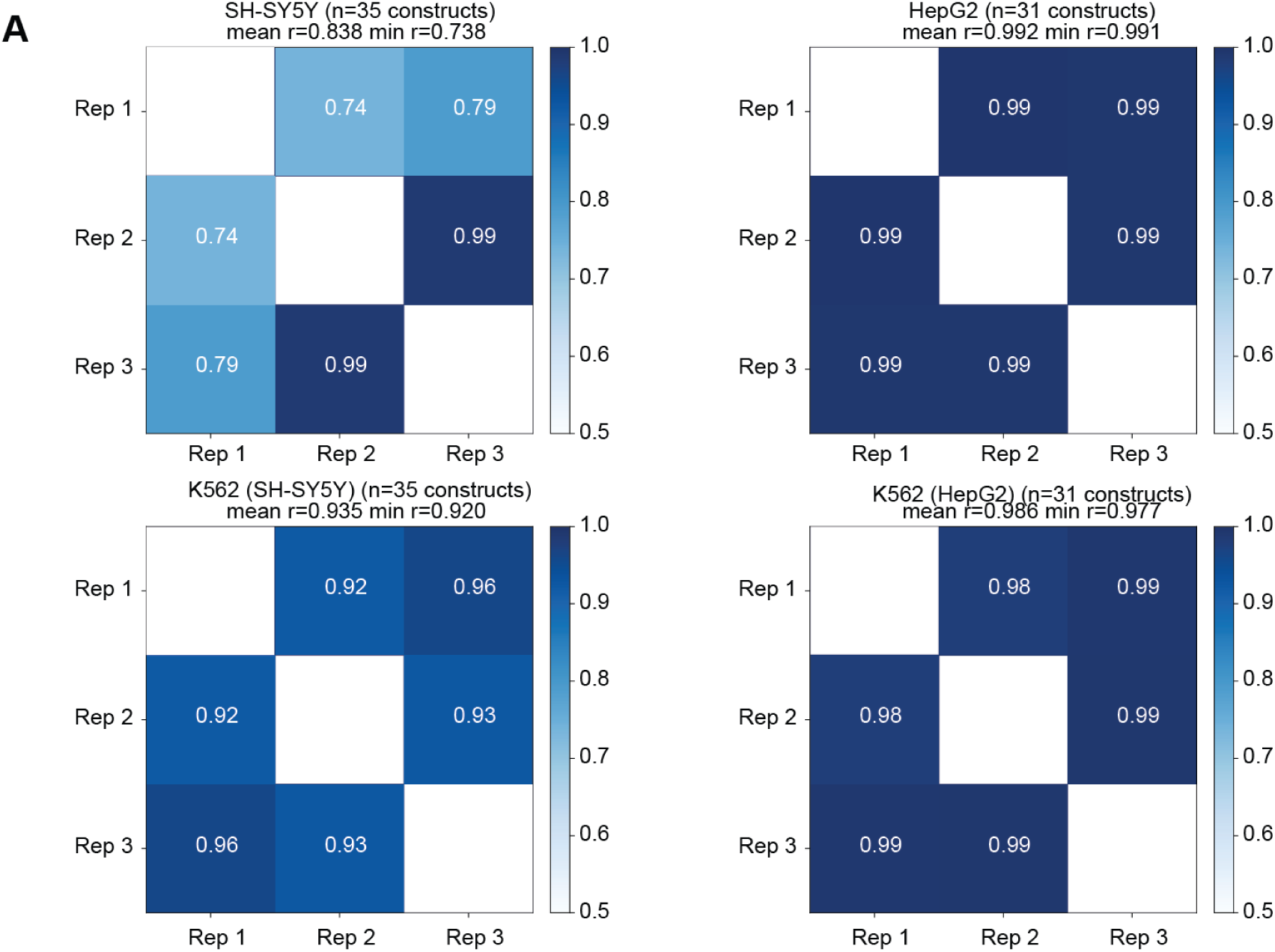
| Sequencing replicate concordance within NGS biological replication batches. (**A**) Pairwise replicate concordance for NGS amplicon sequencing across SH-SY5Y, HepG2, and K562 samples. Heatmaps show pairwise correlations between biological replicates for each cell line and comparison-specific K562 sample.

**Figure S10.**
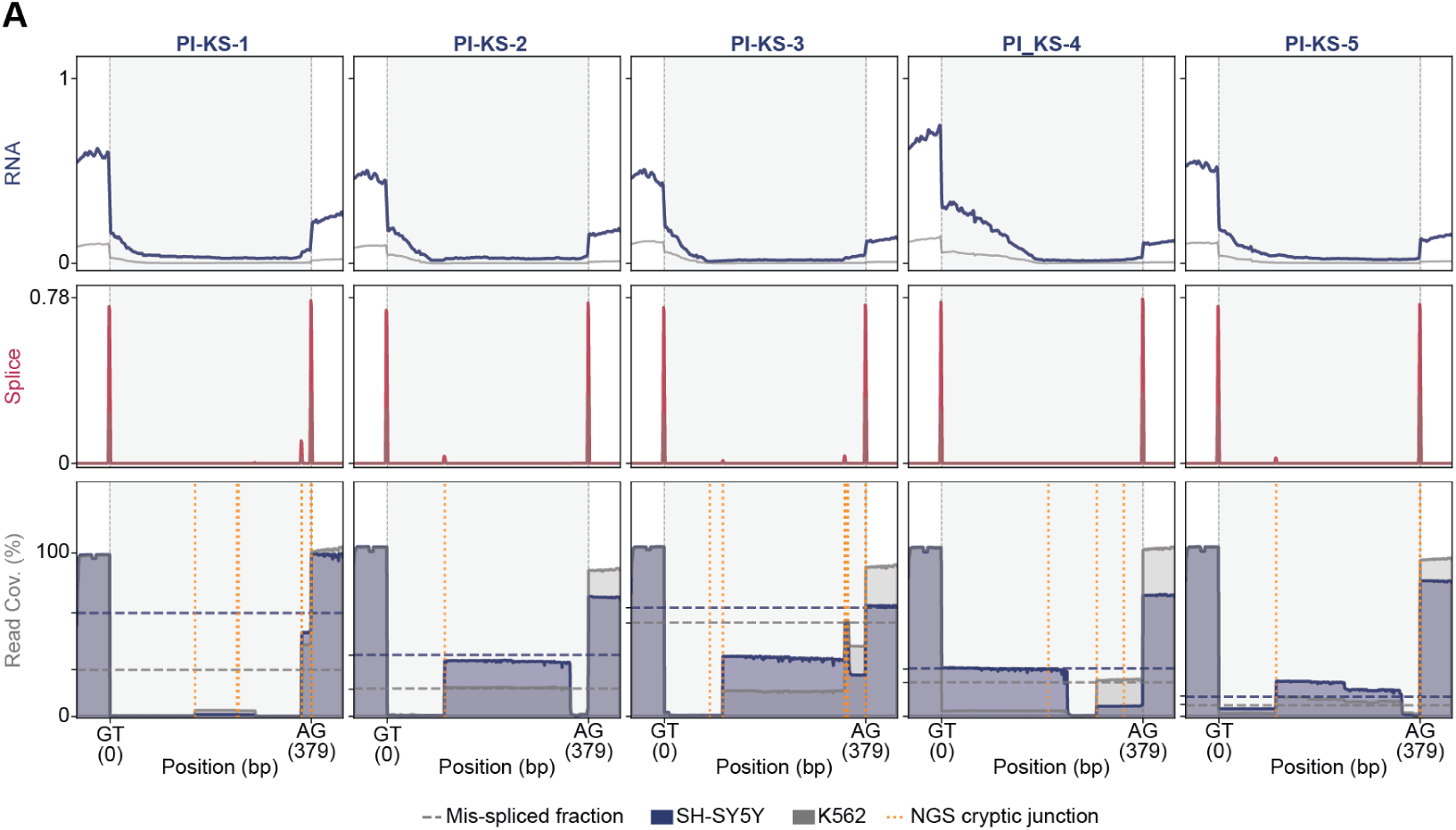
| Amplicon NGS read coverage of differentially spliced ProtoIntrons compared to AlphaGenome predictions for SH-SY5Y low splicing condition. (**A**) RNA, splice-junction, and amplicon NGS read-coverage tracks for ProtoIntrons selected to decrease designed splicing in SH-SY5Y relative to K562. Top and middle tracks show SH-SY5Y-low ProtoIntrons generally show reduced designed and predicted splice-junction usage and increased off-target or cryptic splicing in SH-SY5Y relative to K562.

**Figure S11.**
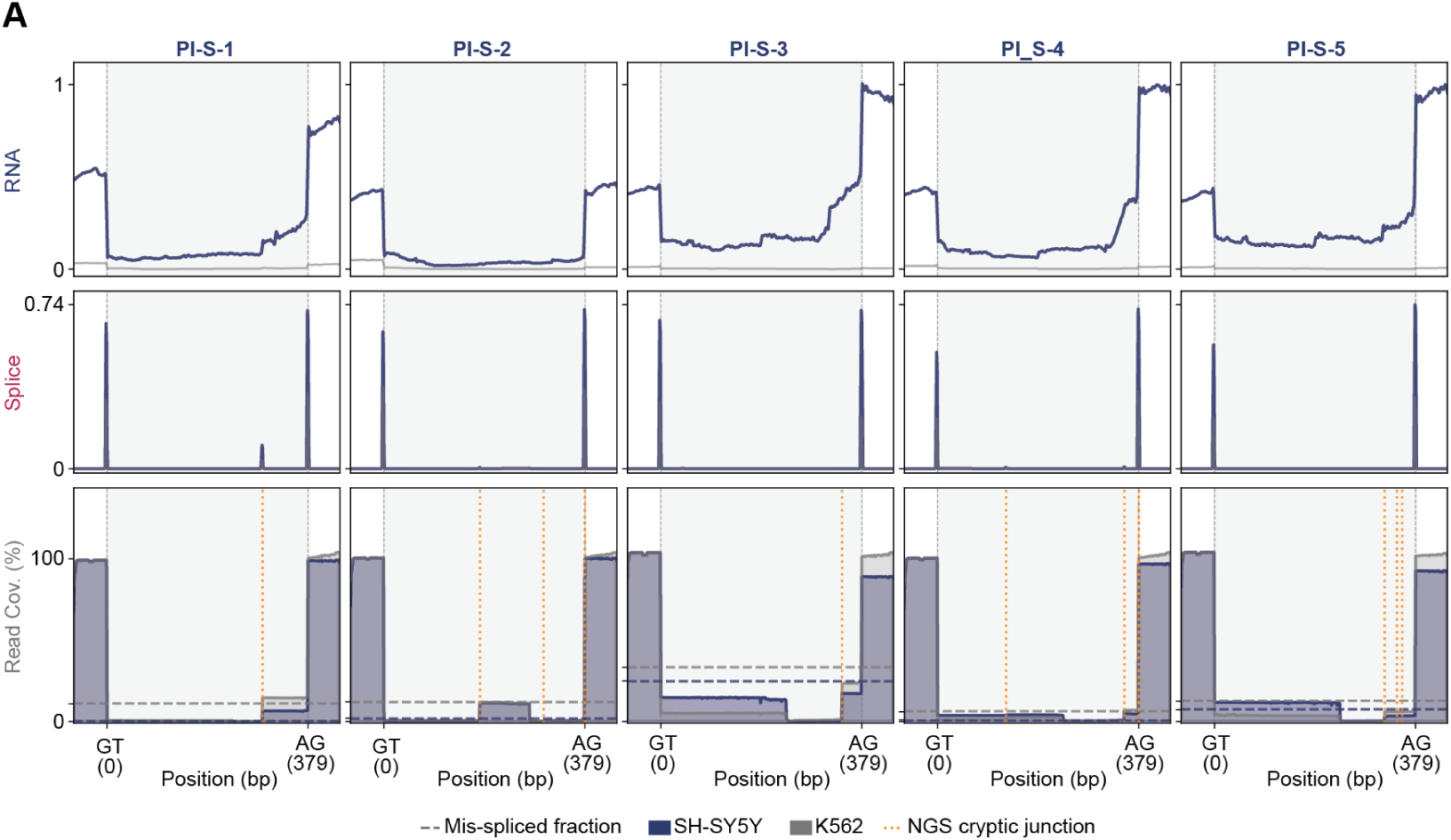
| Amplicon NGS read coverage of differentially spliced ProtoIntrons compared to AlphaGenome predictions for SH-SY5Y high splicing condition. (**A**) RNA, splice-junction, and amplicon NGS read-coverage tracks for ProtoIntrons selected to increase designed splicing in SH-SY5Y relative to K562. Top and middle tracks show AlphaGenome-predicted RNA coverage and splice-junction usage across each ProtoIntron. SH-SY5Y-high ProtoIntrons generally show increased designed and predicted splice-junction usage and reduced off-target or cryptic splicing in SH-SY5Y relative to K562.

**Figure S12.**
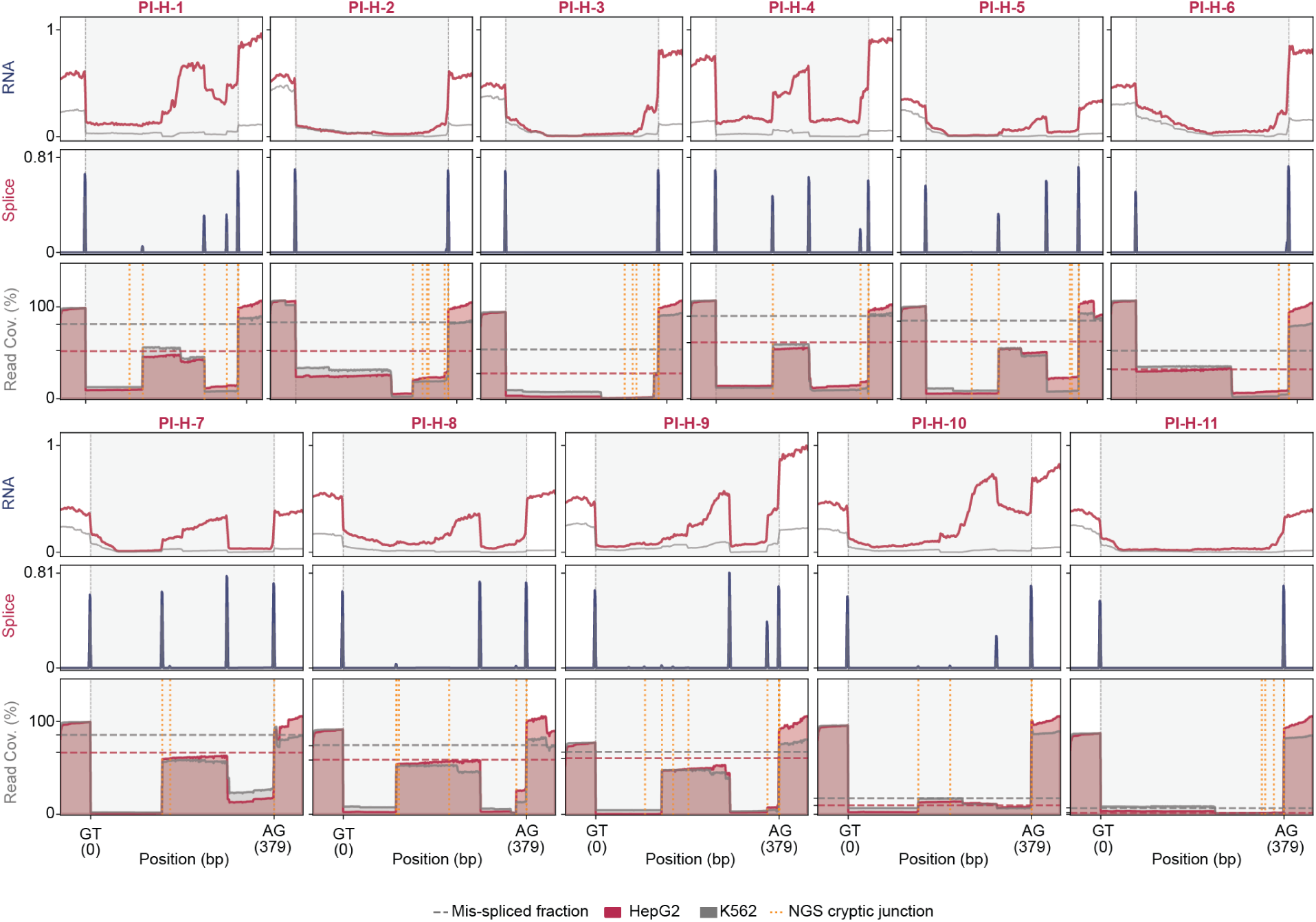
| Amplicon NGS read coverage of differentially spliced ProtoIntrons compared to AlphaGenome predictions for HepG2 high splicing condition. (**A**) RNA, splice-junction, and amplicon NGS read-coverage tracks for ProtoIntrons selected to increase designed splicing in HepG2 relative to K562. Top and middle tracks show AlphaGenome-predicted RNA coverage and splice-junction usage across each ProtoIntron. HepG2-high Pro-toIntrons generally show increased designed splice-junction usage in HepG2 relative to K562, with several constructs also showing cryptic junction usage.

**Figure S13.**
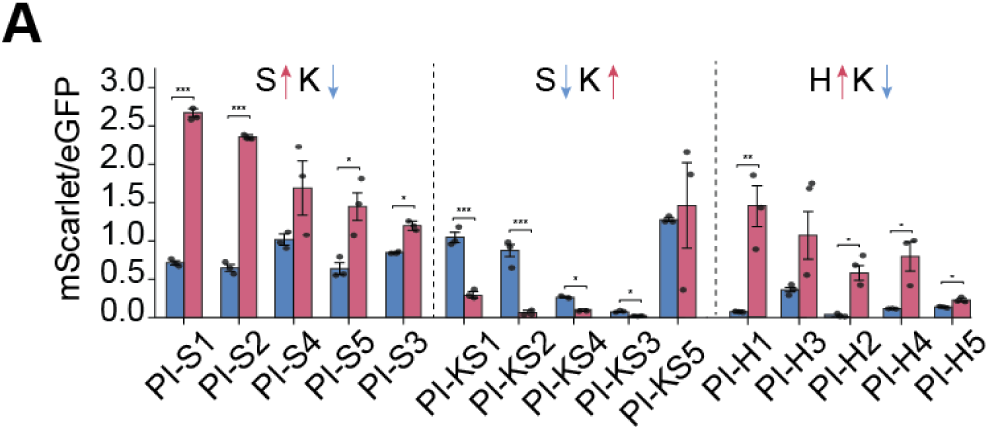
| Full mScarlet differentials for differentially spliced ProtoIntrons as measured by fluorescence microscopy. (**A**) mScarlet/eGFP ratios for differentially spliced ProtoIntrons selected to increase designed splicing in SH-SY5Y relative to K562, K562 relative to SH-SY5Y, and HepG2 relative to K562 as measured by fluorescence microscopy. ProtoIntrons generally produce the intended cell-line-specific fluorescence differences across selected design groups. Bars, mean; circles, individual biological replicates; error bars, standard error of mean. *, *P* < 0.05; **, *P* < 0.01; ***, *P* < 0.001.

**Figure S14.**
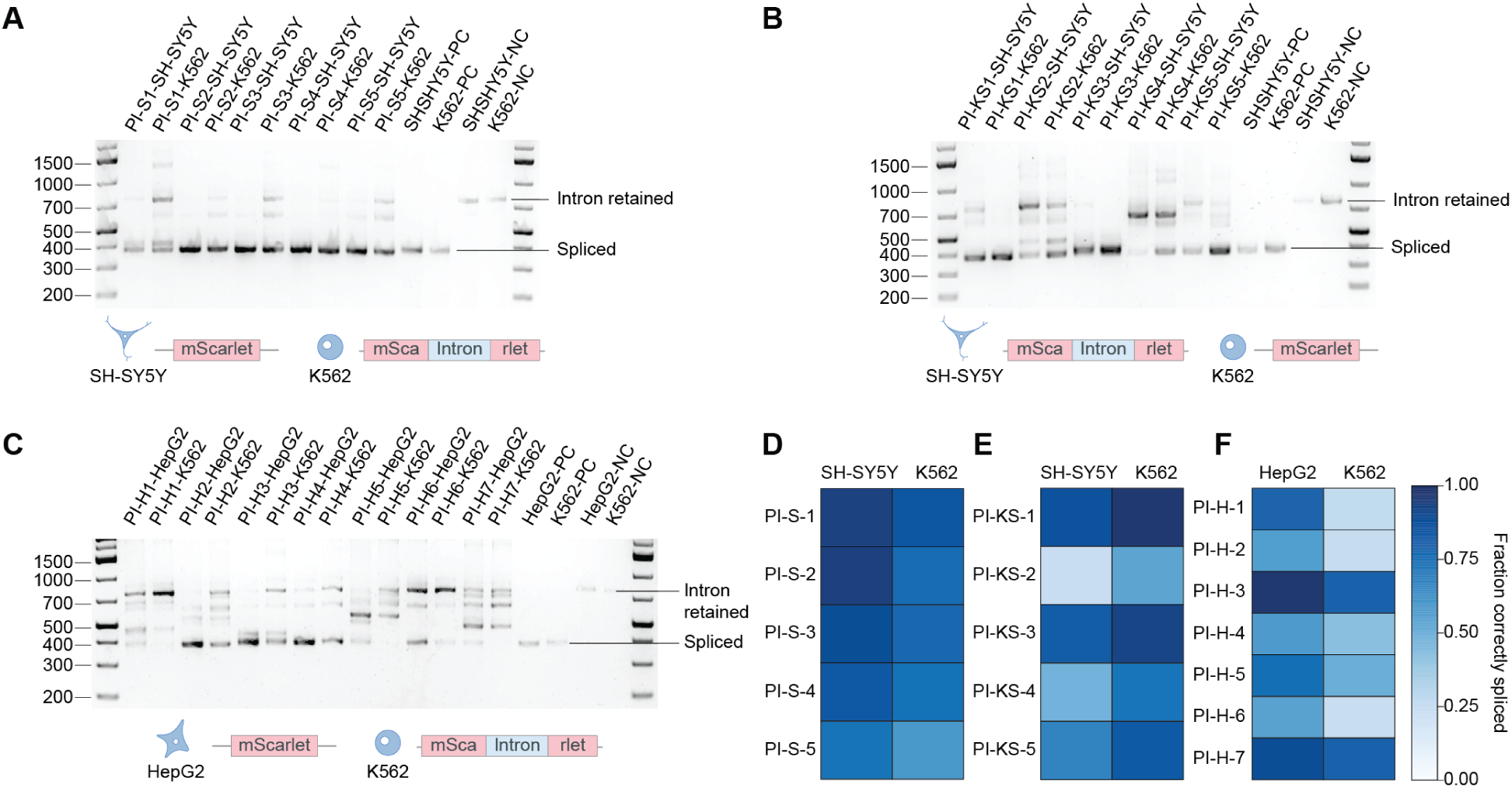
| Agarose gel electrophoresis and nanopore sequencing of PCR products from differentially spliced ProtoIntron constructs. (**A**–**C**) Agarose gel analysis of RT-PCR products from ProtoIntrons selected to increase designed splicing in (**A**) SH-SY5Y relative to K562, (**B**) K562 relative to SH-SY5Y, and (**C**) HepG2 relative to K562. Differentially spliced ProtoIntrons generally produce the expected shift between spliced and intron-retained products across the paired cell lines. PC, HBB2c intron positive control; NC, reversed HBB2c intron negative control. (**D**–**F**) Fraction correctly spliced reads as measured by nanopore sequencing for (**D**) SH-SY5Y-correct/K562-mis-spliced, (**E**) SH-SY5Y-mis-spliced/K562-correct, and (**F**) HepG2-correct/K562-mis-spliced conditions.

**Figure S15.**
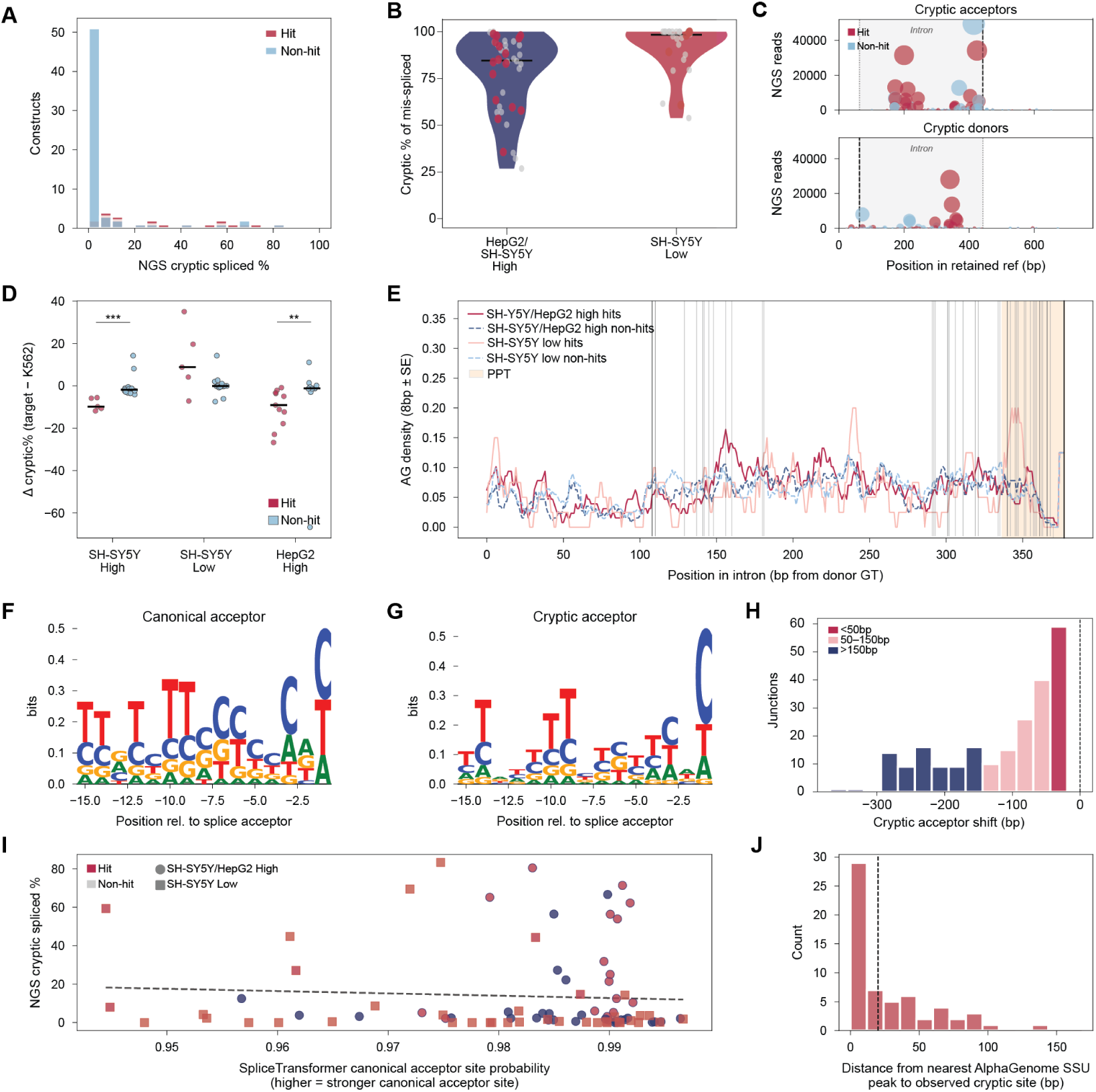
| Characterization of cryptic splicing in ProtoIntrons. (**A**) Distribution of NGS-measured cryptic spliced fraction across screened ProtoIntrons, stratified by experimentally successful hits and non-hits. (**B**) Fraction of mis-spliced reads attributable to cryptic splicing for SH-SY5Y/HepG2-high and SH-SY5Y-low design groups. Cryptic splicing accounts for a substantial fraction of mis-spliced reads, particularly in SH-SY5Y-low designs. (**C**) Positions and read counts of cryptic acceptor and donor junctions detected by NGS relative to the retained-reference sequence. Cryptic acceptors are more frequent than cryptic donors and occur predominantly within the designed intron upstream of the canonical acceptor. (**D**) Difference in cryptic splicing between the target cell line and K562 for each design group. Successful high-splicing hits generally show reduced cryptic splicing in the target cell line relative to K562, whereas SH-SY5Y-low hits show increased cryptic splicing in SH-SY5Y. (**E**) AG dinucleotide density across the designed intron for hits and non-hits, with the polypyrimidine tract (PPT) highlighted. Cryptic splice usage occurs in regions with local AG enrichment. (**F, G**) Sequence logos of canonical and cryptic acceptor sites. Cryptic acceptors retain core splice-acceptor features but show weaker consensus than canonical acceptors. (**H**) Distribution of positional shifts from canonical to cryptic acceptor sites. Most cryptic acceptors are located upstream of the canonical acceptor, often within 150 bp. (**I**) NGS-measured cryptic spliced fraction versus SpliceTransformer canonical acceptor-site probability. Cryptic splicing is largely independent of predicted canonical acceptor strength. (**J**) Distance from the nearest AlphaGenome splice-site-usage peak to the observed cryptic site. Most cryptic sites occur near AlphaGenome-predicted splice-site-usage peaks.

**Figure S16.**
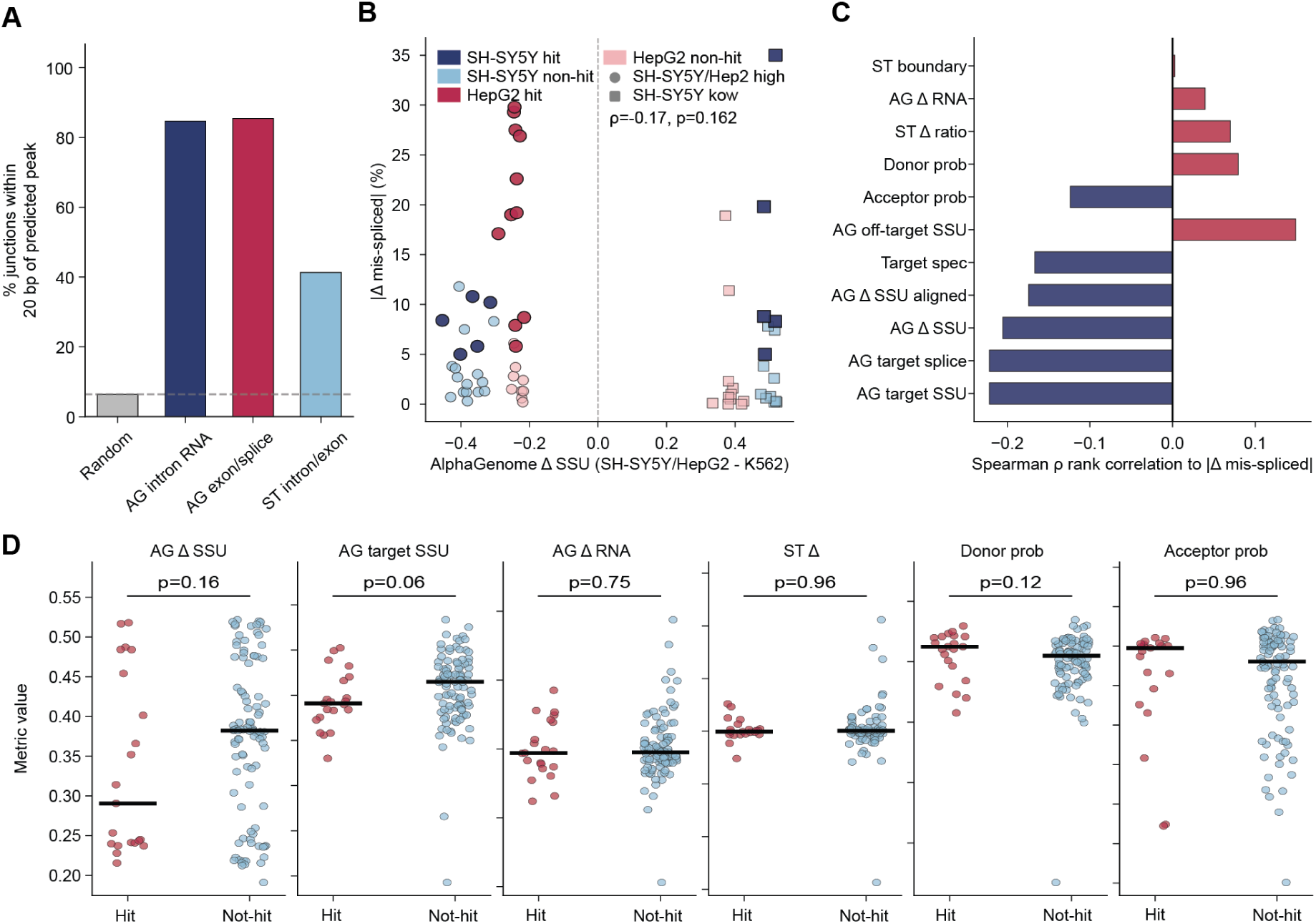
| Predictive power of AlphaGenome and SpliceTransformer constraints on predicting experimental splicing outcomes. (**A**) Fraction of observed splice junctions located within 20 bp of a predicted junction for random positions, AlphaGenome intronic RNA peaks, AlphaGenome splice-site-usage peaks, and SpliceTransformer intron/exon predictions. AlphaGenome splice-site-usage and intronic RNA predictions capture most observed junctions. (**B**) Change in mis-splicing between the target cell line and K562 versus AlphaGenome-predicted differential splice-site usage for experimentally screened ProtoIntrons. Points indicate individual ProtoIntrons, stratified by design group and hit status. (**C**) Spearman rank correlation between individual *in-silico* design metrics and absolute differential mis-splicing. No single metric strongly predicts experimental success across designs. (**D**) Comparison of selected AlphaGenome and SpliceTransformer metric values between hits and non-hits. Individual metrics do not significantly separate successful from unsuccessful ProtoIntrons.

**Figure S17.**
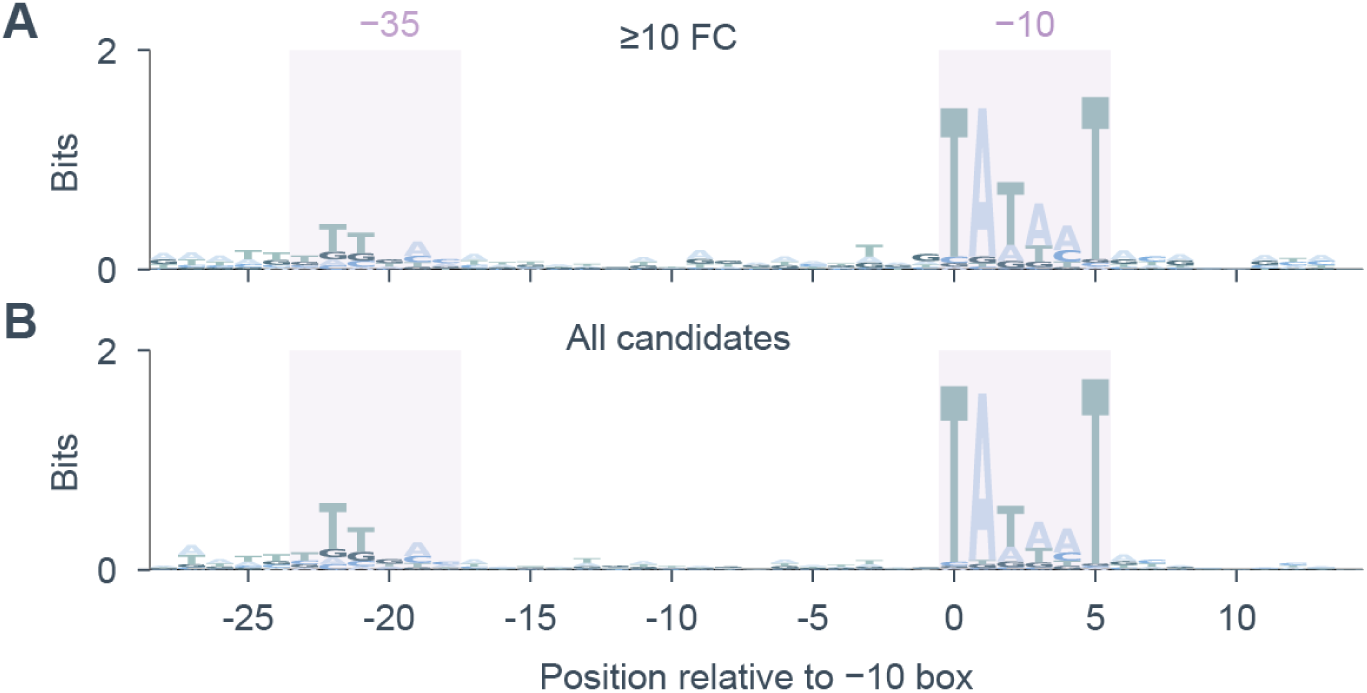
| ProtoPromoter sequence logo plots. (**A, B**) Sequence logos for (**A**) ProtoPromoters with ≥10-fold eGFP induction and (**B**) all selected ProtoPromoter candidates, aligned relative to the −10 box. Shaded regions indicate the expected positions of the −35 and −10 promoter elements. Highly active ProtoPromoters show slightly less conservation of σ^70^-like −10 and −35 box sequence, while the −35 region is generally more weakly conserved across candidates.

**Figure S18.**
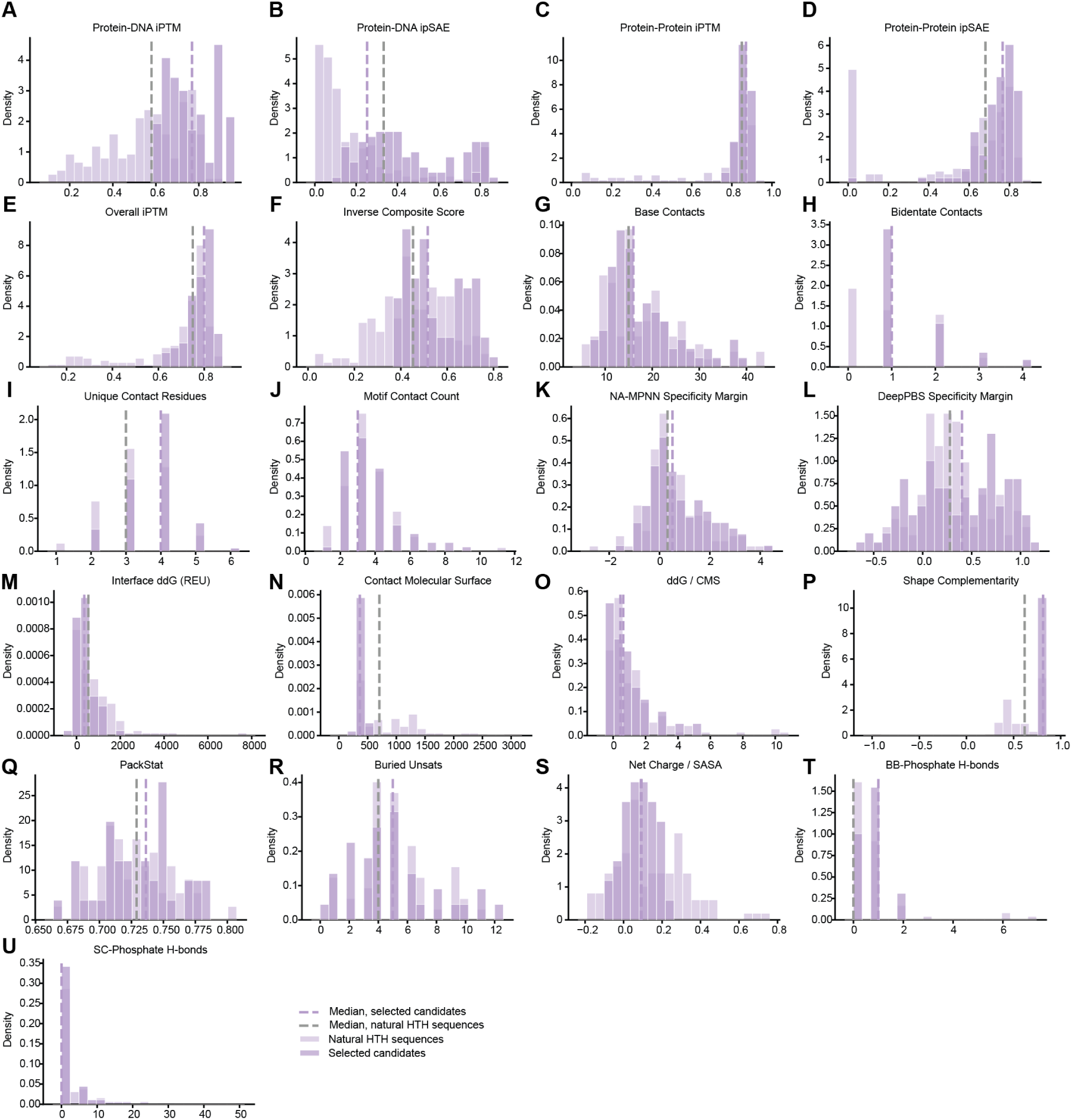
| Comparison of constraint value distribution among selected repressor designs compared to natural DNA-binding sequences. (**A–U**) Distributions of *in-silico* design metrics for selected ProtoRepressor candidates compared to natural helix-turn-helix (HTH) DNA-binding sequences, including (**A**) protein-DNA ipTM, (**B**) protein-DNA ipSAE, (**C**) protein-protein ipTM, (**D**) protein-protein ipSAE, (**E**) overall ipTM, (**F**) inverse composite score, (**G**) base contacts, (**H**) bidentate contacts, (**I**) unique contact residues, (**J**) motif contact count, (**K**) NA-MPNN specificity margin, (**L**) DeepPBS specificity margin, (**M**) interface ddG, (**N**) contact molecular surface, (**O**) ddG normalized by contact molecular surface, (**P**) shape complementarity, (**Q**) PackStat, (**R**) buried unsatisfied polar atoms, (**S**) net charge normalized by solvent-accessible surface area, (**T**) backbone-phosphate hydrogen bonds, and (**U**) side-chain-phosphate hydrogen bonds. Dashed vertical lines indicate distribution medians for selected candidates and natural HTH sequences. Selected ProtoRepressors generally show similar or improved predicted complex confidence, protein-DNA contact geometry, and specificity metrics relative to natural HTH sequences, while maintaining comparable interface-quality and surface-chemistry properties.

**Figure S19.**
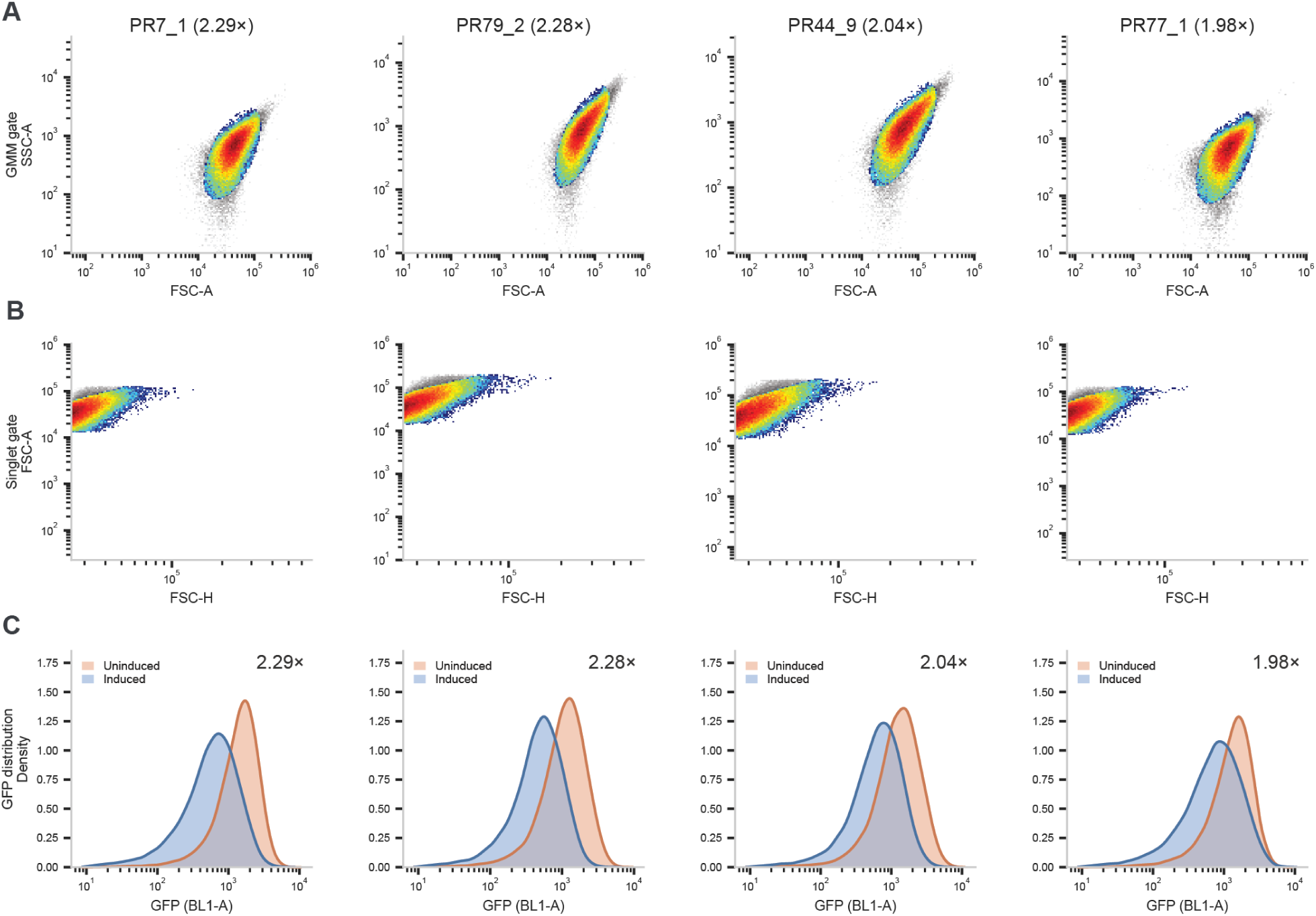
| Example gating strategy for flow cytometry analysis of ProtoRepressors. (**A**) Forward scatter area (FSC-A) versus side scatter area (SSC-A) for representative uninduced wells. A Gaussian mixture model fit in log FSC-A/SSC-A space was used to isolate the live-cell population (pseudocolor); excluded events are shown in grey. (**B**) Representative FSC-A versus FSC-H of GMM-gated events, used to exclude doublets via a median absolute deviation filter on the FSC-A/FSC-H ratio (grey = excluded doublets, pseudocolor = retained singlets). (**C**) Representative GFP fluorescence (BL1-A) distributions for uninduced (red) and repressor-induced (blue) cells after applying both gates. Fold repression (geometric mean across all replicates) is annotated for each candidate.

**Figure S20.**
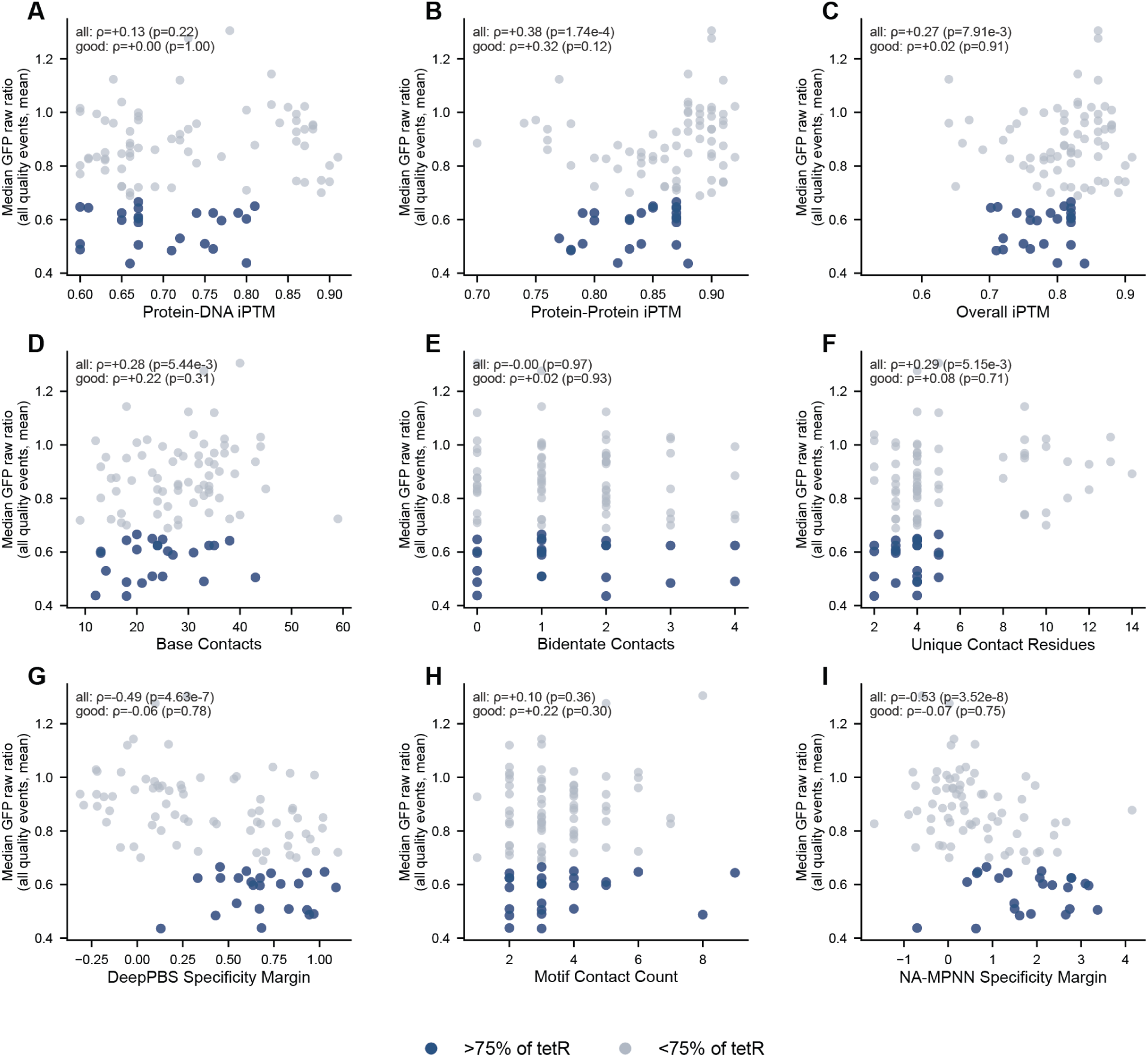
| Correlation between repressor constraints and experimental success rates among promoter-repressor pairs. Fold repression across screened promoter-repressor pairs plotted against individual *in-silico* Pro-toRepressor design metrics, including protein-DNA ipTM, protein-protein ipTM, overall ipTM, NA-MPNN specificity margin, DeepPBS specificity margin, motif contact count, base contacts, bidentate contacts, and unique contact residues. Points correspond to individual promoter-repressor pairs and are colored by repression strength relative to the indicated fold-repression threshold. Several metrics show modest associations with repression, including specificity margins, protein-protein complex confidence, and overall ipTM. However, no single metric cleanly separates strongly repressive from weakly repressive promoter-repressor pairs.

**Figure S21.**
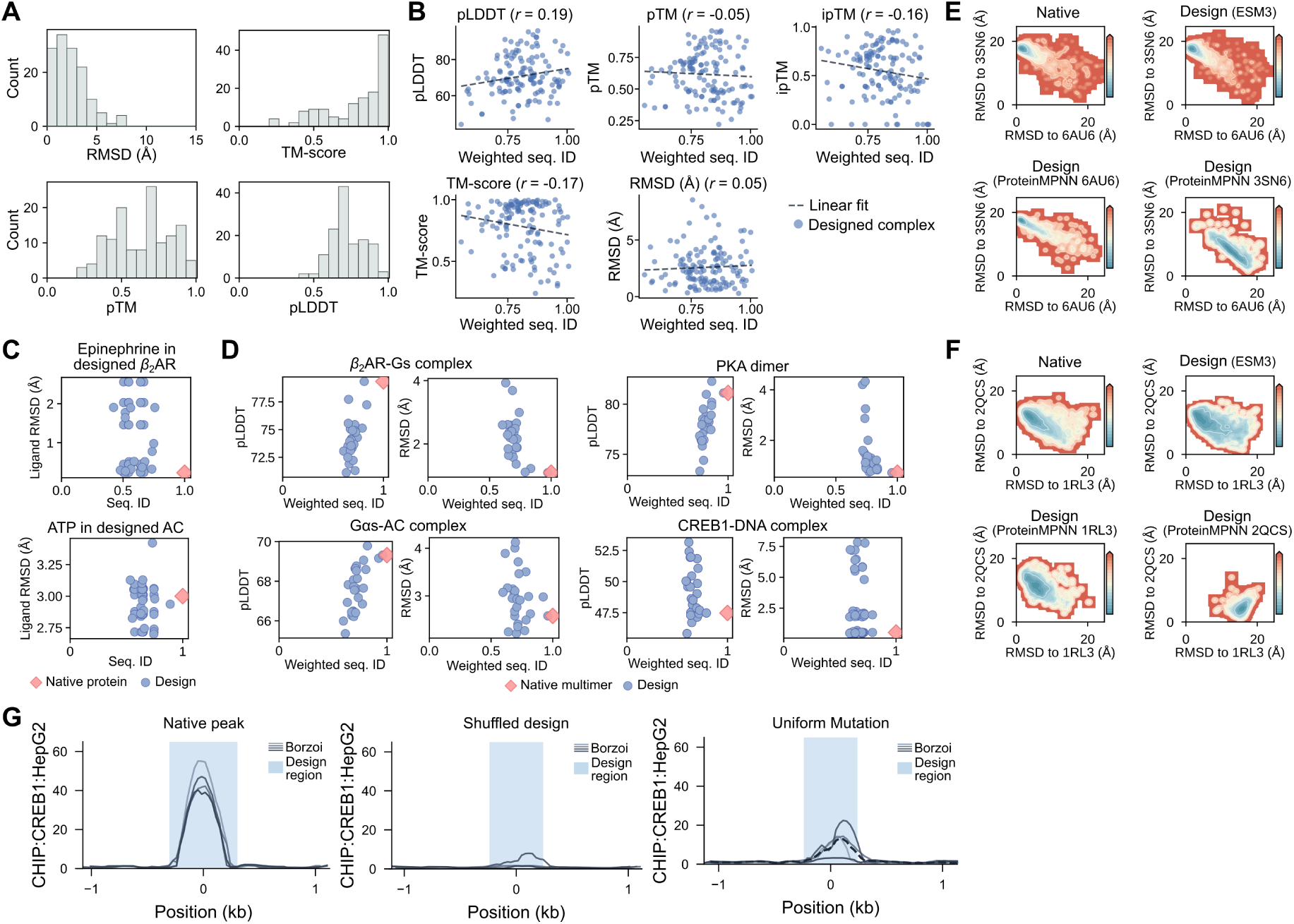
| Additional data on large-scale diversification tasks. (**A**) Distributions of AF3 multimeric predictions on native sequences, including the RMSD of the AF3-predicted native complex to the native experimental structure (median 1.5 Å), AF3 TM-score to native (median 0.89), as well as AF3 structural confidence metrics pTM (median 0.68) and pLDDT (median 68). Compare to the distributions for ESM3-diversified AF3 predictions in Figure 5C. (**B**) Scatter plots where each point corresponds to a different designed multimer. On the horizontal axis is the sequence identity of a designed complex compared with the native complex, followed by taking the average over all unique sequences in the complex and weighted by sequence length (“Weighted seq. ID”). On the vertical axis are the AF3-predicted confidence scores (pLDDT, pTM, ipTM) or structural similarity metrics of the AF3-predicted structures compared to the corresponding experimental structures (TM-score and RMSD). A least-squares-fit regression line and the Pearson correlation (*r*) are also visualized. (**C**) Scatter plots where each point corresponds to a different designed variant of a given protein chain. The native protein chain is plotted as a red diamond. On the horizontal axis is the sequence identity of ESM3-diversified *β*2AR or adenylyl cyclase compared to the native sequence. On the vertical axis is the pocket-aligned RMSD between the AF3-docked epinephrine or ATP molecules and the experimentally determined molecule. (**D**) Scatter plots where each point corresponds to a different designed variant of a given multimeric complex in the diversified signaling pathway. The native multimer complex is plotted as a red diamond. On the horizontal axis is the weighted protein sequence identity as described in (**B**). On the vertical axis is the AF3-predicted pLDDT or the all-atom aligned RMSD between the AF3-predicted structure and the experimental structure of that complex. (**E, F**) Heatmaps of BioEmu ensembles related to G*α*s (**E**) and the PKA regulatory subunit (**F**). The colorbar indicates the density of conformations, where blue indicates high density and red indicates low density. The top left of each panel shows the BioEmu-predicted ensemble of the native sequence. The top right shows the BioEmu-predicted ensemble of the best designed sequence (the sequence within the highest confidence AF3-predicted multimer structure); replicated in Figure 5G,**I**. The bottom of each panel shows the BioEmu-predicted ensemble of conformationally biased sequences using low temperature sampling from ProteinMPNN. (**G**) Borzoi-predicted CREB1 ChIP-seq signal over (from left to right) a native CREB1 peak transplanted into the genetic context, a GC-matched random sequence formed by permuting the sequence of the Evo 2-generated design, and a sequence formed by replacing Evo 2 with a uniform mutation generator.

**Figure S22.**
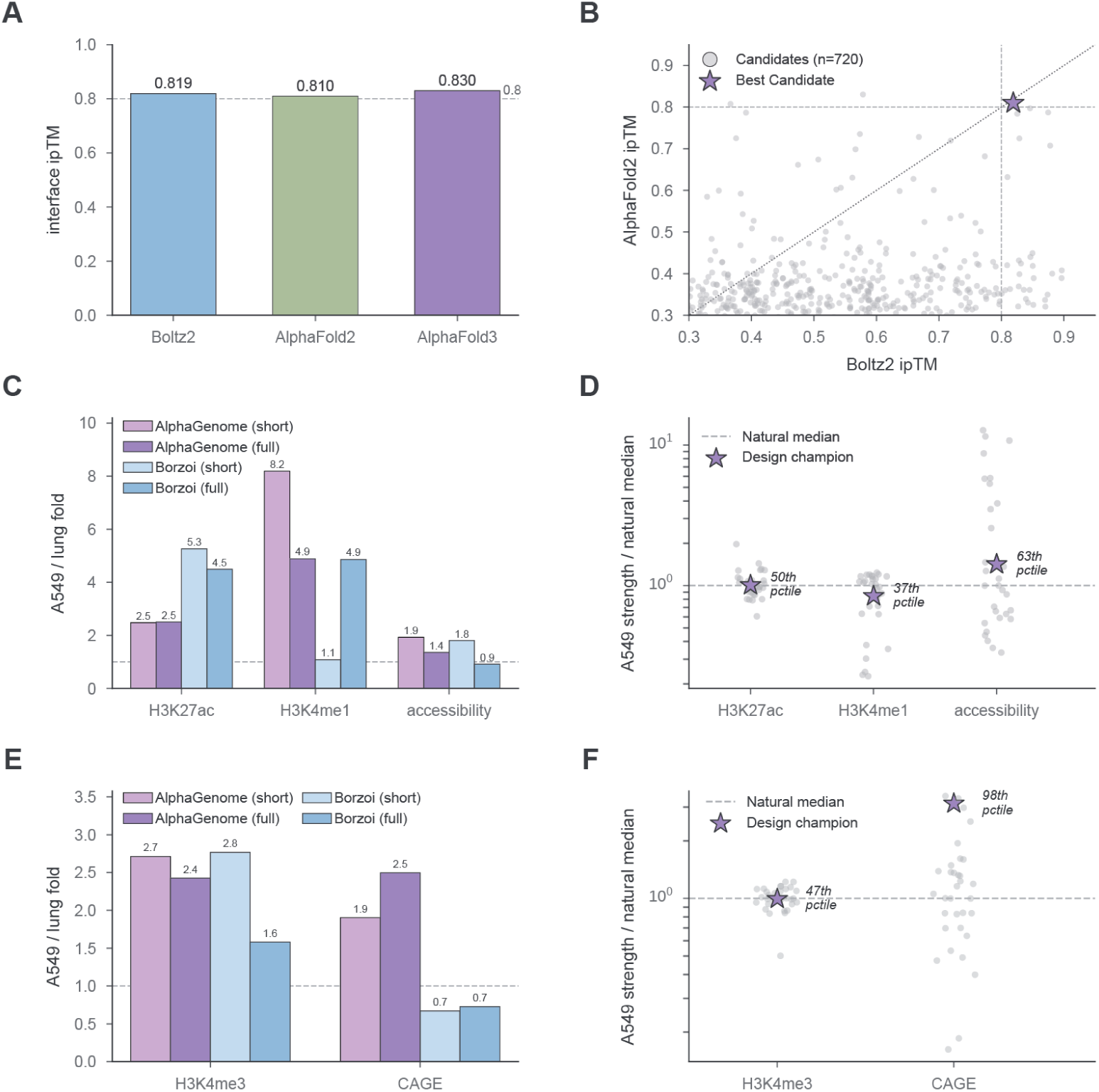
| Agreement of the designed NSCLC binder and regulatory cassette with multiple structure-prediction models, sequence-to-function models, and natural regulatory sequences. (**A**) ipTM of the lowest energy EGFR binder under three independent structure-prediction oracles shows agreement across all three families. (**B**) Boltz-2 versus AlphaFold 2 ipTM for the initial binder candidate pool; the champion sits in the high-confidence consensus corner (upper right), whereas most candidates pass one oracle but fail the other. (**C**) A549/lung chromatin fold for the designed enhancer at two active integration loci (GAPDH, EEF1A1; geometric mean), for AlphaGenome and Borzoi, comparing fold change in shorter scoring context (16 kb of integration site context) to maximal genomic context (524-kb window) for H3K27ac, H3K4me1, and accessibility (ATAC for Al-phaGenome, DNase for Borzoi); dashed line at fold = 1. Both models retain A549-selective enhancer chromatin at full genomic context, suggesting the selectivity is not an artifact of the short scoring window. (**D**) Best designed enhancer versus natural enhancers on Borzoi-predicted A549 signal at the active loci, expressed as signal normalized to the per-mark natural median. (**E**) A549/lung fold for the best designed 100-nt promoter at the active integration loci (GAPDH, EEF1A1; geometric mean), for AlphaGenome and Borzoi, comparing the production short scoring context (16 kb of integration site context) to full maximal genomic context (524-kb window), for H3K4me3 and CAGE-predicted transcription initiation. At full genomic context, H3K4me3 is A549-favored in both models while CAGE is A549-favored for AlphaGenome but not A549-selective for Borzoi. (**F**) Best designed promoter versus natural promoters on Borzoi-predicted absolute A549 signal at the active loci, expressed as signal normalized to the per-mark natural median. Best candidate’s A549 H3K4me3 falls within the natural-promoter range, while its A549 CAGE signal is high compared to the natural set.

**Figure S23.**
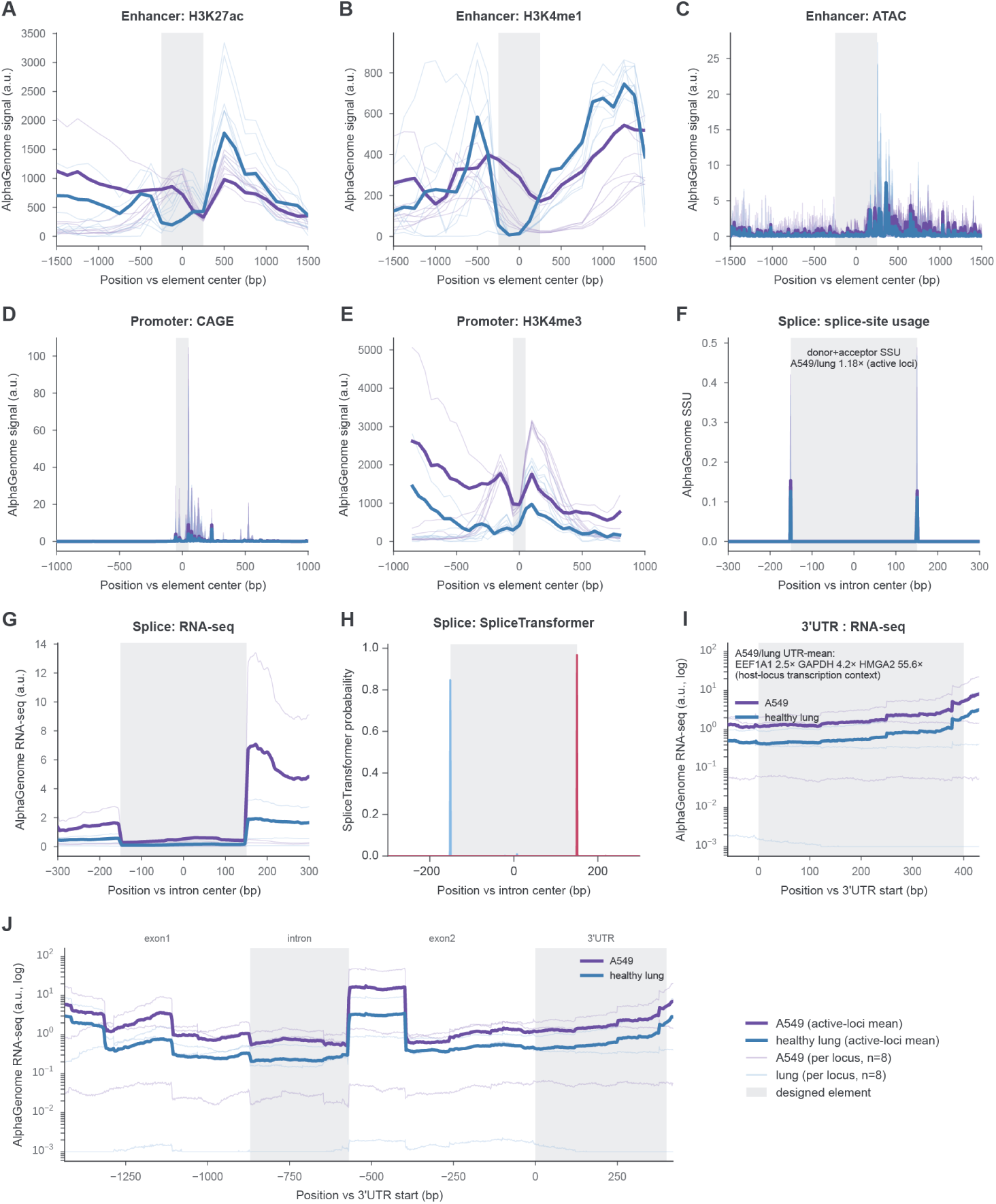
| Per-position regulatory-track predictions for designed NSCLC-selective regulatory cassette across various integration contexts. All tracks are AlphaGenome predictions for A549 lung adenocarcinoma (EFO:0001086, purple) versus healthy lung (UBERON:0002048, blue). (**A–C**) Chromatin signal for the best designed 500-nt enhancer embedded in its generation cassette across all integration loci (GAPDH, ACTB, EEF1A1, FTL, and HMGA2) (**A**) H3K27ac, (**B**) H3K4me1, and (**C**) chromatin accessibility (ATAC). Activities were compared within the gray interval. Signal downstream of this interval reflects the HSV-TK exon 1 region, which showed high predicted ATAC-seq activity invariant of context. (**D, E**) Chromatin signal for the best designed 100-nt promoter across the same eight loci: (**D**) CAGE-predicted transcription initiation and (**E**) H3K4me3. (**F–H**) Per-position AlphaGenome predictions for the best designed 301-nt synthetic intron within the HSV-TK exon1-intron-exon2 transgene, across the three integration loci with available healthy-lung tracks (GAPDH, EEF1A1, HMGA2), zoomed to the intron ± 300 bp. (**F**) SSU, which is sharply localized to the donor (intron 5^′^) and acceptor (intron 3^′^) spikes flanking the spliced-out intron body, annotated with the donor and acceptor site locations. (**G**) RNA-seq coverage, with the peak just past the intron 3^′^ edge corresponding to the downstream exon-junction signal. (**H**) SpliceTrans-former donor and acceptor strength predictions for designed intron. (**I**) AlphaGenome RNA-seq coverage for the best designed 400-nt 3^′^UTR off-switch across GAPDH, EEF1A1, and HMGA2. (**J**) RNA-seq coverage over the full transgene, showing all components of the expressed HSV-TK segment in lung tissue and A549.

**Table S1.** | Parameter sweeps and number of samples for intron designs. We conducted sweeps with different numbers of sampled sequence that explored different parameters (initial sequence, intron length, SpliceTrans-former tissue specificity guidance) for K562 on/off and with either HepG2 or SH-SY5Y as the complementary on/off cell line. As AlphaGenome did not directly have an SH-SY5Y output, we used a combination of neural cell predictions as a proxy (**Methods**).

| Target condition | Off-target condition | Initialization | Intron length (bp) | SpliceTransformer specificity | Final samples |
| --- | --- | --- | --- | --- | --- |
| Neural/SH-SY5Y | K562 | HBB 2 chimera | 379 | Off | 355 |
| Neural/SH-SY5Y | K562 | HBB 2 chimera | 379 | On | 657 |
| Neural/SH-SY5Y | K562 | Random | 151 | On | 27 |
| Neural/SH-SY5Y | K562 | Random | 301 | On | 100 |
| HepG2 | K562 | HBB 2 chimera | 379 | Off | 600 |
| HepG2 | K562 | HBB 2 chimera | 379 | On | 600 |
| HepG2 | K562 | Random | 151 | On | 33 |
| HepG2 | K562 | Random | 301 | Off | 50 |
| HepG2 | K562 | Random | 301 | On | 50 |
| K562 | HepG2 | HBB 2 chimera | 379 | Off | 100 |
| K562 | HepG2 | HBB 2 chimera | 379 | On | 100 |
| K562 | Neural/SH-SY5Y | HBB 2 chimera | 379 | Off | 100 |
| K562 | Neural/SH-SY5Y | HBB 2 chimera | 379 | On | 100 |
| Total |  |  |  |  | 2,872 |

**Table S2.** | Filters, thresholds, and acceptance criteria across the ProtoRepressor design pipeline. Hard filters, gate thresholds, and their favorable directions are listed for each stage, from initial candidate sourcing through Stage 3 MCMC refinement and final candidate selection. Thresholds operate on raw metric values; the [0, 1] framework scaling used inside the Stage 3 objective (0 is best) is independent of these gates and is summarized separately in the objective-weights (**Methods**). Calibrated Rosetta prefilter criteria (filter_eq/filter_cut) follow Glasscock et al. (2025).

| Stage | Metric | Direction (raw value) | Threshold |
| --- | --- | --- | --- |
| Stage 1 candidate filter | HMM sequence E-value | lower better | $\leq 5 \times 10^{-3}$ |
|  | Length | window | 80–200 aa |
| | Low complexity | lower better | $\leq 0.3$ |
| | Repetitiveness | lower better | $\leq 0.3$ |
| | Diversity | higher better | $\geq 0.3$ |
| | Energy score (weighted avg; scaled [0, 1]) | lower better | $\leq 0.3$ |
| Stage 1 co-fold gate (Boltz-2) | Overall ipTM | higher better | $\geq 0.5$ |
| | Protein-DNA ipTM | higher better | $\geq 0.5$ |
| | Protein-protein ipTM | higher better | $\geq 0.5$ |
| | Protein pLDDT | higher better | $\geq 70$ |
| Stage 2 LigandMPNN acceptance | Protein-DNA base contacts (4.0 Å) | higher better | $\geq 1$ |
|  | Calibrated Rosetta filter (interface ddG, CMS) | — | filter_eq/filter_cut (Glasscock et al., 2025) |
| Stage 2 AF3 cascade gate | Protein-DNA ipTM | higher better | $\geq 0.6$ |
| | Overall ipTM | higher better | $\geq 0.6$ |
| | Protein-protein ipTM | higher better | $\geq 0.7$ |
| | Protein pLDDT | higher better | $\geq 70$ |
| Stage 2 Rosetta rescore | CMS | higher better | $\geq 225 \text{ Å}^2$ |
| | interface_sc | higher better | $\geq 0.60$ |
| | packstat | higher better | $\geq 0.55$ |
| | base_score | higher better | $\geq 10$ |
| | bidentate_score | higher better | $\geq 1.0$ |
| | Buried-unsatisfied polar atoms | lower better | $\leq 2.0$ |
| Stage 3 motif gate | Protein-DNA contacts / unique contacting residues / unique contacted bases (4.0 Å) | higher better | $\geq 1$ each |
| Final selection | On-target protein-DNA ipTM | higher better | $> 0.6$ |
| | NA-MPNN specificity margin | higher better | $> 0$ |
| | DeepPBS specificity margin | higher better | $> 0$ |
| | Designs selected per promoter | — | $\leq 14$ |

## References

J. Abramson, J. Adler, J. Dunger, R. Evans, T. Green, A. Pritzel, O. Ronneberger, L. Willmore, A. J. Ballard, J. Bambrick, S. W. Bodenstein, D. A. Evans, C.-C. Hung, M. O’Neill, D. Reiman, K. Tunyasuvunakool, Z. Wu, A. Žemgulytė, E. Arvaniti, C. Beattie, O. Bertolli, A. Bridgland, A. Cherepanov, M. Congreve, A. I. Cowen-Rivers, A. Cowie, M. Figurnov, F. B. Fuchs, H. Gladman, R. Jain, Y. A. Khan, C. M. R. Low, K. Perlin, A. Potapenko, P. Savy, S. Singh, A. Stecula, A. Thillaisundaram, C. Tong, S. Yakneen, E. D. Zhong, M. Zielinski, A. Žídek, V. Bapst, P. Kohli, M. Jaderberg, D. Hassabis, and J. M. Jumper. Accurate structure prediction of biomolecular interactions with AlphaFold 3. Nature, 630(8016):493–500, June 2024. ISSN 1476-4687. doi: 10.1038/s41586-024-07487-w.

Anthropic. Introducing Claude Opus 4.8. https://www.anthropic.com/news/claude-opus-4-8, May 2026.

F. Arnold. Design by directed evolution. Accounts of Chemical Research, 31(3):125–131, 1998. doi: 10.1021/ar960017f.

Ž. Avsec, V. Agarwal, D. Visentin, J. R. Ledsam, A. Grabska-Barwinska, K. R. Taylor, Y. Assael, J. Jumper, P. Kohli, and D. R. Kelley. Effective gene expression prediction from sequence by integrating long-range interactions. Nature Methods, 18(10):1196–1203, Oct. 2021. ISSN 1548-7105. doi: 10.1038/s41592-021-01252-x.

Ž. Avsec, N. Latysheva, J. Cheng, G. Novati, K. Taylor, T. Ward, C. Bycroft, L. Nicolaisen, E. Arvaniti, J. Pan, R. Thomas, V. Dutordoir, M. Perino, S. De, A. Karollus, A. Gayoso, T. Sargeant, A. Mottram, L. Wong, and P. Kohli. Advancing regulatory variant effect prediction with AlphaGenome. Nature, 649(8099):1206–1218, 2026. doi: 10.1038/s41586-025-10014-0.

Y. Barash, J. Calarco, W. Gao, Q. Pan, X. Wang, O. Shai, B. Blencowe, and B. Frey. Deciphering the splicing code. Nature, 465(7294):53–59, 2010. doi: 10.1038/nature09000.

S. Basu and B. Wallner. DockQ: A Quality Measure for Protein-Protein Docking Models. PLOS ONE, 11(8): e0161879, Aug. 2016. ISSN 1932-6203. doi: 10.1371/journal.pone.0161879.

S. Basu, Y. Gerchman, C. Collins, F. Arnold, and R. Weiss. A synthetic multicellular system for programmed pattern formation. Nature, 434(7037):1130–1134, 2005. doi: 10.1038/nature03461.

A. Bhatnagar, S. Jain, J. Beazer, S. C. Curran, A. M. Hoffnagle, K. S. Ching, M. Martyn, S. Nayfach, J. A. Ruffolo, and A. Madani. Scaling unlocks broader generation and deeper functional understanding of proteins. bioRxiv : the preprint server for biology, 2025. doi: 10.1101/2025.04.15.649055.

L. Bilitchenko, A. Liu, S. Cheung, E. Weeding, B. Xia, M. Leguia, J. Anderson, and D. Densmore. Eugene – a domain specific language for specifying and constraining synthetic biological parts, devices, and systems. PLOS ONE, 6(4):18882, 2011. doi: 10.1371/journal.pone.0018882.

E. Bingham, J. P. Chen, M. Jankowiak, F. Obermeyer, N. Pradhan, T. Karaletsos, R. Singh, P. Szerlip, P. Horsfall, and N. D. Goodman. Pyro: Deep Universal Probabilistic Programming. https://arxiv.org/abs/1810.09538, Oct. 2018.

C. Bland, T. L. Ramsey, F. Sabree, M. Lowe, K. Brown, N. C. Kyrpides, and P. Hugenholtz. CRISPR Recognition Tool (CRT): A tool for automatic detection of clustered regularly interspaced palindromic repeats. BMC Bioinformatics, 8(1):209, June 2007. ISSN 1471-2105. doi: 10.1186/1471-2105-8-209.

N. Boyd, S. Guns, and EscalanteBio. Mosaic. https://github.com/escalante-bio/mosaic, 2025.

G. Brixi, M. G. Durrant, J. Ku, M. Naghipourfar, M. Poli, G. Sun, G. Brockman, D. Chang, A. Fanton, G. A. Gonzalez, S. H. King, D. B. Li, A. T. Merchant, E. Nguyen, C. Ricci-Tam, D. W. Romero, J. C. Schmok, A. Taghibakhshi, A. Vorontsov, B. Yang, M. Deng, L. Gorton, N. Nguyen, N. K. Wang, M. T. Pearce, E. Simon, E. Adams, Z. J. Amador, E. A. Ashley, S. A. Baccus, H. Dai, S. Dillmann, S. Ermon, D. Guo, M. H. Herschl, R. Ilango, K. Janik, A. X. Lu, R. Mehta, M. R. K. Mofrad, M. Y. Ng, J. Pannu, C. Ré, J. St. John, J. Sullivan, J. Tey, B. Viggiano, K. Zhu, G. Zynda, D. Balsam, P. Collison, A. B. Costa, T. Hernandez-Boussard, E. Ho, M.-Y. Liu, T. McGrath, K. Powell, S. Pinglay, D. P. Burke, H. Goodarzi, P. D. Hsu, and B. L. Hie. Genome modelling and design across all domains of life with Evo 2. Nature, 652(8112):1349–1361, Apr. 2026. ISSN 1476-4687. doi: 10.1038/s41586-026-10176-5.

D. H. Brookes, H. Park, and J. Listgarten. Conditioning by adaptive sampling for robust design. https://arxiv.org/abs/1901.10060v9, Jan. 2019.

J. Brophy and C. Voigt. Principles of genetic circuit design. Nature Methods, 11(5):508–520, 2014. doi: 10.1038/nmeth.2926.

T. B. Brown, B. Mann, N. Ryder, M. Subbiah, J. Kaplan, P. Dhariwal, A. Neelakantan, P. Shyam, G. Sastry, A. Askell, S. Agarwal, A. Herbert-Voss, G. Krueger, T. Henighan, R. Child, A. Ramesh, D. M. Ziegler, J. Wu, C. Winter, C. Hesse, M. Chen, E. Sigler, M. Litwin, S. Gray, B. Chess, J. Clark, C. Berner, S. McCandlish, A. Radford, I. Sutskever, and D. Amodei. Language Models are Few-Shot Learners. https://arxiv.org/abs/2005.14165, July 2020.

J. Butcher, R. Krishna, R. Mitra, R. I. Brent, Y. Li, N. Corley, P. Kim, J. Funk, S. Mathis, S. Salike, A. Muraishi, H. Eisenach, T. R. Thompson, J. Chen, Y. Politanska, E. Sehgal, B. Coventry, O. Zhang, B. Qiang, K. Didi, M. Kazman, F. DiMaio, and D. Baker. De novo design of all-atom biomolecular interactions with rfdiffusion3. bioRxiv : the preprint server for biology, 2025. doi: 10.1101/2025.09.18.676967.

D. Cameron, C. Bashor, and J. Collins. A brief history of synthetic biology. Nature Reviews Microbiology, 12(5): 381–390, 2014. doi: 10.1038/nrmicro3239.

D. Cane, C. Walsh, and C. Khosla. Harnessing the biosynthetic code: Combinations, permutations, and mutations. Science, 282(5386):63–68, 1998. doi: 10.1126/science.282.5386.63.

B. Canton, A. Labno, and D. Endy. Refinement and standardization of synthetic biological parts and devices. Nature Biotechnology, 26(7):787–793, 2008. doi: 10.1038/nbt1413.

S. R. Carter, S. Curtis, C. Emerson, J. Gray, I. C. Haydon, A. Hebbeler, C. Qureshi, N. Randolph, A. Rives, and L. Stuart. Community values, guiding principles, and commitments for the responsible development of ai for protein design. https://responsiblebiodesign.ai/, Mar. 2023.

Chai Discovery, J. Boitreaud, J. Dent, M. McPartlon, J. Meier, V. Reis, A. Rogozhnikov, and K. Wu. Chai-1: Decoding the molecular interactions of life. bioRxiv : the preprint server for biology, 2024. doi: 10.1101/2024.10.10.615955.

S. Chaudhury, S. Lyskov, and J. J. Gray. PyRosetta: A script-based interface for implementing molecular modeling algorithms using Rosetta. Bioinformatics, 26(5):689–691, Mar. 2010. ISSN 1367-4803. doi: 10.1093/bioinformatics/btq007.

M. Chen, J. Tworek, H. Jun, Q. Yuan, H. P. d. O. Pinto, J. Kaplan, H. Edwards, Y. Burda, N. Joseph, G. Brockman, A. Ray, R. Puri, G. Krueger, M. Petrov, H. Khlaaf, G. Sastry, P. Mishkin, B. Chan, S. Gray, N. Ryder, M. Pavlov, A. Power, L. Kaiser, M. Bavarian, C. Winter, P. Tillet, F. P. Such, D. Cummings, M. Plappert, F. Chantzis, E. Barnes, A. Herbert-Voss, W. H. Guss, A. Nichol, A. Paino, N. Tezak, J. Tang, I. Babuschkin, S. Balaji, S. Jain, W. Saunders, C. Hesse, A. N. Carr, J. Leike, J. Achiam, V. Misra, E. Morikawa, A. Radford, M. Knight, M. Brundage, M. Murati, K. Mayer, P. Welinder, B. McGrew, D. Amodei, S. McCandlish, I. Sutskever, and W. Zaremba. Evaluating Large Language Models Trained on Code. https://arxiv.org/abs/2107.03374, July 2021.

X. D. Chen, M. Jim, M. Vallurupalli, K. Cao, A. N. Torres, J. W. Leong, Y. Zhang, D. Wollensak, Q. Gong, J. Sun, M. Borji, G. Schor, S. Mrowka, M. Hu, A. Laumas, J. A. Roth, T. Golub, and F. Chen. Generative design of cell type-specific RNA splicing elements for programmable gene regulation. bioRxiv : the preprint server for biology, 2025. doi: 10.1101/2025.11.05.686847.

Y. Cho, G. Rangel, G. Bhardwaj, and S. Ovchinnikov. Protein Hunter: Exploiting structure hallucination within diffusion for protein design. bioRxiv : the preprint server for biology, 2025. doi: 10.1101/2025.10.10.681530.

P. J. A. Cock, J. M. Chilton, B. Grüning, J. E. Johnson, and N. Soranzo. NCBI BLAST+ integrated into Galaxy. GigaScience, 4(1):s13742–015–0080–7, Dec. 2015. ISSN 2047-217X. doi: 10.1186/s13742-015-0080-7.

J. Dauparas, I. Anishchenko, N. Bennett, H. Bai, R. J. Ragotte, L. F. Milles, B. I. M. Wicky, A. Courbet, R. J. de Haas, N. Bethel, P. J. Y. Leung, T. F. Huddy, S. Pellock, D. Tischer, F. Chan, B. Koepnick, H. Nguyen, A. Kang, B. Sankaran, A. K. Bera, N. P. King, and D. Baker. Robust deep learning–based protein sequence design using ProteinMPNN. Science, 378(6615):49–56, Oct. 2022. doi: 10.1126/science.add2187.

J. Dauparas, G. R. Lee, R. Pecoraro, L. An, I. Anishchenko, C. Glasscock, and D. Baker. Atomic context-conditioned protein sequence design using LigandMPNN. Nature Methods, 22(4):717–723, Apr. 2025. ISSN 1548-7105. doi: 10.1038/s41592-025-02626-1.

F. A. Dreyer, D. Cutting, C. Schneider, H. Kenlay, and C. M. Deane. Inverse folding for antibody sequence design using deep learning. https://arxiv.org/abs/2310.19513, Oct. 2023.

Y. Du and L. Kaelbling. Compositional generative modeling: A single model is not all you need. https://arxiv.org/abs/2402.01103, June 2024.

V. Dubey and L. Shen. Personalized gene expression prediction in the era of deep learning: A review. Briefings in Bioinformatics, 27(1):022, 2026. doi: 10.1093/bib/bbag022.

K. Dudnyk, D. Cai, C. Shi, J. Xu, and J. Zhou. Sequence basis of transcription initiation in the human genome. Science, 384(6694):eadj0116, Apr. 2024. doi: 10.1126/science.adj0116.

M. Elowitz and S. Leibler. A synthetic oscillatory network of transcriptional regulators. Nature, 403(6767):335–338, 2000. doi: 10.1038/35002125.

D. Endy. Foundations for engineering biology. Nature, 438(7067):449–453, 2005. doi: 10.1038/nature04342.

R. Finn, J. Clements, and S. Eddy. HMMER web server: Interactive sequence similarity searching. Nucleic Acids Research, 39(suppl_2):29–37, 2011. doi: 10.1093/nar/gkr367.

M. Galdzicki, K. Clancy, E. Oberortner, M. Pocock, J. Quinn, C. Rodriguez, N. Roehner, M. Wilson, L. Adam, J. Anderson, B. Bartley, J. Beal, D. Chandran, J. Chen, D. Densmore, D. Endy, R. Grünberg, J. Hallinan, N. Hillson, and H. Sauro. The Synthetic Biology Open Language (SBOL) provides a community standard for communicating designs in synthetic biology. Nature Biotechnology, 32(6):545–550, 2014. doi: 10.1038/nbt.2891.

T. Gardner, C. Cantor, and J. Collins. Construction of a genetic toggle switch in Escherichia coli. Nature, 403 (6767):339–342, 2000. doi: 10.1038/35002131.

C. Glasscock, R. Pecoraro, R. McHugh, L. Doyle, W. Chen, O. Boivin, B. Lonnquist, E. Na, Y. Politanska, H. Haddox, D. Cox, C. Norn, B. Coventry, I. Goreshnik, D. Vafeados, G. Lee, R. Gordân, B. Stoddard, F. DiMaio, and D. Baker. Computational design of sequence-specific DNA-binding proteins. Nature Structural & Molecular Biology, 32 (11):2252–2261, 2025. doi: 10.1038/s41594-025-01669-4.

S. Gosai, R. Castro, N. Fuentes, J. Butts, K. Mouri, M. Alasoadura, S. Kales, T. Nguyen, R. Noche, A. Rao, M. Joy, P. Sabeti, S. Reilly, and R. Tewhey. Machine-guided design of cell-type-targeting cis-regulatory elements. Nature, 634(8036):1211–1220, 2024. doi: 10.1038/s41586-024-08070-z.

G. Gross, T. Waks, and Z. Eshhar. Expression of immunoglobulin-T-cell receptor chimeric molecules as functional receptors with antibody-type specificity. Proceedings of the National Academy of Sciences, 86(24):10024–10028, 1989. doi: 10.1073/pnas.86.24.10024.

W. Hastings. Monte Carlo sampling methods using Markov chains and their applications. Biometrika, 57(1): 97–109, 1970. doi: 10.1093/biomet/57.1.97.

T. Hayes, R. Rao, H. Akin, N. J. Sofroniew, D. Oktay, Z. Lin, R. Verkuil, V. Q. Tran, J. Deaton, M. Wiggert, R. Badkun-dri, I. Shafkat, J. Gong, A. Derry, R. S. Molina, N. Thomas, Y. A. Khan, C. Mishra, C. Kim, L. J. Bartie, M. Nemeth, P. D. Hsu, T. Sercu, S. Candido, and A. Rives. Simulating 500 million years of evolution with a language model. Science, 387(6736):850–858, Feb. 2025. doi: 10.1126/science.ads0018.

B. Hie, S. Candido, Z. Lin, O. Kabeli, R. Rao, N. Smetanin, T. Sercu, and A. Rives. A high-level programming language for generative protein design. bioRxiv : the preprint server for biology, 2022.doi: 10.1101/2022.12.21.521526.

B. Hie, V. Shanker, D. Xu, T. Bruun, P. Weidenbacher, S. Tang, W. Wu, J. Pak, and P. Kim. Efficient evolution of human antibodies from general protein language models. Nature Biotechnology, 42(2):275–283, 2024. doi: 10.1038/s41587-023-01763-2.

B. L. Hie and K. K. Yang. Adaptive machine learning for protein engineering. Current Opinion in Structural Biology, 72:145–152, Feb. 2022. ISSN 0959-440X. doi: 10.1016/j.sbi.2021.11.002.

G. Hinton. Training products of experts by minimizing contrastive divergence. Neural Computation, 14(8):1771–1800, 2002. doi: 10.1162/089976602760128018.

F. Hirsch, M. Varella-Garcia, P. Bunn, M. Maria, R. Veve, R. Bremmes, A. Barón, C. Zeng, and W. Franklin. Epidermal growth factor receptor in non-small-cell lung carcinomas: Correlation between gene copy number and protein expression and impact on prognosis. Journal of Clinical Oncology: Official Journal of the American Society of Clinical Oncology, 21(20):3798–3807, 2003. doi: 10.1200/JCO.2003.11.069.

T. Hopf, A. Gazizov, S. Busto, E. Eschbach, S. Lee, M. Mirdita, R. Orenbuch, K. Belahsen, D. Ross, C. Sander, M. Steinegger, S. d’Oelsnitz, and D. Marks. In Evedesign: Accessible Biosequence Design with a Unified Framework, page 2026 03 17 712115. 2026. doi: 10.64898/2026.03.17.712115.

P.-S. Huang, S. Boyken, and D. Baker. The coming of age of de novo protein design. Nature, 537(7620):320–327, 2016. doi: 10.1038/nature19946.

D. Hyatt, G.-L. Chen, P. F. LoCascio, M. L. Land, F. W. Larimer, and L. J. Hauser. Prodigal: Prokaryotic gene recognition and translation initiation site identification. BMC Bioinformatics, 11(1):119, Mar. 2010. ISSN 1471-2105. doi: 10.1186/1471-2105-11-119.

J. Ingraham, M. Baranov, Z. Costello, K. Barber, W. Wang, A. Ismail, V. Frappier, D. Lord, C. Ng-Thow-Hing, E. Vlack, S. Tie, V. Xue, S. Cowles, A. Leung, J. Rodrigues, C. Morales-Perez, A. Ayoub, R. Green, K. Puentes, and G. Grigoryan. Illuminating protein space with a programmable generative model. Nature, 623(7989): 1070–1078, 2023. doi: 10.1038/s41586-023-06728-8.

Z. Ji, N. Lee, R. Frieske, T. Yu, D. Su, Y. Xu, E. Ishii, Y. Bang, D. Chen, W. Dai, H. Chan, A. Madotto, and P. Fung. Survey of hallucination in natural language generation. ACM Computing Surveys, 55(12):1–38, 2023. doi: 10.1145/3571730.

M. Jinek, K. Chylinski, I. Fonfara, M. Hauer, J. Doudna, and E. Charpentier. A programmable dual-RNA–guided DNA endonuclease in adaptive bacterial immunity. Science, 337(6096):816–821, 2012. doi: 10.1126/science. 1225829.

B. John, A. J. Enright, A. Aravin, T. Tuschl, C. Sander, and D. S. Marks. Human MicroRNA Targets. PLOS Biology, 2(11):e363, Oct. 2004. ISSN 1545-7885. doi: 10.1371/journal.pbio.0020363.

T. Jones, S. Oliveira, C. Myers, C. Voigt, and D. Densmore. Genetic circuit design automation with Cello 2.0. Nature Protocols, 17(4):1097–1113, 2022. doi: 10.1038/s41596-021-00675-2.

J. Jumper, R. Evans, A. Pritzel, T. Green, M. Figurnov, O. Ronneberger, K. Tunyasuvunakool, R. Bates, A. Žídek, A. Potapenko, A. Bridgland, C. Meyer, S. A. A. Kohl, A. J. Ballard, A. Cowie, B. Romera-Paredes, S. Nikolov, R. Jain, J. Adler, T. Back, S. Petersen, D. Reiman, E. Clancy, M. Zielinski, M. Steinegger, M. Pacholska, T. Berghammer, S. Bodenstein, D. Silver, O. Vinyals, A. W. Senior, K. Kavukcuoglu, P. Kohli, and D. Hassabis. Highly accurate protein structure prediction with AlphaFold. Nature, 596(7873):583–589, Aug. 2021. ISSN 1476-4687. doi: 10.1038/s41586-021-03819-2.

A. Karpathy. There’s a new kind of coding i call “vibe coding”. X post, Feb. 2025. https://x.com/karpathy/status/1886192184808149383.

K. Katoh, K. Misawa, K.-i. Kuma, and T. Miyata. MAFFT: A novel method for rapid multiple sequence alignment based on fast Fourier transform. Nucleic Acids Research, 30(14):3059–3066, July 2002. ISSN 0305-1048.

N. Kim, G. Carluccio, K. Zhang, and J. Collins. Generative AI for synthetic biology: Designing biological parts, circuits, and genomes. Cell Systems, 17(2), 2026. doi: 10.1016/j.cels.2026.101533.

S. H. King, C. L. Driscoll, D. B. Li, D. Guo, A. T. Merchant, G. Brixi, M. E. Wilkinson, and B. L. Hie. Generative design of novel bacteriophages with genome language models. bioRxiv : the preprint server for biology, 2025. doi: 10.1101/2025.09.12.675911.

S. Kirkpatrick, C. Gelatt, and M. Vecchi. Optimization by simulated annealing. Science, 220(4598):671–680, 1983. doi: 10.1126/science.220.4598.671.

S. Kosuri, D. Goodman, G. Cambray, V. Mutalik, Y. Gao, A. Arkin, D. Endy, and G. Church. Composability of regulatory sequences controlling transcription and translation in Escherichia coli. Proceedings of the National Academy of Sciences, 110(34):14024–14029, 2013. doi: 10.1073/pnas.1301301110.

F. Kschischang, B. Frey, and H.-A. Loeliger. Factor graphs and the sum-product algorithm. IEEE Transactions on Information Theory, 47(2):498–519, 2001. doi: 10.1109/18.910572.

A. Kubaney, A. Favor, L. McHugh, R. Mitra, R. Pecoraro, J. Dauparas, C. Glasscock, and D. Baker. RNA sequence design and protein–DNA specificity prediction with NA-MPNN. bioRxiv : the preprint server for biology, 2025. doi: 10.1101/2025.10.03.679414.

T. LaFleur, A. Hossain, and H. Salis. Automated model-predictive design of synthetic promoters to control transcriptional profiles in bacteria. Nature Communications, 13(1):5159, 2022. doi: 10.1038/s41467-022-32829-5.

M. Larralde and G. Zeller. PyHMMER: A Python library binding to HMMER for efficient sequence analysis. Bioinformatics, 39(5):btad214, May 2023. ISSN 1367-4811. doi: 10.1093/bioinformatics/btad214.

Y. LeCun, S. Chopra, R. Hadsell, M. Ranzato, and F. J. Huang. A tutorial on energy-based learning. In G. Bakir, T. Hofmann, B. Schölkopf, A. J. Smola, and B. Taskar, editors, Predicting Structured Data. MIT Press, 2006. Version 1.0, August 19, 2006.

M. Lewis, G. Chang, N. Horton, M. Kercher, H. Pace, M. Schumacher, R. Brennan, and P. Lu. Crystal structure of the lactose operon repressor and its complexes with DNA and inducer. Science, 271(5253):1247–1254, 1996. doi: 10.1126/science.271.5253.1247.

S. Lewis, T. Hempel, J. Jiménez-Luna, M. Gastegger, Y. Xie, A. Y. K. Foong, V. G. Satorras, O. Abdin, B. S. Veeling, I. Zaporozhets, Y. Chen, S. Yang, A. E. Foster, A. Schneuing, J. Nigam, F. Barbero, V. Stimper, A. Campbell, J. Yim, M. Lienen, Y. Shi, S. Zheng, H. Schulz, U. Munir, R. Sordillo, R. Tomioka, C. Clementi, and F. Noé. Scalable emulation of protein equilibrium ensembles with generative deep learning. Science, 389(6761):eadv9817, July 2025. doi: 10.1126/science.adv9817.

Z. Lin, H. Akin, R. Rao, B. Hie, Z. Zhu, W. Lu, N. Smetanin, R. Verkuil, O. Kabeli, Y. Shmueli, A. dos Santos Costa, M. Fazel-Zarandi, T. Sercu, S. Candido, and A. Rives. Evolutionary-scale prediction of atomic-level protein structure with a language model. Science, 379(6637):1123–1130, Mar. 2023. doi: 10.1126/science.ade2574.

J. Linder, D. Srivastava, H. Yuan, V. Agarwal, and D. Kelley. Predicting RNA-seq coverage from DNA sequence as a unifying model of gene regulation. Nature Genetics, 57(4):949–961, 2025. doi: 10.1038/s41588-024-02053-6.

J. Ling, A. Bygrave, C. Santiago, R. Carmen-Orozco, V. Trinh, M. Yu, Y. Li, Y. Liu, K. Bowden, L. Duncan, J. Han, K. Taneja, R. Dongmo, T. Babola, P. Parker, L. Jiang, P. Leavey, J. Smith, R. Vistein, and S. Blackshaw. Cell-specific regulation of gene expression using splicing-dependent frameshifting. Nature Communications, 13(1): 5773, 2022. doi: 10.1038/s41467-022-33523-2.

J. Listgarten and H. Jiang. How artificial intelligence is reengineering protein engineering. Science, 392(6794): 159–166, Apr. 2026. doi: 10.1126/science.aec8444.

X.-J. Lu and W. Olson. 3DNA: A versatile, integrated software system for the analysis, rebuilding and visualization of three-dimensional nucleic-acid structures. Nature Protocols, 3(7):1213–1227, 2008. doi: 10.1038/nprot.2008.104.

L. Luebbert. Paving the way for agents in biology. https://www.anthropic.com/research/agents-in-biology, June 2026.

A. Madani, B. Krause, E. Greene, S. Subramanian, B. Mohr, J. Holton, J. Olmos, C. Xiong, Z. Sun, R. Socher, J. Fraser, and N. Naik. Large language models generate functional protein sequences across diverse families. Nature Biotechnology, 41(8):1099–1106, 2023. doi: 10.1038/s41587-022-01618-2.

J. Maloney, M. Resnick, N. Rusk, B. Silverman, and E. Eastmond. The Scratch Programming Language and Environment. ACM Transactions on Computing Education (TOCE), 10(4):16:1–16:15, Nov. 2010. doi: 10.1145/ 1868358.1868363.

S. E. McGeary, K. S. Lin, C. Y. Shi, T. M. Pham, N. Bisaria, G. M. Kelley, and D. P. Bartel. The biochemical basis of microRNA targeting efficacy. Science, 366(6472):eaav1741, Dec. 2019. ISSN 1095-9203. doi:10.1126/ science.aav1741.

A. Merchant, S. King, E. Nguyen, and B. Hie. Semantic design of functional de novo genes from a genomic language model. Nature, 1–10, 2025. doi: 10.1038/s41586-025-09749-7.

N. Metropolis, A. Rosenbluth, M. Rosenbluth, A. Teller, and E. Teller. Equation of state calculations by fast computing machines. The Journal of Chemical Physics, 21(6):1087–1092, 1953. doi: 10.1063/1.1699114.

L. S. Mille-Fragoso, J. N. Wang, C. L. Driscoll, H. Dai, T. Widatalla, X. Zhang, B. L. Hie, and X. J. Gao. Efficient generation of epitope-targeted de novo antibodies with Germinal. bioRxiv : the preprint server for biology, 2025. doi: 10.1101/2025.09.19.677421.

R. Mitra, J. Li, J. M. Sagendorf, Y. Jiang, A. S. Cohen, T.-P. Chiu, C. J. Glasscock, and R. Rohs. Geometric deep learning of protein–DNA binding specificity. Nature Methods, 21(9):1674–1683, Sept. 2024. ISSN 1548-7105. doi: 10.1038/s41592-024-02372-w.

A. Mitrofanov, M. Ziemann, O. S. Alkhnbashi, W. R. Hess, and R. Backofen. CRISPRtracrRNA: Robust approach for CRISPR tracrRNA detection. Bioinformatics, 38(Supplement_2):ii42–ii48, Sept. 2022. ISSN 1367-4803. doi: 10.1093/bioinformatics/btac466.

F. Moolten. Tumor chemosensitivity conferred by inserted herpes thymidine kinase genes: Paradigm for a prospective cancer control strategy. Cancer Research, 46(10):5276–5281, 1986.

T. Moon, C. Lou, A. Tamsir, B. Stanton, and C. Voigt. Genetic programs constructed from layered logic gates in single cells. Nature, 491(7423):249–253, 2012. doi: 10.1038/nature11516.

L. Morsut, K. Roybal, X. Xiong, R. Gordley, S. Coyle, M. Thomson, and W. Lim. Engineering customized cell sensing and response behaviors using synthetic notch receptors. Cell, 164(4):780–791, 2016. doi: 10.1016/j.cell.2016.01.012.

E. Nguyen, M. Poli, M. Durrant, B. Kang, D. Katrekar, D. Li, L. Bartie, A. Thomas, S. King, G. Brixi, J. Sullivan, M. Ng, A. Lewis, A. Lou, S. Ermon, S. Baccus, T. Hernandez-Boussard, C. Ré, P. Hsu, and B. Hie. Sequence modeling and design from molecular to genome scale with Evo. Science, 386(6723):9336, 2024. doi:10.1126/ science.ado9336.

A. Nielsen, B. Der, J. Shin, P. Vaidyanathan, V. Paralanov, E. Strychalski, D. Ross, D. Densmore, and C. Voigt. Genetic circuit design automation. Science, 352(6281):7341, 2016. doi: 10.1126/science.aac7341.

B. Nieuwenhuis, B. Haenzi, S. Hilton, A. Carnicer-Lombarte, B. Hobo, J. Verhaagen, and J. W. Fawcett. Optimization of adeno-associated viral vector-mediated transduction of the corticospinal tract: Comparison of four promoters. Gene Therapy, 28(1-2):56–74, Feb. 2021. ISSN 1476-5462. doi: 10.1038/s41434-020-0169-1.

T. H. Olsen, I. H. Moal, and C. M. Deane. AbLang: An antibody language model for completing antibody sequences. Bioinformatics Advances, 2(1):vbac046, Jan. 2022. ISSN 2635-0041. doi: 10.1093/bioadv/vbac046. OpenAI. GPT-5.5 System Card. https://deploymentsafety.openai.com/gpt-5-5, May 2026.

S. Ovchinnikov, S. Feng, J. Dauparas, W. Wu, and C. Frank. Sokrypton/ColabDesign. https://github.com/sokrypton/ColabDesign, 2025.

M. Pacesa, L. Nickel, C. Schellhaas, J. Schmidt, E. Pyatova, L. Kissling, P. Barendse, J. Choudhury, S. Kapoor, A. Alcaraz-Serna, Y. Cho, K. Ghamary, L. Vinué, B. Yachnin, A. Wollacott, S. Buckley, A. Westphal, S. Lindhoud, S. Georgeon, and B. Correia. One-shot design of functional protein binders with BindCraft. Nature, 646(8084): 483–492, 2025. doi: 10.1038/s41586-025-09429-6.

S. Park, S. Myung, and M. Baek. Advancing protein structure prediction beyond AlphaFold2. Current Opinion in Structural Biology, 90:102985, 2025. doi: 10.1016/j.sbi.2025.102985.

S. Passaro, G. Corso, J. Wohlwend, M. Reveiz, S. Thaler, V. R. Somnath, N. Getz, T. Portnoi, J. Roy, H. Stark, D. Kwabi-Addo, D. Beaini, T. Jaakkola, and R. Barzilay. Boltz-2: Towards accurate and efficient binding affinity prediction. bioRxiv : the preprint server for biology, 2025. doi: 10.1101/2025.06.14.659707.

N. Robinson, W. Zhang, R. Ghosh, B. Gerber, H. Zhang, C. Sanfiorenzo, S. Wang, D. Carlo, and K. Wang. Construction of complex and diverse DNA sequences using DNA three-way junctions. Nature, 651(8105):491–500, 2026. doi: 10.1038/s41586-025-10006-0.

J. Roney and S. Ovchinnikov. State-of-the-art estimation of protein model accuracy using AlphaFold. Physical Review Letters, 129(23):238101, 2022. doi: 10.1103/PhysRevLett.129.238101.

J. Ruffolo, S. Nayfach, J. Gallagher, A. Bhatnagar, J. Beazer, R. Hussain, J. Russ, J. Yip, E. Hill, M. Pacesa, A. Meeske, P. Cameron, and A. Madani. Design of highly functional genome editors by modelling CRISPR–Cas sequences. Nature, 645(8080):518–525, 2025. doi: 10.1038/s41586-025-09298-z.

H. Salgado, S. Gama-Castro, P. Lara, C. Mejia-Almonte, G. Alarcón-Carranza, A. G. López-Almazo, F. Betancourt-Figueroa, P. Peña-Loredo, S. Alquicira-Hernández, D. Ledezma-Tejeida, L. Arizmendi-Zagal, F. Mendez-Hernandez, A. K. Diaz-Gomez, E. Ochoa-Praxedis, L. J. Muñiz-Rascado, J. S. García-Sotelo, F. A. Flores-Gallegos, L. Gómez, C. Bonavides-Martínez, V. M. del Moral-Chávez, A. J. Hernández-Alvarez, A. Santos-Zavaleta, S. Capella-Gutierrez, J. L. Gelpi, and J. Collado-Vides. RegulonDB v12.0: A comprehensive resource of transcriptional regulation in E. coli K-12. Nucleic Acids Research, 52(D1):D255–D264, Nov. 2023. ISSN 0305-1048. doi: 10.1093/nar/gkad1072.

T. Schick, J. Dwivedi-Yu, R. Dessì, R. Raileanu, M. Lomeli, L. Zettlemoyer, N. Cancedda, and T. Scialom. Toolformer: Language models can teach themselves. https://arxiv.org/abs/2302.04761, 2023.

Schrödinger, LLC. The AxPyMOL Molecular Graphics Plugin for Microsoft PowerPoint, Version 1.8. Nov. 2015.

E. Sehgal, Y. Politanska, R. Mitra, P. T. Kim, N. G. Rodríguez, T. Warrier, A. Kubaney, A. Morishita, R. Quijano, J. Butcher, R. Krishna, R. J. Pecoraro, B. Belmont, N. Roullier, I. Goreshnik, D. K. Vafeados, P. Kwon, R. Ramarao, J. Taipale, C. J. Glasscock, and D. Baker. Generative design of sequence specific DNA binding proteins. bioRxiv : the preprint server for biology, 2026. doi: 10.64898/2026.04.27.720408.

V. Shanker, T. Bruun, B. Hie, and P. Kim. Unsupervised evolution of protein and antibody complexes with a structure-informed language model. Science, 385(6704):46–53, 2024. doi: 10.1126/science.adk8946.

S. Sharma, D. Bell, J. Settleman, and D. Haber. Epidermal growth factor receptor mutations in lung cancer. Nature Reviews. Cancer, 7(3):169–181, 2007. doi: 10.1038/nrc2088.

R. Shetty, D. Endy, and T. Knight. Engineering BioBrick vectors from BioBrick parts. Journal of Biological Engineering, 2:5, 2008. doi: 10.1186/1754-1611-2-5.

U. Singh and E. S. Wurtele. Orfipy: A fast and flexible tool for extracting ORFs. Bioinformatics, 37(18):3019–3020, Sept. 2021. ISSN 1367-4803. doi: 10.1093/bioinformatics/btab090.

M. Steinegger and J. Söding. MMseqs2 enables sensitive protein sequence searching for the analysis of massive data sets. Nature Biotechnology, 35(11):1026–1028, Nov. 2017. ISSN 1546-1696. doi: 10.1038/nbt.3988.

B. Teague. Cytoflow: A python toolbox for flow cytometry. bioRxiv : the preprint server for biology, 2022.doi: 10.1101/2022.07.22.501078.

D. Tran, M. D. Hoffman, R. A. Saurous, E. Brevdo, K. Murphy, and D. M. Blei. Deep Probabilistic Programming. https://arxiv.org/abs/1701.03757, Mar. 2017.

J.-W. van de Meent, B. Paige, H. Yang, and F. Wood. An Introduction to Probabilistic Programming. https://arxiv.org/abs/1809.10756, Oct. 2021.

C. Voigt, C. Martinez, Z.-G. Wang, S. Mayo, and F. Arnold. Protein building blocks preserved by recombination. Nature Structural Biology, 9(7):553–558, 2002. doi: 10.1038/nsb805.

E. Wang, R. Sandberg, S. Luo, I. Khrebtukova, L. Zhang, C. Mayr, S. Kingsmore, G. Schroth, and C. Burge. Alternative isoform regulation in human tissue transcriptomes. Nature, 456(7221):470–476, 2008. doi: 10.1038/nature07509.

H. H. Wang, F. J. Isaacs, P. A. Carr, Z. Z. Sun, G. Xu, C. R. Forest, and G. M. Church. Programming cells by multiplex genome engineering and accelerated evolution. Nature, 460(7257):894–898, Aug. 2009. ISSN 1476-4687. doi: 10.1038/nature08187.

J. Wang and J. Doudna. CRISPR technology: A decade of genome editing is only the beginning. Science, 379 (6629):8643, 2023. doi: 10.1126/science.add8643.

S. K. Wang, B. Deng, S. Nair, X. Ren, J. Li, J. Tijerina, P. Prakhar, Z. Luo, C. Nnebe, S. H. Kim, Y. Zhou, S. H. Shah, A. Davis, R. Mahajan, Y. Qiao, Y. Zhou, J. Zhang, Y. Xue, J. L. Goldberg, W. Wei, A. Kundaje, H. Y. Chang, and S. Wang. Deep learning-guided design of cell type-specific AAV promoters. bioRxiv : the preprint server for biology, 2026. doi: 10.64898/2026.01.13.699371.

J. Watson, D. Juergens, N. Bennett, B. Trippe, J. Yim, H. Eisenach, W. Ahern, A. Borst, R. Ragotte, L. Milles, B. Wicky, N. Hanikel, S. Pellock, A. Courbet, W. Sheffler, J. Wang, P. Venkatesh, I. Sappington, S. Torres, and D. Baker. De novo design of protein structure and function with RFdiffusion. Nature, 620(7976):1089–1100, 2023. doi: 10.1038/s41586-023-06415-8.

J. Wei, X. Wang, D. Schuurmans, M. Bosma, E. H. Chi, Q. Le, and D. Zhou. Chain of thought prompting elicits reasoning in large language models. CoRR, abs/2201.11903, 2022.

N. You, C. Liu, Y. Gu, R. Wang, H. Jia, T. Zhang, S. Jiang, J. Shi, M. Chen, M.-X. Guan, S. Sun, S. Pei, Z. Liu, and N. Shen. SpliceTransformer predicts tissue-specific splicing linked to human diseases. Nature Communications, 15(1):9129, Oct. 2024. ISSN 2041-1723. doi: 10.1038/s41467-024-53088-6.

C. Zhang, M. Shine, A. Pyle, and Y. Zhang. US-align: Universal structure alignments of proteins, nucleic acids, and macromolecular complexes. Nature Methods, 19(9):1109–1115, 2022. doi: 10.1038/s41592-022-01585-1.

